# T cell infiltration stratifies mismatch repair-proficient colorectal cancer independently of tumor genotype

**DOI:** 10.1101/2024.09.24.614701

**Authors:** Jeremiah Wala, Kristin Qian, Yi Daniel Lu, Clarence Yapp, Ino de Bruijn, Shannon Coy, Andréanne Gagné, Ben Gaudio, Sabrina Chan, Jia-Ren Lin, Yu-An Chen, John Hoffer, Jeremy Muhlich, Nikolaus Schultz, Sandro Santagata, Peter K Sorger

## Abstract

Cancer subtyping increasingly relies on both tumor mutational status and the spatial organization of the tumor microenvironment (TME). In colorectal cancer (CRC), mismatch repair deficient tumors (dMMR) are generally more immune infiltrated than mismatch repair proficient (pMMR) tumors, have better prognosis following resection, and respond to immune checkpoint inhibitors. Whether some pMMR tumors also harbor tumor-engaged T cells is unknown. Integrating spatial profiling with genomic analysis, we find that ∼20% of pMMR tumors exhibit similar levels of T cell infiltration as dMMR tumors but without any specific somatic driver mutation or higher tumor mutational burden. These T cell-infiltrated pMMR (tipMMR) tumors contain abundant cytotoxic PD1^+^ Granzyme B^+^ CD8^+^ T cells in direct contact with tumor epithelium and immune clusters enriched for TCF1^+^ precursor exhausted T cells. tipMMR CRC tumors are similar in frequency as dMMR tumors and their immunologically active state suggests that immunotherapy should be considered in this population.

## INTRODUCTION

In colorectal cancer (CRC), the strongest link between mutational status and the TME involves a lymphocytic infiltrate specifically in tumors that are mismatch repair deficient (dMMR) or microsatellite instability-high (MSI-H). These two phenotypes are assessed differently (immunohistochemistry for dMMR detection and sequencing or PCR for MSI-H detection) but are highly concordant and account for ∼15% of CRC cases. dMMR tumors are characterized by high tumor mutational burden (TMB), high levels of T cells in the tumor core^1^, and sensitivity to PD1/PDL1-directed immune checkpoint inhibitors (ICIs)^2,3^. A high level of T cell infiltration and ICI-responsiveness is also observed in the 1-2% of CRCs that harbor polymerase epsilon (*POLE*) mutations. In contrast, most CRCs are mismatch repair-proficient or microsatellite stable (pMMR/MSS) and have a low TMB. These pMMR CRCs are substantially less infiltrated with lymphocytes than dMMR tumors and are currently treated with chemotherapy, not immunotherapy.

While there is currently no established role for ICIs in pMMR CRC (apart from *POLE* mutated tumors), attempts are underway to identify disease settings or tumor characteristics that may enrich for a minority population of responders. For example, the NICHE 2 trial reported a pathological response rate of 26% in previously untreated early-stage pMMR colon cancer following a single dose of ipilimumab and two doses of nivolumab (which block the CTLA4 and PD1 checkpoint receptors, respectively)^4^. Reports of ICI responsiveness in at least some pMMR CRCs, the high prevalence of pMMR relative to dMMR CRC, and the limited treatment options for pMMR CRC apart from cytotoxic chemotherapy, all argue for additional analysis of ICIs and other immunotherapies in pMMR CRC.

The idea that pMMR CRC has a universally “cold” TME is increasingly challenged by immunoprofiling data. When the numbers of lymphocytes expressing the T cell markers CD3 and CD8 in the tumor compartment of localized CRC are measured using immunohistochemistry (IHC) via the Immunoscore assay^5–7^, infiltration is strongly prognostic independent of MMR status. The Immunoscore is included in European treatment guidelines as a means of refining prognosis in conjunction with TNM staging^8^. A related approach, ImmunoProfile, which measures CD8, PD1, FOXP3, and PDL1 in a single multiplexed immunofluorescence (IF) test, has also demonstrated that the degree of intratumoral CD8^+^ or PD1^+^ T cells is associated with significantly greater survival in CRC and other solid tumor types.^9^

In this study, we integrate spatial immunoprofiling with clinical annotation, MMR status, targeted somatic sequencing, and deep T cell-subtype profiling via cyclic immunofluorescence imaging (CyCIF) to enable a systematic analysis of TME features in a cohort of primarily untreated CRC spanning MMR types. We define a relatively common subset of highly T cell-infiltrated pMMR (tipMMR) tumors that exhibit the immune invasion markers and extended survival characteristic of dMMR tumors. tipMMR tumors show high infiltration of cytotoxic CD8^+^ T cells and elevated expression of the immune checkpoint proteins PD1 and PDL1 but are otherwise indistinguishable from T cell-depleted MMR-proficient (tdpMMR) tumors in grade, tumor mutational burden (TMB), *POLE* mutations, somatic copy-number burden or the presence of other previously identified CRC driver mutations. tipMMR tumors differ from dMMR tumors, however, in having a higher abundance of FOXP3^+^ regulatory T cells (Tregs). We propose that tipMMR defines a population poised for immunotherapy but potentially requiring additional agents directed at these suppressive programs.

## RESULTS

### Spatial profiling identifies a population of highly T cell-infiltrated pMMR tumors

We curated clinical records for a previously described 74-patient CRC cohort^10^ (**Figure 1A; Supplementary Table 1**) and abstracted pathology reports for features such as lymphovascular invasion (LVI), perineural invasion (PNI), and histological grade. For this study we defined progression-free survival (PFS) as the time to radiographic progression following resection. Median follow-up was 64.6 months. All tumors were primary resections, and most patients were treatment-naïve (66/74), although a small number (8/74) received neoadjuvant chemotherapy and had residual tumor (pathological stage ypT3 or greater). MMR status was established by IHC by clinical pathology and when some ambiguity was identified on re-review, additional IHC was performed to confirm the status of each tumor. We also reviewed OncoPanel^11^ clinical targeted sequencing data (which covers 275 to 447 cancer genes) for *MLH1*, *MSH2*, *MSH6*, and *PMS2* alterations. For all but one patient, we confirmed that the MMR status determined by OncoPanel sequencing matched the IHC results. One patient exhibited intact nuclear staining for MMR proteins but had microsatellite instability by sequencing and was classified as dMMR in our analysis. Overall, this yielded 10 dMMR or MSI-H and 64 pMMR or MSS tumors. Consistent with data from other cohorts^12^, PFS was also longer for dMMR than pMMR tumors in our cohort (TNM stage-adjusted HR = 0.17, 95% CI 0.02-1.26; **Extended Figure 1A**).

**Figure 1:**
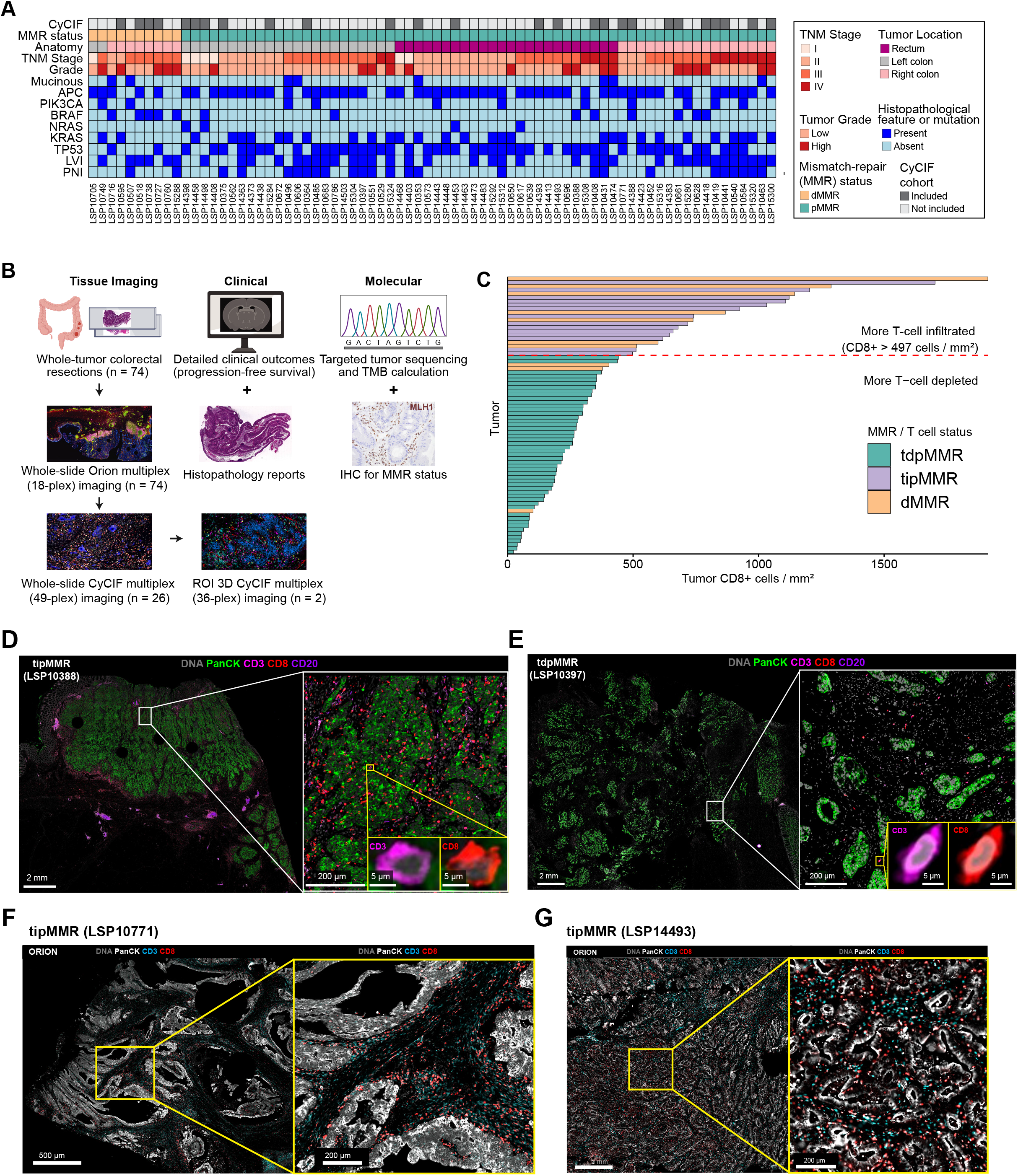
Features of the cohort. Colorectal cancers (CRC) subjected to spatial profiling using Orion multiplex immunofluorescence imaging were analyzed for clinical, histopathology and molecular features. (A) Cohort plot of 74 patients showing inclusion in the CyCIF cohort (n = 26), MMR status, anatomic location, TNM stage, histological grade, mucinous histology, and status of multiple binary genomic and histopathological features (LVI = lymphovascular invasion; PNI = perineural invasion). (B) Overview of the multimodal dataset. (C) Waterfall plot of all 74 tumors ordered (y-axis) by CD8^+^ cell density in the tumor compartment (x-axis; cells per mm^2^). Bars are colored by MMR and T cell status: pMMR tumors above the dashed line (CD8^+^ density > 497 cells/mm^2^) are designated T cell-infiltrated pMMR (tipMMR, n = 14) and those below it T cell-depleted pMMR (tdpMMR, n = 50). (D) Orion multiplex image of a tipMMR tumor showing dense tumor cells and a high degree of intermixed CD3^+^ and CD8^+^ T cells. Three different magnifications shown, with highest (lower right) depicting single cells. (E) T cell-depleted pMMR (tdpMMR) tumor with intermixed stroma comprised of relatively few T cells. (F) Orion multiplex IF images of a peritumoral lymphocytic reaction in two tipMMR tumors.

The overall pattern of mutations in our specimens was characteristic of CRC more broadly, including enrichment of *BRAF* alterations in right-sided and dMMR tumors (p < 0.05; Fisher’s exact test; **Extended Figure 1B**). Among mutations, *KRAS* mutations were the strongest negative predictor of PFS (TNM stage adjusted HR 1.63, 95% CI 0.83-3.20, p = 0.16, Wald test), as has been previously reported^13^. We also recapitulated the finding of worse outcomes for tumors positive for LVI and PNI (**Extended Figures 1C-D**)^8^. We conclude that our cohort reflects the well-established clinical, histopathology, and molecular features of the general CRC population, making it reasonable to compare the prognostic power of these features with the imaging-derived features described below.

To quantify the spatial immune landscape of primary CRC tumors, we combined 18-plex Orion images with same-section H&E imaging (**Figure 1B**). Individual cell types were identified hierarchically, similarly to flow cytometry, by setting marker thresholds through combined computational analysis and expert review. This yielded 9.6 x 10^7^ segmented cells, identified as CD3^+^ T cells (including CD8^+^ cytotoxic T cells and CD4^+^ helper T cells), CD20^+^ B cells, CD163^+^ and CD68^+^ monocytes (macrophages and dendritic cells), as well as vessels and supporting stroma (based on CD31 and smooth muscle actin αSMA staining; **Extended Figure 1E, F**). The tumor region was identified by having a pathologist annotate same-section H&E and multiplexed immunofluorescence (mIF) Orion images. Immune cell infiltration into the tumor and surrounding non-tumor stroma was quantified as the mean density of each cell type (in cells/mm^2^).

Across all specimens we observed that CD8^+^ T cell infiltration into the tumor domain varied over a 90-fold range, from 15 cells/mm^2^ in the least infiltrated (“coldest”) tumor to 1,380 cells/mm^2^ in the most infiltrated (“hottest”) tumor. Overall, dMMR tumors were significantly more infiltrated than pMMR tumors (median CD8^+^ cells/mm^2^: pMMR=187, dMMR=520; p = 0.002). Unexpectedly, the most highly infiltrated specimens included not only dMMR tumors, but a similar number of pMMR tumors. We classified tumors as T cell-infiltrated MMR-proficient (tipMMR) when CD8^+^ T cell density was greater than 497 cells/mm^2^, a cutoff that captured 70% (7/10) of all dMMR tumors. This resulted in 22% (14/64) of pMMR tumors classified as tipMMR (**Figure 1C**). The remaining pMMR tumors were defined as T cell-depleted MMR-proficient (tdpMMR). We then used these labels to compare genomic, clinical, and spatial features.

While each tipMMR tumor, by definition, exhibited higher than average T cell infiltration, inspection of multiplexed images revealed heterogeneity in the spatial organization of tumor-immune interactions. Several tipMMR tumors showed significant intermixing of tumor cells and T cells, resembling an inflamed phenotype (**Figure 1D**) that differed from the cold TME of tdpMMR tumors (**Figure 1E**). Other tipMMR tumors exhibited dense bands of T cells surrounding tumor nests, a pattern previously described in CRCs that corresponds to a lymphocytic reaction (**Figure 1F-G**)^14^. Qualitatively, the tumor compartments of both tdpMMR and tipMMR samples showed similar variation in morphology, ranging from dense sheets of high-grade tumor cells to more organized PanCK^+^ tumor nests surrounded by bands of PanCK^NEG^ stromal cells. The origins and significance of these differences are as yet unknown, but we found no significant difference in cellular grade between tipMMR and tdpMMR tumors, as assessed by traditional clinical pathology review (p = 0.49; Fisher’s exact). Additionally, when we assessed the relative proportions of stromal, epithelial, and immune cell markers among dMMR, tdpMMR and tipMMR, we observed no significant differences in expression of PanCK, E-cadherin (ECAD) or αSMA, or the density of proliferating Ki67^+^ tumor cells. Thus, while tipMMR tumors represent a distinct CRC state they share a similar epithelial and stromal landscape with other CRCs.

### tipMMR status is independent of tumor genotype and is associated with favorable outcomes

TMB correlates with neoantigen load, which is an established mediator of T cell infiltration and activation^15^. As expected, TMB values computed from OncoPanel data for dMMR tumors were a significantly higher than for either tipMMR or tdpMMR tumors (p < 4 x 10^-4^; unless otherwise stated, group comparisons were made with two-sided Wilcoxon rank-sum tests; **Figure 2A**). In contrast, we did not observe any significant difference between tipMMR and tdpMMR tumors (p = 0.61), suggesting that TMB was not a primary determinant of greater immune infiltration in tipMMR tumors.

**Figure 2:**
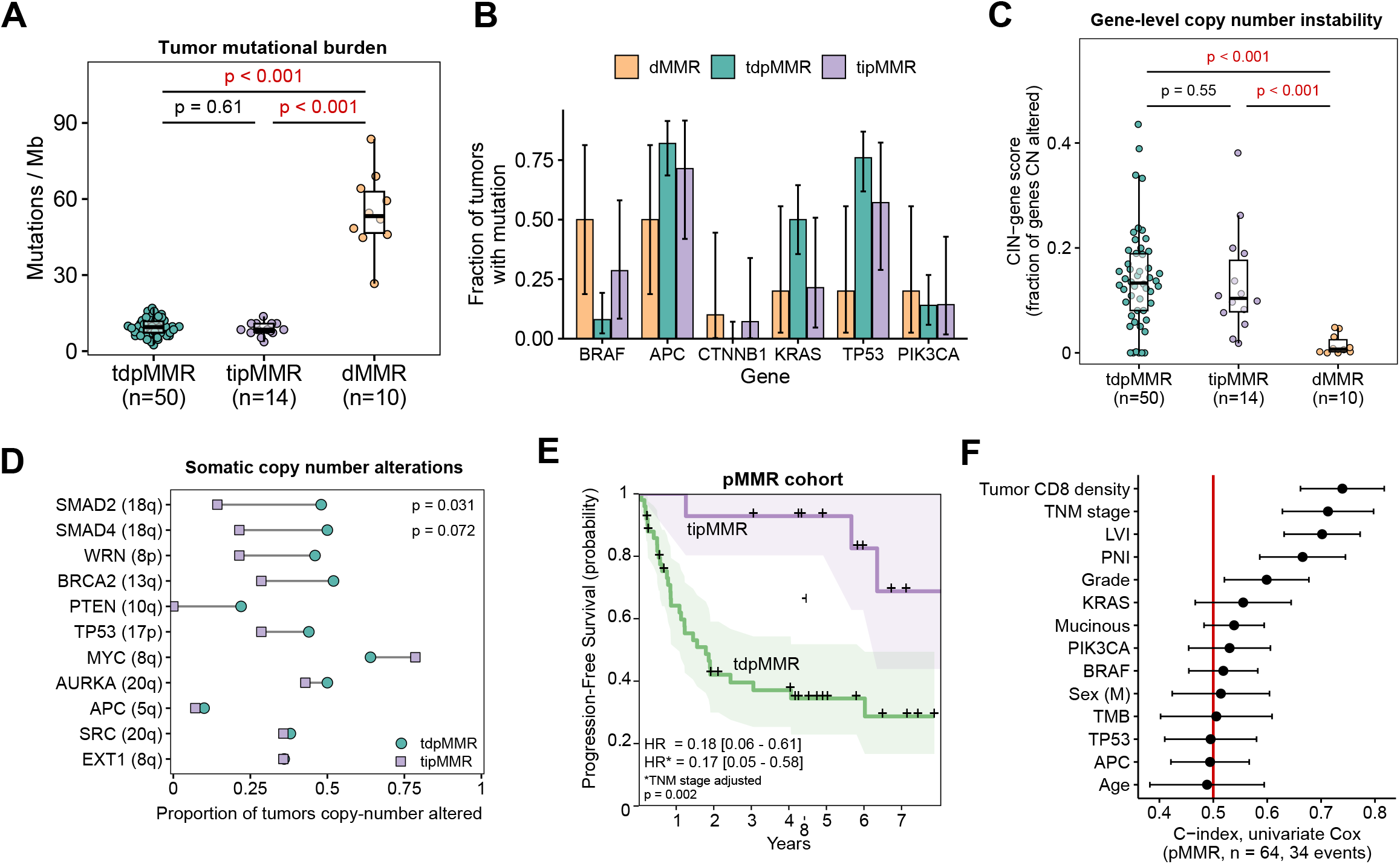
tipMMR status is associated with improved clinical outcomes, independent of molecular and histopathological factors. (A) Tumor mutational burden (y-axis) as calculated from OncoPanel targeted panel sequencing, among dMMR (n = 10), tdpMMR (n = 50) and tipMMR (n = 14) tumors; boxes show the median and interquartile range (IQR), whiskers extend to the most extreme values within 1.5 × IQR, and every tumor is shown; p values, two-sided Wilcoxon rank-sum test. (B) Fraction of dMMR, tdpMMR and tipMMR tumors carrying a mutation in each gene; error bars, exact (Clopper-Pearson) 95% confidence intervals. (C) Gene-level copy-number instability (CIN-gene score: proportion of the 496 reported genes with an absolute copy-number call of 1 or greater; see Methods) in all 74 tumors (dMMR n = 10, tdpMMR n = 50, tipMMR n = 14); box plots and statistics as in (A). (D) Proportion of tipMMR (squares) and tdpMMR (circles) tumors carrying a copy-number alteration at each of 11 canonical CRC chromosomal instability loci, ordered by the magnitude of the difference. Nominal Fisher’s exact p values are shown where p < 0.1; no locus remained significant after correction for multiple testing across the 11 loci (Benjamini-Hochberg q > 0.3 for all). (E) Kaplan-Meier curve for tipMMR (n=14) vs tdpMMR (n=50; dMMR tumors not included in this analysis). Shaded bands, 95% confidence intervals; HR, Cox hazard ratio (tipMMR vs tdpMMR); HR*, adjusted for TNM stage; p, log-rank test. (F) Concordance indices (C-index) of univariate Cox proportional hazards models for PFS in patients with pMMR tumors (n = 64; 34 events), one model for each of 14 molecular, histopathological, clinical and CD8 infiltration features. Points, C-index; bars, 95% confidence intervals; red line, C-index = 0.5 (no discrimination).

We next asked whether tipMMR tumors were enriched for mutations in known CRC driver genes (**Figure 2B**; **Supplementary Table 2**). Across the 94 genes recurrently mutated in this cohort (as defined by a protein-altering variant found in at least three tumors), no genes reached significance when tipMMR and tdpMMR tumors were compared (Benjamini-Hochberg q > 0.9 for all comparisons). Frameshift variants tend to generate more immunogenic neoepitopes than missense substitutions^16^ but we observed no significant difference in frameshift burden between tipMMR and tdpMMR tumors (p = 0.24). No pMMR tumor carried a pathogenic *POLE* or *POLD1* exonuclease-domain variant. tipMMR and tdpMMR tumors were also indistinguishable by clonality as assessed by median variant allele fraction (p = 0.56; read counts available for 61 of the 64 pMMR tumors).

Because chromosomal instability has recently been associated with pathological response to neoadjuvant ICI in pMMR colon cancer^4^, we examined aggregate copy number burden (**Supplementary Table 3)** by calculating the proportion of assessed genes with a copy number alteration (**Extended Figure 2A-B**). This separated the near-diploid dMMR from pMMR tumors (p < 10^-4^), but did not distinguish tipMMR from tdpMMR tumors (p = 0.55; **Figure 2C**). We found that tdpMMR tumors were enriched for 18q deletions encompassing *SMAD2* and *SMAD4*, although this was not significant after correcting for multiple hypothesis testing (**Figure 2D**). Thus, within the limits of clinical panel sequencing, we did not find that the mutation profile, mutational burden, clonality, or frameshift burden distinguished tipMMR from tdpMMR tumors.

tipMMR status was a strong predictor of superior PFS (TNM stage adjusted HR 0.17, 95% CI 0.05-0.58, p = 0.005, Wald test; **Figure 2E)** consistent with Immunoscore data^5^. In other aspects, tipMMR tumors were similar to tdpMMR tumors: both tipMMR and tdpMMR tumors were significantly more likely to be located distally than dMMR tumors (p = 0.047 and p = 0.001, respectively, Fisher’s exact; **Extended Figure 2C)**. We found no significant difference in age among the MMR/T cell groups (p > 0.05; **Extended Figure 2D**). tipMMR tumors in our cohort trended towards a lower TNM stage than tdpMMR and dMMR tumors (**Extended Figure 2E)** and tended to have less aggressive histopathological features than tdpMMR or dMMR tumors, including lower rates of LVI and PNI (**Extended Figure 2F**). We also found no significant difference in the number of patients who received neoadjuvant chemotherapy among the three groups (p = 0.45, Fisher’s exact). To determine the relative impact of molecular, spatial-immune and clinical factors within our pMMR cohort, we used univariate Cox proportional hazards modeling across 14 different features. We assessed each model’s concordance index (C-index), a measure of the accuracy for predicting PFS outcomes and found that the density of tumor-infiltrating CD8^+^ T cells had the highest C-index (0.74, 95% CI 0.66 - 0.82), followed by TNM stage (0.71, 95% CI 0.63 - 0.80) and LVI (0.70, 95% CI 0.63 - 0.77; **Figure 2F**).

### tipMMR tumors match dMMR tumors in cytotoxic effector T cell abundance and tumor cell engagement

A key question about the tipMMR TME is whether infiltrating lymphocytes have the properties expected of antigen-experienced and tumor-directed T cells. To address this question, we performed deep 49-plex immunophenotyping using cyclic immunofluorescence (CyCIF) on 26 specimens (CyCIF; 13 tdpMMR, 4 dMMR, and 9 tipMMR; **Figure 1A**)^17^. We defined CD8^+^ cytotoxic effector T lymphocytes (CTLs) as CD3^+^ CD8^+^ T cells expressing the activation/exhaustion marker PD1 and containing secretory granules positive for the cytolytic serine protease granzyme B (GZMB). When the density of CD3^+^ CD8^+^ PD1^+^ GZMB^+^ cells was quantified in the tumor compartment, we found that tipMMR and dMMR tumors contained 15 to 30-fold more CTLs than tdpMMR tumors (p < 0.001 and p = 0.002, respectively; **Figure 3A-B**). No significant difference in the number of CD8^+^ PD1^+^ GZMB^+^ cells was observed between tipMMR and dMMR tumors (p = 0.60; **Figure 3B**). We also scored for the presence of granulysin (GNLY), a pore-forming cytolytic granule component co-expressed in some T cells with GZMB (**Extended Figure 3A**); CD3^+^ CD8^+^ PD1^+^ GNLY^+^ cells were also more abundant in tipMMR and dMMR than tdpMMR tumors (p < 0.001 and p = 0.002, respectively; **Figure 3C-D**). To determine whether there was a difference in the relative abundance of CTLs per CRC group we normalized for total CD3^+^ or CD8^+^ T cell infiltration. We found that dMMR tumors had a 5 to 7-fold greater fraction of T cells in a CTL state relative to tdpMMR tumors (p ≤ 0.002), and tipMMR tumors a 2 to 5-fold greater fraction, significant among CD3^+^ cells as a whole (p = 0.02) but not among the CD8^+^ subtype cells (p = 0.39; **Extended Figure 3B-C)**. As shown below, this result is explained by the presence of cytotoxic CD4^+^ cells. Thus, tipMMR and dMMR tumors are infiltrated to a similar degree by antigen-experienced PD1^+^ cytolytic effector CD8^+^ T cells, a biologically relevant observation since the high level of CTLs in dMMR tumors is thought to explain their lower rate of progression^18^.

**Figure 3.**
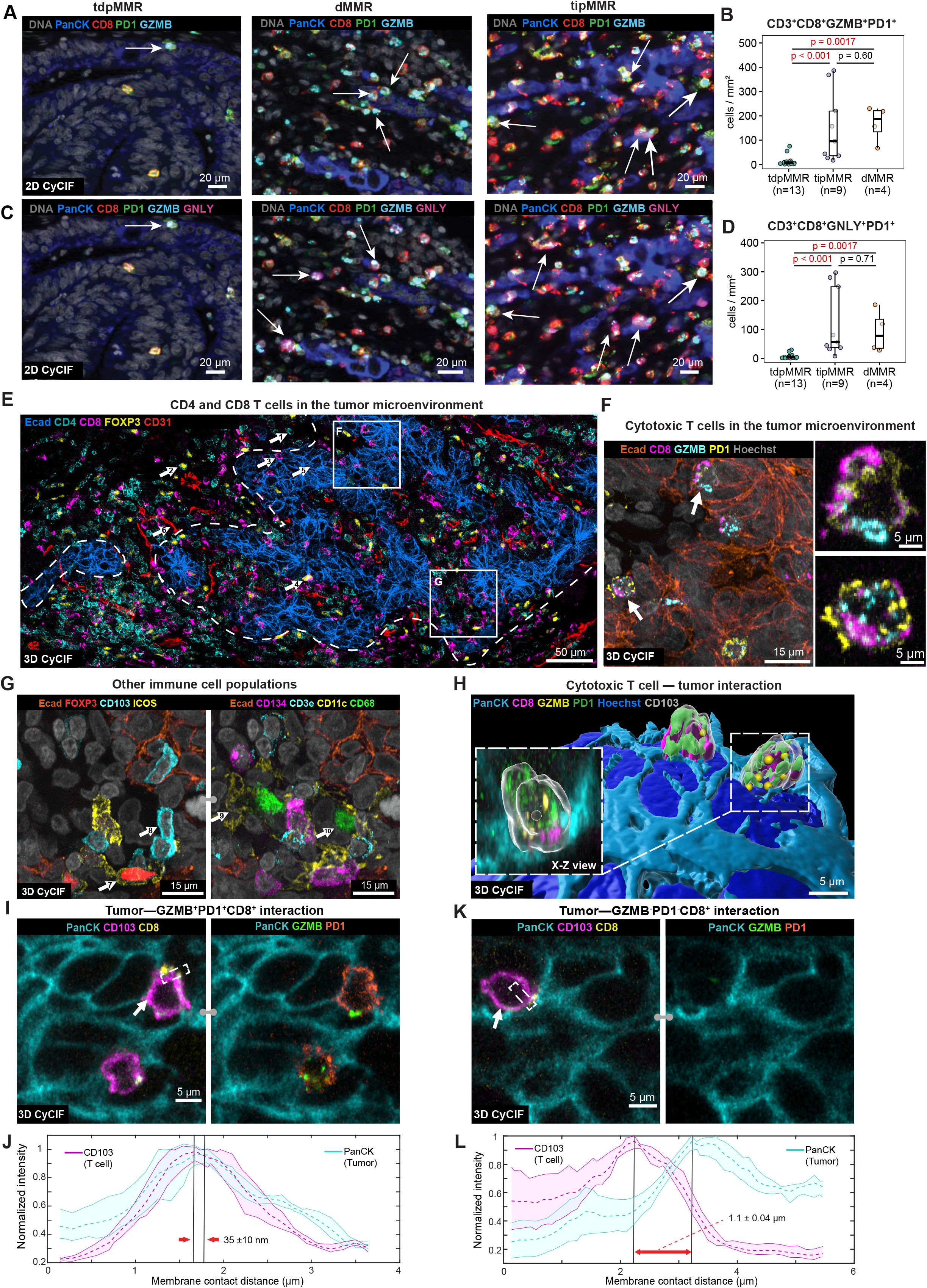
tipMMR and dMMR tumors contain high levels of cytotoxic T cells. (A) 2D CyCIF images of a representative CRC tumor from each MMR class (tdpMMR, dMMR, tipMMR) depicting infiltration of CD8^+^ PD1^+^ Granzyme B^+^ (GZMB^+^) cells. (B) Density of CD3^+^ CD8^+^ PD1^+^ Granzyme B^+^ cells in the tumor compartment (2D CyCIF; tdpMMR n = 13, tipMMR n = 9, dMMR n = 4). (C) 2D CyCIF images of a representative CRC tumor from each MMR class (tdpMMR, dMMR, tipMMR) depicting infiltration of CD3^+^ CD8^+^ PD1^+^ Granulysin^+^ (GNLY^+^) cells with Granzyme B. (D) Density of CD3^+^ CD8^+^ PD1^+^ Granulysin^+^ cells in the tumor compartment (2D CyCIF; n as in B). In (B) and (D), density is cells per mm^2^ of the painted tumor-compartment area (see Methods); boxes show the median and interquartile range (IQR), whiskers extend to the most extreme values within 1.5 × IQR, and every tumor is shown; p values, two-sided Wilcoxon rank-sum test. (E) 3D multiplexed immunofluorescence image showing spatial distribution of immune cells and tumor epithelium in a tumor microenvironment. Markers include ECAD (tumor epithelium, blue), CD4 (cyan), CD8 (magenta), FOXP3 (Tregs, yellow) and CD31 (vasculature, glow). White arrows point to representative cell types (1: helper T cell, 2: CD4+ regulatory T cell, 3: cytotoxic T cell, 4: CD8+ regulatory T cell, 5: tumor cell, 6: vasculature). Scale bar: 50 μm. The image is shown as a maximum intensity projection (MIP). (F-G) Zoomed-in view of two regions of interest (ROIs) from (E) showing spatial distribution of cytotoxic T cells, regulatory T cells, tissue-resident memory T (TRM) cells, dendritic cells, and macrophages in the tumor microenvironment. Markers include ECAD (tumor epithelium, orange) and Hoechst (nucleus, gray); in (F), CD8 (CD8 T cell, magenta), GZMB (cyan), PD1 (yellow). In (G), FOXP3 (Treg, red, left), CD103 (TRM, cyan, left), ICOS (T cell costimulatory receptor, yellow, left), CD3e (T cell, cyan, right), CD11c (dendritic cell, yellow, right), CD134 (T cell costimulatory receptor, magenta, right), and CD68 (macrophage, green, right). The images are shown as a MIP. (H) 3D renderings of two CD8⁺ CD103⁺ PD1⁺ GZMB⁺ cytotoxic T cells in direct contact with the PanCK⁺ tumor surface. Tumor epithelium is shown in light blue, with embedded T cell markers overlaid (CD8, magenta; GZMB, yellow spots; PD1, green; CD103, transparent; Hoechst, blue). (I) High resolution images of two CD103⁺ CD8⁺ cytotoxic T cells interacting with PanCK⁺ tumor epithelium (cyan). Left: CD103 (magenta) and CD8 (yellow) expression; Right: co-expression of GZMB (green) and PD1 (orange) within the same T cells. Dashed box highlights T cell-tumor interface, white arrows point to interacting cells. (J) Line profile analysis showing normalized intensity of tumor and cytotoxic T cells across the interface of interaction. Peak-to-peak distance of 35 ± 10 nm indicates tight contact, also referred to as type I interaction^19^. (K) Similar imaging analysis as in (I) for a CD103⁺ CD8⁺ T cell lacking GZMB and PD1 expression. (L) Line profile analysis of tumor-naive CD8⁺ T cell interaction with PanCK⁺ epithelium, showing a broader peak-to-peak distance of 1.1 µm, suggesting looser contact, also referred to as type III interaction^19^.

To localize CD8^+^ CTLs and map their interactions with neighboring tumor cells, we performed high resolution 3D CyCIF on thick (35 μm) specimens^19^ (**Figure 3E, F**). 3D CyCIF enables cell-cell contacts to be mapped precisely without the cell fragmentation typical of conventional histology on 5 µm thick specimens. However, it is restricted to ∼1 mm x 1 mm regions of interest versus conventional 2D CyCIF, which is a whole slide imaging approach. Tumor cells were identified in 3D images based on PanCK and Ecad staining, endothelial cells and vasculature with CD31, and immune cells subtyped using CD4, CD8, CD103, PD1, GZMB, CD68, and CD11c as well as and immune checkpoint proteins ICOS, CD134, and LAG3 (**Figure 3G**, **Supplementary Table 6**). In 3D images, 99% of CD8^+^ PD1^+^ GZMB^+^ cells were also CD103^+^; CD103 marks intraepithelial lymphocytes that bind ECAD, suggesting that cytotoxic, antigen-experienced CD8 cells were largely tissue-resident (**Figure 3H; Extended Figure 3D-E**). We observed CD8^+^ and CD4^+^ T cells in both stromal regions between sheets of epithelial cells (the lamina propria) and within glandular structures in the tumor compartment (**Figure 3E**).

The number of GZMB-containing lytic granules in CD8 CTLs within the tumor compartment (N=77 CTL cells out of 143 total T cells) ranged from 2 to 27 and did not differ significantly from CTLs in the stroma (p = 0.41; **Extended Figure 3F, G**). In a typical interaction between a PD1^+^ GZMB^+^ CD8 T cell and an adjacent tumor cell, we determined the intermembrane separation to be 35 ± 10 nm (mean ± SD of 10 line profiles along one interface; **Figure 3I-J**), whereas in a single PD1^NEG^ GZMB^NEG^ CD8 T cell the separation from the nearest tumor cell was 1.10 ± 0.04 μm (**Figure 3K-L**). We have previously proposed that intermembrane distances of ∼30-50 nm represent immune synapse formation whereas distances of ∼1 μm are typical of cells not engaged in structural or signaling interactions^19^. Taken together, these data demonstrate that CTLs in tipMMR specimens have the properties expected of active tumor recognition and cell killing.

### tipMMR and dMMR tumors have elevated immune checkpoint expression

Results from the NICHE trial^4^ in pMMR CRC and CheckMate 8HW^20^ in metastatic dMMR CRC have drawn attention to the superior outcomes possible with use of ICIs in CRC^11^/19/26 3:55:00 PM. In our cohort, we observed a significantly higher density of CD3^+^ CD8^+^ PD1^+^ cells per area of tumor in dMMR and tipMMR as compared to tdpMMR tumors (p = 0.002 and p < 0.001 respectively; **Figure 4A-B**) but no difference between tipMMR and dMMR tumors (p = 0.67). LAG3 is another checkpoint protein targeted therapeutically in some cancer types^21^; we found that CD8^+^ LAG3^+^ PD1^+^ T cells were also more abundant in tipMMR and dMMR tumors than tdpMMR tumors (p = 0.003 and p = 0.002, respectively; **Figure 4C-D**). Similarly, if the density of CD8^+^ cells in the tumor compartment expressing any quantified checkpoint protein (LAG3, PD1, or CTLA4) is taken as a measure of prevalence, tipMMR and dMMR were significantly more infiltrated than tdpMMR tumors (p < 0.001 and p = 0.002, respectively; **Figure 4E**). We conclude that tipMMR tumors have elevated levels of T cells expressing one or more checkpoint proteins potentially amenable to ICI-based reactivation.

**Figure 4.**
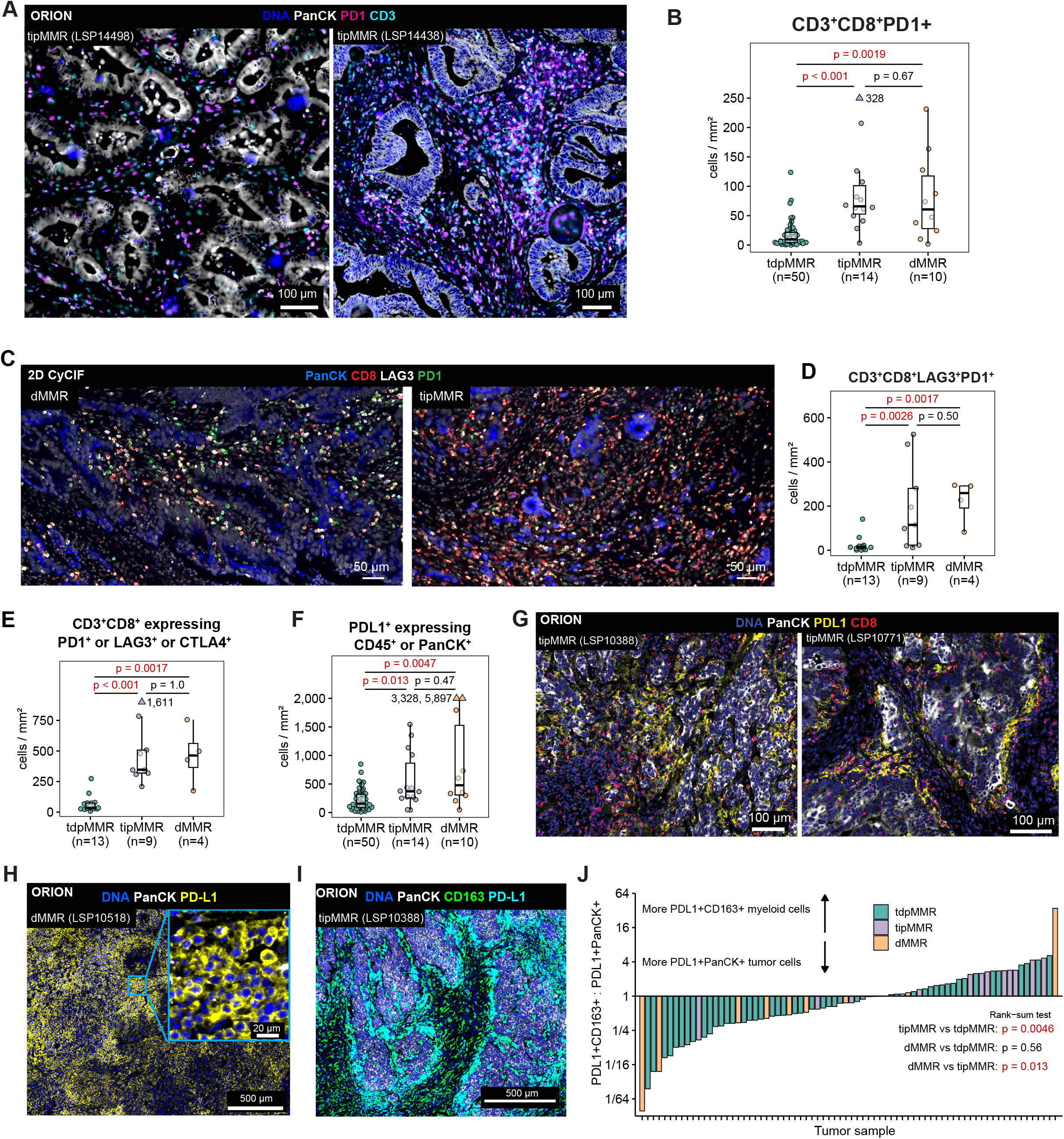
tipMMR tumors exhibit increased immune checkpoint expression. (A) Orion multiplex images highlighting PD1 (magenta) expression for exemplar tipMMR tumors. (B) Density of CD3^+^ CD8^+^ PD1^+^ T cells in the tumor compartment (Orion; tdpMMR n = 50, tipMMR n = 14, dMMR n = 10). The y axis is truncated at 250 cells/mm^2^; one tipMMR tumor above it (328 cells/mm^2^) is drawn as a triangle at the top of the axis, labeled with its value, and is included in the box plot and statistics. (C) 2D CyCIF images highlighting LAG3 (white) and PD1 (green) expression on CD8s (red) in exemplar dMMR and tipMMR tumors. (D) Density of CD3^+^ CD8^+^ LAG3^+^ PD1^+^ T cells in the tumor compartment (2D CyCIF; tdpMMR n = 13, tipMMR n = 9, dMMR n = 4). (E) Density of CD3^+^ CD8^+^ T cells expressing at least one checkpoint protein (PD1, LAG3 or CTLA4) in the tumor compartment (2D CyCIF; n as in D). The y axis is truncated at 900 cells/mm^2^; one tipMMR tumor above it (1,611 cells/mm^2^) is shown as in (B). (F) Density of PDL1^+^ cells that are also CD45^+^ or PanCK^+^ in the tumor compartment (Orion; n as in B). The y axis is truncated at 2,000 cells/mm^2^; two dMMR tumors above it (3,328 and 5,897 cells/mm^2^) are shown as in (B). (G) Orion multiplex images highlighting PDL1 (yellow) staining in exemplar tipMMR tumors. (H) dMMR tumor (LSP10518) with exceptionally high expression of PDL1 on PanCK+ tumor cells. PDL1 staining (yellow) is membranous, consistent with the known cellular localization of PDL1. (I) Orion image of tipMMR tumor expressing CD163 (green) and PDL1 (cyan). (J) Ratio of PDL1^+^CD163^+^ myeloid cells to PDL1^+^PanCK^+^ tumor cells in the tumor compartment (cells positive for both CD163 and PanCK excluded) and colored by group. Boxes show the median and interquartile range (IQR), whiskers extend to the most extreme values within 1.5 × IQR, and every tumor is shown. In (B, D-F, J), p values, two-sided Wilcoxon rank-sum test.

PDL1 is the ligand for the PD1 receptor that is targeted by ICIs. PDL1 is normally expressed on macrophages, dendritic cells, and NK cells and can also be expressed by tumor cells. In cancers such as non-small cell lung cancer, the fraction of viable tumor cells expressing PDL1 (the tumor proportion score) and a composite measure of PDL1 expression by all cells (the combined proportion score) can be predictive of ICI response^22^. Across all cell types in our cohort, tipMMR and dMMR tumors contained PDL1^+^ cells at significantly higher densities than tdpMMR tumors (p = 0.013 and p = 0.005, respectively) but were comparable to each other (p = 0.47; **Figure 4F-G**). In both CRC subtypes the identifies of cells expressing PDL1 and their spatial distributions varied considerably among specimens. Only two tumors (both dMMR) exhibited high expression of PDL1 throughout the tumor compartment (>3,000 PDL1^+^ cells/mm^2^; representing 93% and 63% of PanCK^+^ tumor cells, respectively; **Figure 4H**). The median level of tumor PDL1 expression across all samples was much lower at 82 PDL1^+^ PanCK^+^ cells/mm^2^. Qualitatively, PDL1 expression typically involved strong PDL1^+^ staining in palisading bands of CD163^+^ PDL1^+^ myeloid cells found along the periphery of the tumor compartment (**Figure 4I**). This observation is consistent with previous data showing that PDL1 is most commonly expressed in CRCs by macrophages and dendritic cells^23^. The relative prevalence of PDL1 in myeloid and tumor cells varied across CRC subgroups with a preference for myeloid expression in tipMMR relative to dMMR or tdpMMR tumors (p = 0.013 and p = 0.005, respectively; **Figure 4J)**.

### TCF1^+^ T cells in tipMMR are abundant and highly enriched in immune clusters

The immune response to CRC is associated with formation of immune clusters (ICs) such as tertiary lymphoid structures (TLS), which serve to program and activate anti-tumor B and T cells. In CRC, the presence of TLSs is strongly associated with improved overall survival and in other solid tumors it has been shown to predict greater responsiveness to ICIs^24^. Groups of CD20^+^ B cells were identified in our specimens by using k nearest neighbors (KNN) clustering followed by expansion of the IC mask to include all adjacent immune cells (**Figure 5A; Extended Figure 4A-B**). This resulted in a total of 1,407 clusters, 262 of which corresponded to solitary intestinal lymphoid tissue (SILTs) in adjacent normal tissue. The total IC counts in dMMR and tipMMR tumors were significantly higher than those in tdpMMR (p = 0.0018 and p = 0.0047, respectively; **Figure 5B**), but there was no significant difference in IC count between dMMR and tipMMR tumors (p = 0.77).

**Figure 5:**
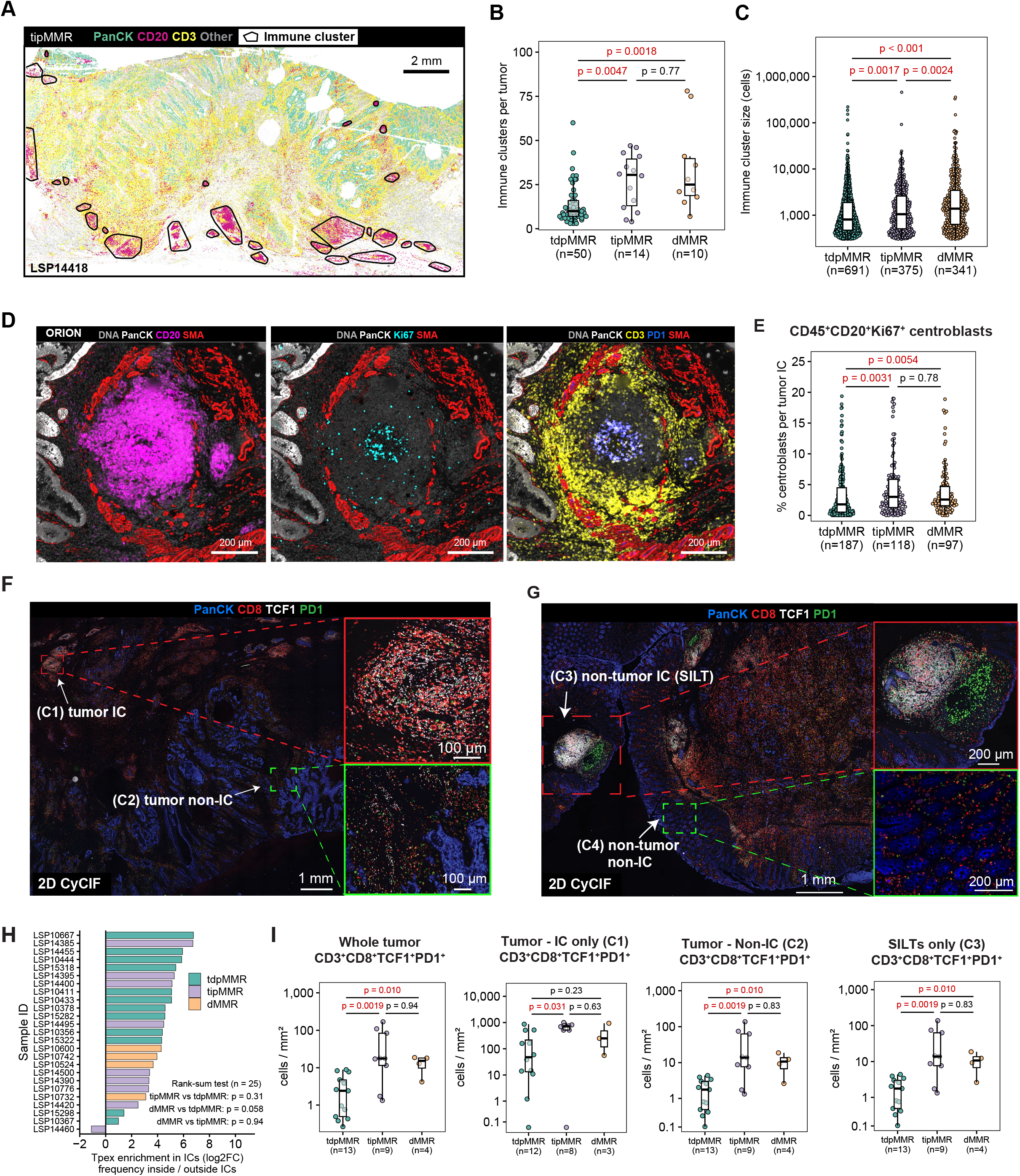
Spatial distribution of immune clusters and TCF1^+^ T cells in tipMMR tumors. (A) Processed whole-slide image showing a subset of markers highlighting immune cells (cells are depicted as circles centered on nuclei and colored based on marker positivity, as indicated) for a representative tipMMR tumor. Computationally detected immune clusters are circled with black lines. (B) Number of immune clusters (ICs) per tumor (tdpMMR n = 50, tipMMR n = 14, dMMR n = 10) and (C) size of each IC (number of cells; 691, 375 and 341 ICs, respectively; log_10_ axis), measured by Orion across the whole slide. Both include all 1,407 ICs, including the 262 solitary intestinal lymphoid tissues (SILTs; ICs with any cell in the annotated normal mucosa). (D) Orion multiplex images of an exemplar immune cluster in a tipMMR tumor highlighting PanCK, CD20, Ki67, SMA, CD3 and PD1 markers. (E) Percentage of CD45^+^CD20^+^Ki67^+^ proliferating B cells (centroblasts) among all cells of each tumor-associated IC, defined as an IC with at least 50% of its cells in the tumor compartment (419 ICs). ICs in which centroblasts made up more than 20% of all cells were excluded as outliers (n = 17), leaving 402 ICs (187 tdpMMR, 118 tipMMR, 97 dMMR); each point is one IC. (F) 2D CyCIF images of TCF1^+^ cells in the tumor: (C1) inside an immune cluster, tumor IC, or (C2) outside a cluster, tumor non-IC. (G) 2D CyCIF images of TCF1^+^ cells outside the tumor: (C3) inside an immune cluster, non-tumor IC, or (C4) outside a cluster, non-tumor non-IC. (H) Log2 fold change (log2FC) of Tpex (CD3^+^ CD8^+^ TCF1^+^ PD1^+^) frequency inside immune clusters (IC) relative to outside ICs, for each individual tumor sample profiled by 2D CyCIF, ordered by value and colored by MMR/T cell status. Frequency is Tpex cells divided by all cells in the region, over the whole slide. Positive values indicate enrichment of Tpex cells within ICs. One tumor with no Tpex cells inside ICs (LSP10422) has no defined log2FC and is omitted (n = 25). Rank-sum tests compare log2FC distributions between MMR subgroups. (I) Density of Tpex cells measured by 2D CyCIF in the whole tumor compartment (C1 + C2; n = 26), tumor ICs (C1; n = 23), tumor non-IC regions (C2; n = 26) and non-tumor ICs/SILTs (C3; n = 25), stratified by MMR status; tumors lacking a region are omitted; tumors with no Tpex cells are shown at 0. In (B, C, E, I), boxes show the median and interquartile range (IQR), whiskers extend to the most extreme values within 1.5 × IQR, and every point is shown; p values, two-sided Wilcoxon rank-sum test (in C and E, ICs pooled within each group).

ICs from tipMMR and dMMR tumors were significantly larger (median 1,051 and 1,385 cells in 2D images, respectively) than those in tdpMMR tumors (median 806 cells; p = 0.002 and p < 0.001, respectively; **Figure 5C**). Intra-tumoral ICs were also more likely to exhibit features consistent with TLS formation and active immune surveillance such as zonation (a central cluster of CD20^+^ B cells surrounded by CD3^+^ T cells; **Figure 5D**) as well as a significantly greater proportion of replicating Ki67^+^ CD20^+^ centroblasts (p = 0.0054 for dMMR vs. tdpMMR and p = 0.0031 for tipMMR vs. tdpMMR; **Figure 5E**). Thus, the proportion of tumor-infiltrating lymphocytes is not only greater in tipMMR than tdpMMR tumors, but also more organized.

Precursor exhausted T cells (Tpex) have recently been identified as a cell population associated with better patient survival in multiple tumor types that can also be activated by ICIs^25,26^. Tpex cells co-express PD1 and TCF1, the master transcriptional regulator of T cell differentiation. Tpex cells retain stem-like properties and are found in tumor-draining lymph nodes, where migratory dendritic cells deliver and present tumor antigen^27^. They have also been reported to occur in ICs in pMMR and dMMR CRC^28^. To quantify Tpex distributions we divided the mucosa, submucosa, and muscularis into four non-overlapping compartments: (C1) ICs located within the tumor domain (‘tumor IC’), (C2) the tumor compartment itself but not including ICs (‘tumor non-IC’), (C3) ICs outside of the tumor domain (‘non-tumor IC’ which corresponds to solitary intestinal lymphoid tissue; SILT), and (C4) the stromal and adjacent normal region not including ICs (‘non-tumor non-IC’) (**Figure 5F-G**). Tpex cells were strongly enriched within ICs (compartments C1 and C3) compared to outside ICs (C2 and C4) across nearly all samples (log2FC > 0 in 24 of 25 tumors) and MMR/T cell classes (**Figure 5H**). Tpex cell density was significantly higher in tipMMR compared to tdpMMR in tumor ICs (C1, p = 0.03; **Figure 5I**), with no difference relative to dMMR. Tpex cell density was also significantly higher in tipMMR (p = 0.002) and dMMR (p = 0.01) compared to tdpMMR in the whole tumor (combining compartments C1 and C2) and in tumor non-IC compartments (C2; **Figure 5I**). In contrast, Tpex density did not differ significantly between MMR subgroups within SILTs (C3). These data indicate that Tpex density is elevated in tipMMR relative to tdpMMR in both IC (C1) and non-IC (C2) tumor compartments, and in dMMR relative to tdpMMR tumors in the non-IC (C2) compartment. Tpex density did not differ between MMR subgroups in SILTs (C3). This suggests a reservoir of potentially ICI-responsive Tpex cells, a promising observation for therapeutic options in tipMMR.

### tipMMR tumors are uniquely infiltrated with FOXP3^+^ regulatory T cells compared with dMMR or tdpMMR tumors

CD4^+^ T cells, comprise both effector populations and FOXP3^+^ regulatory T cells (Tregs). In tipMMR tumors CD4^+^ T cells were distributed spatially both within CD8^+^-rich neighborhoods and in regions largely devoid of CD8^+^ cells (**Extended Figure 5A**). By Orion imaging on all specimens, the density of CD3^+^ CD4^+^ FOXP3^NEG^ cells was ∼2-fold higher in tipMMR than tdpMMR tumors (p = 0.004) and ∼2.5-fold higher in dMMR than in tdpMMR tumors (p = 0.009; **Figure 6A**). By CyCIF on a subset of specimens we found that CD3^+^ CD4^+^ FOXP3^NEG^ GZMB^+^ (cytotoxic CD4 T cells) were 4-fold greater in tipMMR than in tdpMMR tumors (p = 0.007; **Figure 6B**). Using high resolution 3D CyCIF of a dMMR sample we confirmed that cytotoxic CD3^+^ CD4^+^ FOXP3^NEG^ GZMB^+^ T cells were in close contact with tumor cells and in proximity to CD8^+^ GZMB^+^ cells also engaged with tumor cells (**Figure 6C**). We conclude that these CD3^+^ CD4^+^ GZMB^+^ FOXP3^NEG^ cells are also likely to be tumor-reactive and are significantly enriched in both dMMR and tipMMR tumors compared with tdpMMR tumors.

**Figure 6.**
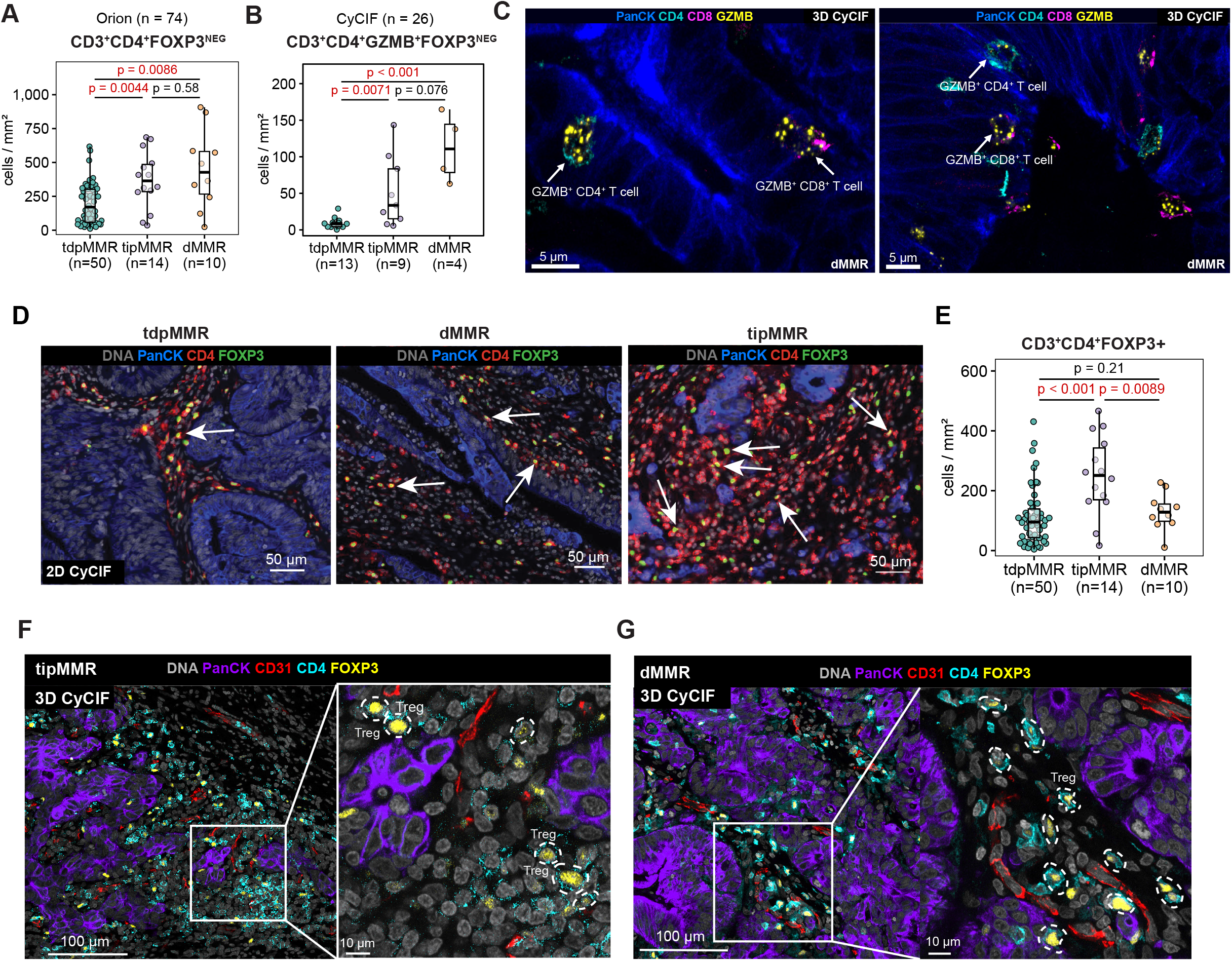
Tregs distinguish the tipMMR immune landscape. (A) Density of CD3^+^ CD4^+^ FOXP3^-^ T cells in the tumor compartment (Orion; tdpMMR n = 50, tipMMR n = 14, dMMR n = 10). (B) Density of CD3^+^ CD4^+^ FOXP3^-^ GZMB^+^ cytotoxic CD4 T cells in the tumor compartment (2D CyCIF; tdpMMR n = 13, tipMMR n = 9, dMMR n = 4). (C) 3D CyCIF image of a dMMR tumor showing spatial distribution of immune cells and tumor epithelium. Markers include Ecad (tumor epithelium, blue), CD4 (cyan), CD8 (magenta), GZMB (yellow). (D) 2D CyCIF images of a representative CRC tumor from each MMR class (tdpMMR, dMMR, tipMMR) depicting infiltration of FOXP3^+^ (green) CD4^+^ (red) Tregs. (E) Density of CD3^+^ CD4^+^ FOXP3^+^ regulatory T cells (Tregs) in the tumor compartment (Orion; n as in A; all tumors shown). (F-G) 3D multiplexed immunofluorescence image (maximum intensity projection, MIP) showing spatial distribution of Tregs in an exemplar tipMMR and dMMR tumor. In (A, B, E), density is cells per mm^2^ of the painted tumor-compartment area (see Methods); boxes show the median and interquartile range (IQR), whiskers extend to the most extreme values within 1.5 × IQR, and every tumor is shown; p values, two-sided Wilcoxon rank-sum test.

We found that the density of CD3^+^ CD4^+^ FOXP3^+^ T regulatory cells was significantly higher in tipMMR tumors than in either tdpMMR (p < 0.001) or dMMR tumors (p = 0.009; **Figure 6D, E**). High resolution 3D imaging in a tipMMR tumor further highlighted intratumoral Treg enrichment (**Figure 6F-G**). Tregs act to suppress T helper cells and CTLs and are targets of emerging immunotherapies^29^. In CRC, however, dense FOXP3^+^ infiltrates have been associated with better outcomes^30^, in part because many tumor-infiltrating FOXP3⁺ cells score as FOXP3^LO^ in flow cytometry and are non-suppressive activated T cells rather than bona fide Tregs^31^. To further characterize Treg phenotype across the samples, we examined ICOS expression, a marker linked to enhanced Treg suppressive activity and co-stimulatory and activation states in other disease contexts^32^. Although the differences were not statistically significant we found that tdpMMR tumors had a generally low ICOS^+^ to ICOS^-^ ratio among Tregs while tipMMR tumors displayed substantial heterogeneity in ICOS expression, and Tregs in dMMR tumors had higher ICOS^+^:ICOS^-^ ratios (**Extended Figure 5B**). These data suggest that, with respect to Treg cells, the TME of tipMMR CRC is not the same as that of dMMR CRC. This difference in Treg density may imply that targeting Tregs in the tipMMR subset should be considered.

### Integration of spatial, genomic and clinical features via cBioPortal

Cancer genetics has benefitted greatly from the availability of data portals that enable comparison of continuous and categorical variables such as survival, genotype, or patient characteristics across cohorts. cBioPortal is the most widely used platform for this type of exploration^33,34^. To enable the joint exploration of spatial features of the TME, we enhanced cBioPortal via integration of the tissue image viewer Minerva^35,36^. For spatial data, concepts analogous to driver and passenger mutations do not yet exist but as a first pass at generalizable “spatial hallmarks”^37^. We incorporated spatial immune features inclusive proportions of tumor, stromal, and immune cell types, their densities inside different compartments (tumor, stroma, and margins), and immune cluster count. Patient stratification now provides for both a TME feature (e.g. PDL1 cell density) and genomic status and demographics (**Figure 7A**). The *Results View* has been expanded to generate OncoPrint plots that include TME features (**Figure 7B**). Primary tissue images for each tumor are also accessible through a linked Minerva window (**Figure 7C**), allowing users to interact directly with primary image data. We anticipate that this extended version of cBioPortal will find broad use as a means of exploring the intersection between tumor genetics and the biology of the TME.

**Figure 7:**
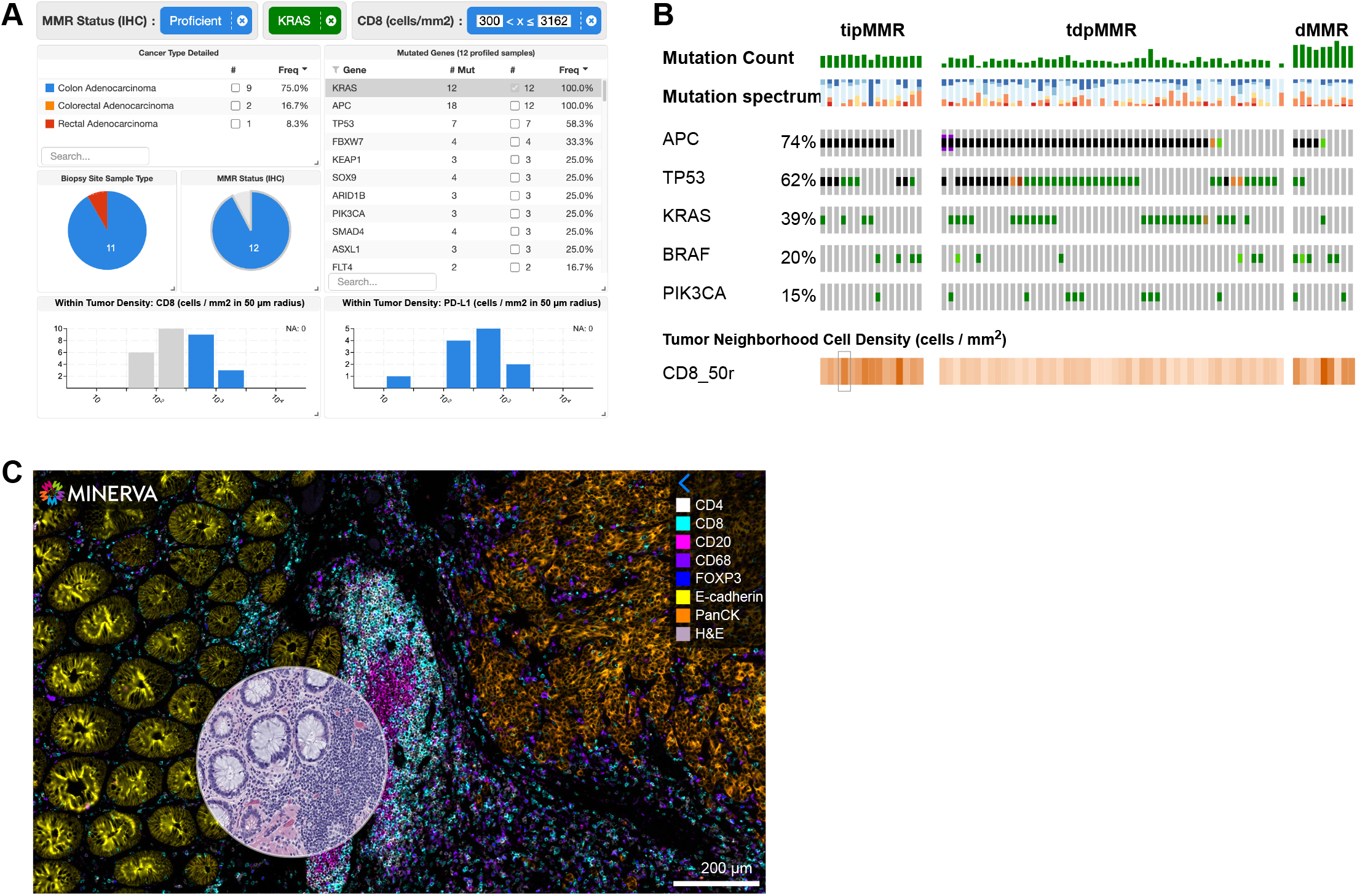
Spatially enhanced cBioPortal facilitates multimodal data exploration. Panels modified from cBioPortal. (A) Subset of pMMR KRAS-mutated patients showing anatomical location, mutation counts, and tumor CD8^+^ and PDL1^+^ cell densities. (B) OncoPrint depicting both mutation and T cell infiltration status. (C) Example of a linked Minerva stories view showing Orion multiplex IF imaging data, with a circular interactive overlay of the same-section H&E.

## DISCUSSION

It has been recognized for over a decade^6,7^ that a subset of pMMR CRCs is lymphocyte-infiltrated and has a better prognosis than other pMMR tumors; recent estimates put this at ∼15% of stage III pMMR CRC^28^. However, potential interactions between MMR phenotype, the degree of infiltration by different types and states of T cells, the potential impact of differences in driver mutations and TMB status, and the presence of other immune cell types has not been examined systematically. By integrating molecular, spatial, and clinical data, we find that this lymphocytic population in the tipMMR subset consists of tumor-engaged effector T cells arising without any detectable genomic correlate and coexisting with a regulatory compartment distinct from that of dMMR or T cell-depleted pMMR tumors. Like dMMR tumors, tipMMR cancers contain numerous GZMB and/or GNLY-positive CD8 CTLs in the tumor compartment and high resolution imaging shows that these CTLs can be found in intimate contact with tumor cells; the estimated intermembrane spacing of ∼30 nm at these sites of contact is likely to reflect formation of a functional immune synapse^19^. Thus, in the absence of any other detectable difference between tipMMR and T cell-depleted tdpMMR tumors with respect to grade, genetics, or prognostic variables, superior survival of patients with tipMMR tumors is likely to be a consequence of enhanced immunosurveillance^8^. This conclusion has implications both for understanding what drives TIL infiltration in solid cancers and for the stratification of pMMR CRC patients for treatment.

Despite their similarities, the immune environment of tipMMR tumors is not identical to that of dMMR tumors: tipMMR tumors harbor higher levels of FOXP3⁺ Treg cells, which may imply that ICIs should be combined with emerging therapies targeting these Tregs^38^. Across all CRCs there are also substantial differences in patterns of expression of checkpoint proteins. For example, we find that only ∼3% of tumors of all types are characterized by high expression of PDL1 on tumor cells, with tipMMR tumors exhibiting the highest proportion of PDL1 expression on CD163⁺ macrophages rather than tumor cells. ICIs targeting PD1 or PDL1 can be effective in both settings^39^ but it may be important to determine whether PDL1 tumor proportion score or combined positive score is most effective with tipMMR CRCs.

Initial trials of ICIs in previously treated pMMR CRC have been disappointing and had the disadvantage of evaluating response following therapy-induced TME remodeling.^3^ However, more recent trials in previously untreated disease are consistent with the possibility that a subset of pMMR CRC (speculatively the tipMMR subset) is ICI responsive. For example, in the phase 2 AtezoTRIBE trial, adding atezolizumab to first-line FOLFOXIRI plus bevacizumab prolonged overall survival with a trend towards improved overall survival in the pMMR subgroup (median 30.8 versus 29.2 months; HR 0.80, 80% CI 0.63-1.02)^40^. A secondary analysis found that benefit from atezolizumab among pMMR tumors was largely confined to those with a high Immunoscore-IC, a quantitative measure of CD8⁺ and PDL1⁺ cell density and proximity (treatment-by-biomarker interaction p = 0.006 for PFS)^41^. In the pMMR cohort of the NICHE trial, pathological response was associated with baseline CD8⁺ PD1⁺ T cell infiltration^42^, and in the expanded cohort responders had higher baseline TCF1 expression and a trend toward a higher proportion of CD103⁺ tissue-resident CD8⁺ T cells^4^. Both features are resolved here at single-cell spatial resolution: tipMMR tumors harbor abundant TCF1⁺ Tpex, enriched within lymphoid aggregates, and 3D CyCIF showed that 99% of their PD1⁺ GZMB⁺ CD8⁺ cells co-express CD103, identifying these T cells as almost entirely tissue-resident. The expanded NICHE cohort also linked response to chromosomal instability^4^, which did not distinguish tipMMR from tdpMMR tumors here, suggesting that T cell infiltration and chromosomal instability may be independent, and potentially complementary, predictors of ICI response in pMMR CRC.

More recently, evidence has emerged for a role for ICI even in pretreated pMMR disease. The BACCI Phase 2 trial evaluating capecitabine and bevacizumab with or without atezolizumab in heavily pretreated mCRC found a significant improvement in PFS (HR 0.66, CI 0.44-0.99) in the pMMR subgroup^43^. We have also reported correlative analyses from a phase 1/2 trial of copanlisib plus nivolumab in heavily pretreated metastatic pMMR CRC, in which clinical responders were distinguished by pre-existing T cell infiltration and macrophage-dominant PDL1 expression^44^. The tipMMR state described here recapitulates that same responder phenotype but in primarily treatment-naïve disease, suggesting that similar TME-remodeling strategies may be expandable to first-line therapy in tipMMR tumors.

Beyond prognostication, is there a potential role for immunoprofiling in prediction of ICI responsiveness in clinical trials? tipMMR tumors contain GZMB/GNLY-positive CD8 CTLs that are also positive for PD1 and CTLA4; both populations are found within a PDL1-high microenvironment. tipMMR tumors also contain substantial numbers of Tpex cells located primarily in TLS and similar lymphoid aggregates in tumor-adjacent tissue; Tpex cells are thought to function as an immunotherapy-recruitable reservoir of T cells^26^. These features of the TME have all previously been shown to promote responsiveness to ICIs in other types of solid tumors and are shared with dMMR tumors^26,39^. An unfortunate aspect of correlative analysis performed on pMMR clinical trials to date is that it is has been incomplete with respect to genomic and immune features, and is not public and so cannot be re-analyzed. In the current cohort study we address this by making all data easily accessible via cBioPortal

A key question for the future will be developing an efficient and reliable means of identifying pMMR tumors with an ICI-poised phenotype. One possibility combines immune and tumor markers with detection of MMR status in a single Orion panel. Since MMR testing is already standard of care, a single combined test of this type might lower the barriers to introduction of immunoprofiling in CRC more broadly.

### Limitations

This study is retrospective rather than prospective and is underpowered to replicate several known genomic predictors with modest impact on survival, such as *BRAF* mutations. We have mitigated technical weaknesses of retrospective analysis by careful histopathology re-review, scoring MMR status both by immunohistochemistry and gene sequencing, and focusing on prognostic features having high effect sizes. We are also limited in TMB estimation by use of targeted panel sequencing (0.75-1.3 Mb of coding sequence), which likely undersamples passenger alterations that could contribute to neoantigen load. Because exome-based analyses have linked neoantigen load to T cell infiltration even within microsatellite-stable CRC^15^, we cannot exclude a contribution of neoantigen burden or quality to the tipMMR phenotype that panel-based TMB would not detect.

## Methods

### Ethics and tissue cohort

The research described in this manuscript complies with and was approved under protocol IRB21-0656, “*Secondary Use of Human Biospecimens”* by the Harvard Medical School Institutional Review Boards (IRB) and also complies with all relevant ethical regulations as reviewed and approved by the Brigham and Women’s Hospital and Dana Farber Cancer Institute (DFCI) IRBs. FFPE tissue blocks were collected using banked tissue obtained after all clinical diagnostic work was completed, from 74 patients with Stage I-IV CRC, with no consideration for patient MMR status, sex (male = 35, female = 39) or mutational or genomics status.

FFPE tissue samples were used after diagnosis, and informed written consent was acquired under DFCI IRB protocol 17-000 and a discarded excess tissue protocol. Two cohorts from the same biobank were assembled, the first with 40 individuals with stage II-IV CRC and the second with 34 individuals with no consideration for the sex of the participants. All samples were collected at the time of initial diagnosis.

### Orion imaging and cell type identification

Orion images were collected and computationally stitched and segmented as described in Lin et al, 2023^10^. Mean single-cell intensity for each marker was computed using MCMICRO^45^ software based on a 5-pixel dilation of the nucleus mask. Marker positivity was determined using a Gaussian mixture model on the mean per-cell marker intensities, as previously described in Lin et al.^10^ Marker panel and antibodies used for Orion imaging are listed in **Supplementary Table 4**. Given the observed diversity of CRC morphologies, we controlled for possible imaging artifacts in several ways. First, we manually reviewed T cell infiltration in both tipMMR and tdpMMR tumors and confirmed that tipMMR tumors displayed strong and specific membranous staining of CD3 and CD8^+^ T cells within the tumor compartment (**Extended Figure 3B**). Second, we tested whether cells scored as double-positive for both CD3 and PanCK - an artifact caused by imperfect cell segmentation and cell-cell overlap, particularly among T cells infiltrating the epithelia of glandular crypts^46,47^ - were present. This artifact affected a median of 12% of CD3^+^ cells but we observed no significant difference in the rate of CD3^+^ PanCK^+^ artifacts among the different MMR/T cell classes (p > 0.05; **Extended Figure 3C**). Third, we compared the number of undefined cell types - those identified by nuclear stains but lacking corresponding positive immunofluorescent staining - between tipMMR, tdpMMR, and dMMR tumors, and found no significant differences (p > 0.05; a median of 16% of all cells, consistent with our previous reports^10^). We also evaluated another metric of T cell infiltration, CD3^+^ cell density. Of the 14 pMMR tumors with the highest CD3^+^ density, 10 were tipMMR, and PFS was similarly stratified by this definition (**Extended Figure 3D-E)**.

### Pathology annotation of Orion images

Expert annotations on the same-section H&E images were obtained from board-certified anatomic pathologists (S.C. and A.G.) as previously described in Lin et al.^10^ Orion multiplex images were then manually annotated in OMERO PathView software on whole-slide images using these annotations. Orion images were annotated with regions of interest (ROIs) for both normal epithelium and tumor compartment. ROIs were then extracted using a custom JavaScript bookmark to obtain a text file containing the positions of all ROI-defining points. Determination of whether a given cell was contained inside an ROI was performed using a ray casting algorithm in *cyftools roi* to solve the “point in polygon” problem, where a point is a cell and the polygon is the ROI. Each cell then received a label as to whether it was part of the tumor or was non-tumor based on expert ROIs.

Occasionally, regional imaging artifacts can lead to false positive and false negative cell-type calling for cells within the artifact region. Artifacts can occur for multiple reasons, including antibody aggregates, tissue folding, optical aberrations, and debris on the slide^47^. To reduce the impact of these artifacts, each Orion multiplex image was reviewed for quality and imaging artifacts were outlined using the ROI function in OMERO. Single-cell quantification tables were then processed with the *cyftools roi* function as above to remove those cells that fell within a region corresponding to or impacted by an imaging artifact. This resulted in the redaction of ∼0.24% of the cells present in the initial dataset.

### Cyclic immunofluorescence (CyCIF)

CyCIF staining of the tissues was performed following previously described protocols^17^. In brief, 5-micron FFPE CRC sections were dewaxed on a BOND RX autostainer (Leica Biosystems, Buffalo Grove, IL, USA), and antigen retrieval was performed with Leica Epitope Retrieval 1 buffer (Cat# AR9961) for 20 minutes. After dewaxing and antigen retrieval, the slides were immediately bleached with 4.5% H_2_O_2_ with LED light to reduce auto-fluorescent background. The antibodies were incubated overnight at 4°C. After washing, the slides were imaged with a CyteFinder scanning fluorescence microscope (RareCyte Inc., Seattle, WA) using 20x objectives. The multiple-cycle images were achieved via repetitive antibody incubation, imaging, and fluorophore inactivation (bleaching). For CyCIF image processing, after the raw images were acquired, image stitching and registration were performed using the ASHLAR^48^ module in the Docker-based NextFlow pipeline MCMICRO^45^ to assemble OME-TIFF images. MCMICRO was subsequently used to segment images and quantify cell fluorescence intensities^45^. For quality control of the single-cell data of the multiplexed images, CyLinter^47^ was used to remove image-processing and segmentation artifacts. To generate binary gates for cell-type identification, assign cell types based on lineage-specific marker expression, and downstream analysis, customized scripts were used in Python, R, and MATLAB. All samples were gated independently using 2-component Gaussian Mixture Modeling and validated by visual inspection. Because Orion and CyCIF data were obtained from adjacent tissue sections, we used non-linear B-spline transformations in Elastix^49,50^ to align images and transfer region-of-interest (ROI) and histopathology annotations. All antibodies used for CyCIF are listed in **Supplementary Table 5.**

### 3D sample preparation and image acquisition

FFPE samples were cut at 35-micron sections onto superfrost slides. Thick section 3D cyclic immunofluorescence was performed similar to its 2D counterpart^17^ but with additional care to minimize sample disintegration, however, primary conjugated antibodies were incubated overnight at 4°C. After 1 hour of washing, the sample was mounted with SlowFade Glass (Thermo Fisher) and a 1.5# coverslip (Thorlabs). The sample was then placed inverted in a slide holder fitted with a spring-loaded clamp, which correspondingly was secured onto the microscope stage in a plateholder. Imaging was performed as described previously^19^. All image data was collected on a LSM980 Airyscan 2 (Carl Zeiss) equipped with a 405nm, 488nm, 561nm, 647nm, and 750nm laser lines, and 5x/0.16NA air, 10x/0.45NA air and 40x/1.3NA oil immersion objective lenses. In ZEN 3.9, a 2D widefield overview scan of Hoechst using the 5x objective lens was used to identify regions of interest for higher resolution imaging at 40x in 3D. Images were sampled at 16-bit at 0.14 microns per pixel in X and Y, and 0.28 microns per pixel in Z for approximately 170 or more optical planes. The pinhole size was set to 35 microns. A focus surface was used to maintain focus. To increase throughput, bidirectional and Frame Fastscanning was used at the highest scan speed. Channels were separated into two tracks: track 1 - Hoechst and Alexafluor 750 (if present). track 2 - Alexafluor 488, Alexafluor 555 and Alexafluor 647. The emission range for Hoechst, Alexafluor 488, Alexafluor 555, Alexafluor 647, and Alexafluor 750 were 380nm-489nm, 499nm-544nm, 579nm-640nm, 660nm-705nm, and 755nm-900nm respectively. Data was saved as separate .czi files for further processing. All antibodies used are listed in **Supplementary Table 6**.

### 3D image processing and registration

To improve signal-to-noise, all cycles acquired on the Zeiss LSM980 were denoised using Zeiss ZEN LSM Plus Processing. Channels were background subtracted by removing a fixed constant gray level from the background. The first cycle (reference) was stitched in ZEN using the Hoechst channel as a reference, and all subsequent cycles were registered to the reference cycle in MATLAB 20 (Mathworks) by translation (in XY and subsequently in Z) followed by elastic deformations using *imregdeform()*. Lastly, all transformations for each cycle were applied to their corresponding channels. All channels were saved as separate TIFF files and imported into Imaris 10.0 (Bitplane) as .ims files for surface rendering and animations.

### Laplacian-of-Gaussian based spot detection

GZMB puncta were detected automatically in 3D by filtering each channel with a Laplacian-of-Gaussian kernel matched to a diffraction-limited spot (σ = 2 voxels), keeping local extrema above a channel-specific intensity threshold as spots, and assigning each spot to the segmented cell that contained it to give per-cell spot counts.

### Somatic mutation testing

OncoPanel targeted exome sequencing was performed for all samples as previously described^11^. Tumors were profiled with any of OncoPanel version 1, 2 or 3, depending on the year of the resection. OncoPanel Version 1 (June 26, 2013 - August 8, 2014) targeted 753 Kbp of exonic DNA spanning 275 cancer genes and 91 introns. OncoPanel Version 2 (August 7, 2014 - November 4, 2016) targeted 826 Kbp of exonic DNA spanning 300 cancer genes. OncoPanel Version 3 targeted 1,315 Kbp of exonic DNA spanning 447 cancer genes. A complete list of included genes is available online at https://researchcores.partners.org/data/wiki_pages/97/OncoPanel.pdf. Each mutation in OncoPanel is reported as belonging to one of five tiers, as described in the above reference. For our study, we considered tumors to have mutations in a gene if they had a protein-altering Tier 1 (well-established alteration with clinical utility), Tier 2 (possible utility for clinical trials, proven clinical utility in other tumor types, or with limited evidence of prognostic association) or Tier 3 mutation. Tier was unavailable for three tumors, in which any protein-altering variant was counted. To calculate the tumor mutational burden, we divided the total number of non-synonymous mutations (of any tier) by the total amount of DNA captured by the corresponding version of OncoPanel. Frameshift burden was calculated in the same way from frameshift variants. Clonality was assessed as the median variant allele fraction of all reported variants in each tumor; read counts were available for 61 of the 64 pMMR tumors. Enrichment of mutations in individual genes between groups was tested with Fisher’s exact test with Benjamini-Hochberg correction.

### Somatic copy-number analysis

Discrete gene-level copy-number calls (−2, deep deletion; −1, loss; 0, neutral; +1, gain; +2, amplification) were obtained from OncoPanel. Because targeted panel sequencing does not provide segment-level data, fraction-of-genome-altered could not be computed directly and a surrogate measure was used. CIN-gene was defined as the proportion of assessed genes with an absolute copy-number call of 1 or greater across the 496 genes reported across specimens. Genes without a reported call in a specimen were treated as neutral.

### Outcomes analysis

PFS and OS outcomes were obtained for each patient from manual review of medical records. PFS was defined as the number of days between primary resection and radiographic progression on cross-sectional imaging, where radiographic progression was defined by increase in the size or number of radiographically evident tumors. This is a composite endpoint of disease-free survival (DFS) for patients with no evidence of disease after resection, with or without adjuvant therapy, and PFS for patients with persistent measurable disease undergoing adjuvant therapy. Outcome analysis was performed using Kaplan-Meier estimation using the “survival” package^51^ (version 3.8.3) in R (version 4.3.2), using the *survfit* function. TNM stage adjustment was performed by adding TNM stage as an ordinal value to a Cox proportional hazards model. tipMMR status was defined by a fixed threshold of 497 cells/mm² for CD8^+^ T cell density within the tumor compartment, corresponding approximately to the upper quartile of the cohort-wide distribution. This threshold was set a priori and was not optimized against outcome data. P values for Kaplan-Meier curves are from log-rank tests.

### Cell density calculation

Cell densities were computed in *cyftools* within the pathologist-annotated tumor compartment (see Pathology annotation of Orion images) as follows. For each specimen, the area of the tumor compartment was estimated by rasterization. Every cell in the compartment painted a disc of 20 µm radius, centered on its nucleus, onto a 1 µm x 1 µm grid, and the compartment area was taken as the total painted area, so overlapping discs were counted once. Because only cell-containing tissue is painted, acellular regions inside the annotation (e.g., pools of extracellular mucin) do not dilute the estimate. The density of a cell type was the number of cells in the compartment positive for all of its defining markers divided by the painted area, in cells/mm^2^. This measure was used to define tipMMR status and for the cell-type densities reported in the text and figures unless otherwise stated. Densities in other compartments (Figure 5I) were computed in the same way, from the painted area of each compartment.

### Immune cluster detection

Immune clusters were identified using the *CallTLS* function in *cyftools*, which implements the following algorithm. First, CD20+ cells were clustered using KNN (in Euclidean space) clustering, labeling B cells with >25% of their 25 nearest neighbors as being part of an immune cluster. CD20+ clusters were then expanded with KNN clustering to include immune cells positive for any marker of CD3, CD4, CD8, CD20, CD45, CD45RO, CD68, CD163, with a minimum immune cell density of 20% of the 100 nearest neighbors. Finally, the DBSCAN algorithm (epsilon = 25) was applied to labeled cells to identify individual clusters, with a minimum cluster size of 300 cells.

For the CyCIF images, a parallel method following the same logic as *cyftools* was implemented in MATLAB. Following the identification of immune clusters, tissue compartments were defined using the cluster masks and tumor regions based on pathologist annotations. Cluster-level measurements (Figure 5C, E) were made per cluster, and clusters were pooled within each T cell/MMR group for testing. Group comparisons used two-sided Wilcoxon rank-sum tests, without correction for multiple pairwise comparisons.

### MMR Immunohistochemistry

An expert pathologist blinded to the molecular status of MMR deficiency analyzed immunostaining by immunohistochemistry for MLH1, PMS2, MSH2 and MSH6. Staining was performed in the BWH pathology core following standard clinical protocols. The loss of expression of MMR proteins was defined as the total absence of nuclear labeling in tumor cells associated with a maintained expression in normal cells (as a positive internal control in the same tissue area).

### Cox proportional hazards modeling

Cox proportional hazards modeling was performed using the “survival”^51^ package (version 3.8.3) in R. Histological features, mutation status and sex were modeled as dichotomous variables (e.g. high-grade or low-grade). TNM stage was modeled as an ordinal number ranging from 1 to 4. Immune infiltration scores (e.g. CD8^+^ density) were modeled as continuous variables. C-index and standard error values were computed from the “concordance” measure of the Cox model, as described in the “survival” package.

### cBioPortal enhancements for the joint analysis of clinical, genomic, and imaging datasets

We made several enhancements to cBioPortal that improve support for imaging and other types of assays: (1) a new resource linking feature, (2) the development of a generic assay data model, and (3) the upload of custom data. These enhancements are all available in the latest version of cBioPortal (v6), allowing anyone with a local installation of cBioPortal to leverage the same functionality. The new resource linking feature enables linking any cohort, patient or sample to a URL or viewer by a cBioPortal study data curator. This facilitated easy integration of the Minerva tissue image viewer on the cBioPortal patient view without making changes to the code. The same functionality could be used for linking any other viewer or clinical system. Over the past few years, we have also developed the flexible generic assay data model, enabling the import of derived features from all kinds of assays, including mutational signature data and now spatial features. In this case we used it to store and visualize the cell type density metrics obtained from MCMICRO. Lastly, the upload of custom data allows users to associate any categorical or numerical data to a patient or sample directly from the cBioPortal website. This decreases the feedback loop between e.g. testing the output of an algorithm and seeing what the results will look like in cBioPortal. In this case one can imagine trying different cell density algorithm parameters on the imaging data, and adding those to a cohort in cBioPortal, immediately being able to explore and assess the quality of the output there using the cBioPortal interface.

## Code Availability

The tools cBioPortal, Minerva and MCMICRO are all open source. The code is available at https://github.com/cbioportal and https://github.com/labsyspharm/. Computation of tumor boundaries, immune cluster identification, and cell densities were performed with open-source cyftools (https://github.com/walaj/cyftools).

## Data Availability

https://www.cbioportal.org/study?id=crc_orion_2024

## Acknowledgements

This work was supported by Ludwig Cancer Research, an Aspire Award from the Mark Foundation for Cancer Research, the Harvard Medical School Quadrangle Fund for Advancing and Seeding Translational Research (Q-FASTR), and by an incubator award from the DF/HCC NCI Cancer Center Core Grant P30 CA006516. JAW is supported by NCI grant 5K99-CA300495. SS is supported by a BWH President’s Scholars Award. cBioPortal is supported by Break Through Cancer, the Gray Foundation, the American Association for Cancer Research (AACR), NCI grants U24 CA274633 (ITCR), U24 CA264028 (GDAN), U24 CA233243 (HTAN), and by a core grant to Memorial Sloan Kettering Cancer Center from the NCI (P30 CA008748). We sincerely thank Marios Giannakis (DFCI) for his assistance in reviewing the manuscript and expert guidance on colon cancer biology.

## Contributions

J.A.W., P.K.S., and K.Q. developed the concept for study. J.A.W., K.Q., Y.L., C.Y., J.-R.L., and B.G. performed experiments, imaging, and analysis. S.C. and A.G. provided pathology expertise. I.B. and N.S. developed software in collaboration with Y.-A.C. and J.M. J.A.W., K.Q., and P.K.S. wrote the manuscript and all authors provided edits and approved the final manuscript. J.A.W. K.Q., Y.L., and C.Y. performed data analysis and prepared the figures. P.K.S. and S.S. provided supervision.

---

**Extended Figure 1:**
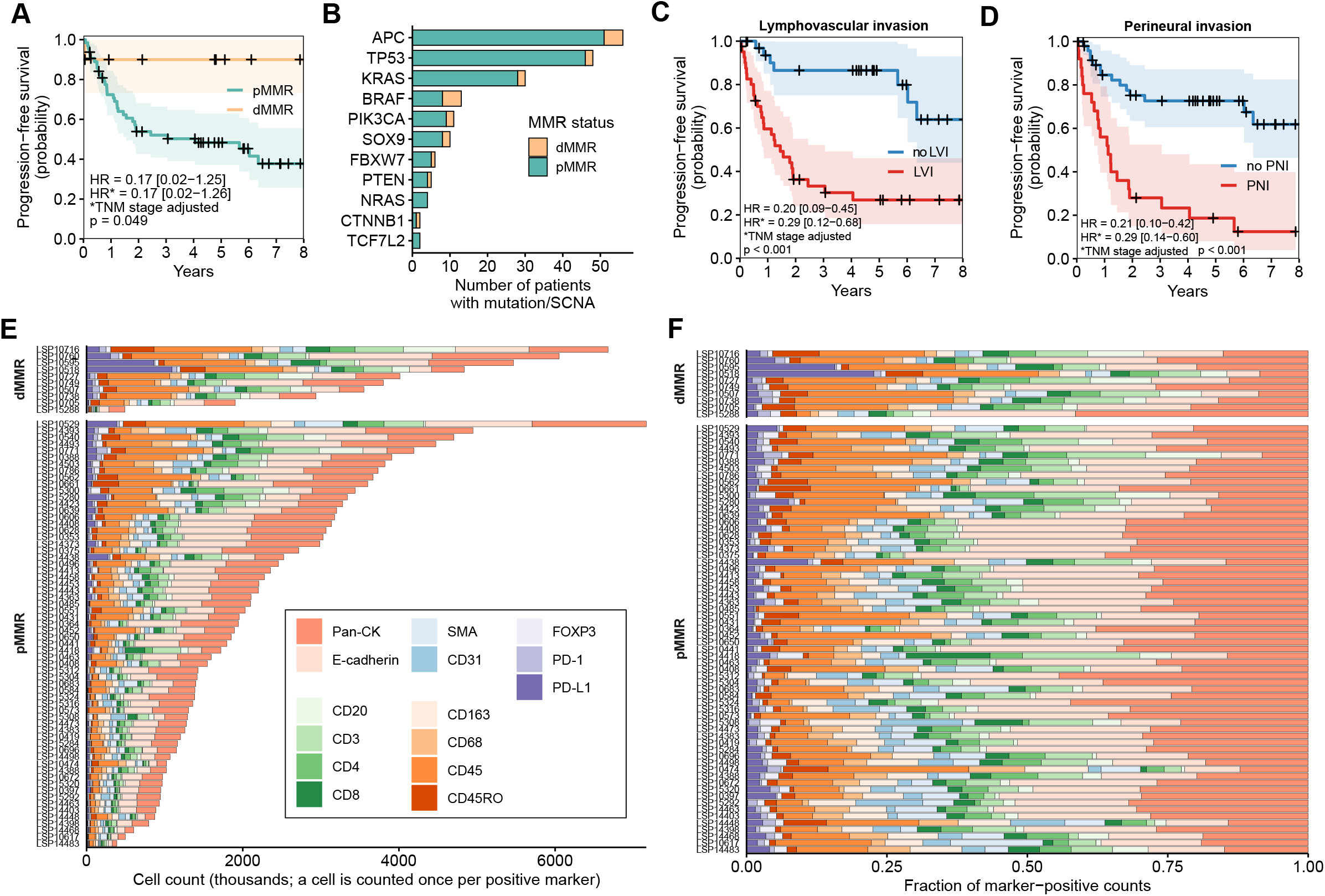
Clinical and cell composition landscape of 74 primary CRCs. (A) Kaplan-Meier curve of probability of progression-free survival stratified by MMR status. (B) Number of patients with a mutation or somatic copy-number alteration (SCNA; deep deletion or amplification) in recurrently altered CRC driver genes, stratified by MMR status (dMMR n = 10, pMMR n = 64). (C) Kaplan-Meier curve of probability of progression-free survival by lymphovascular invasion (LVI) and (D) perineural invasion (PNI) status. (E) Histogram of total cell counts, by marker positivity, across the 74 tumors. Cells can be positive for multiple marks, so total count on X-axis is greater than absolute cell count. (F) The same counts as fractions of each tumor’s total.

**Extended Figure 2:**
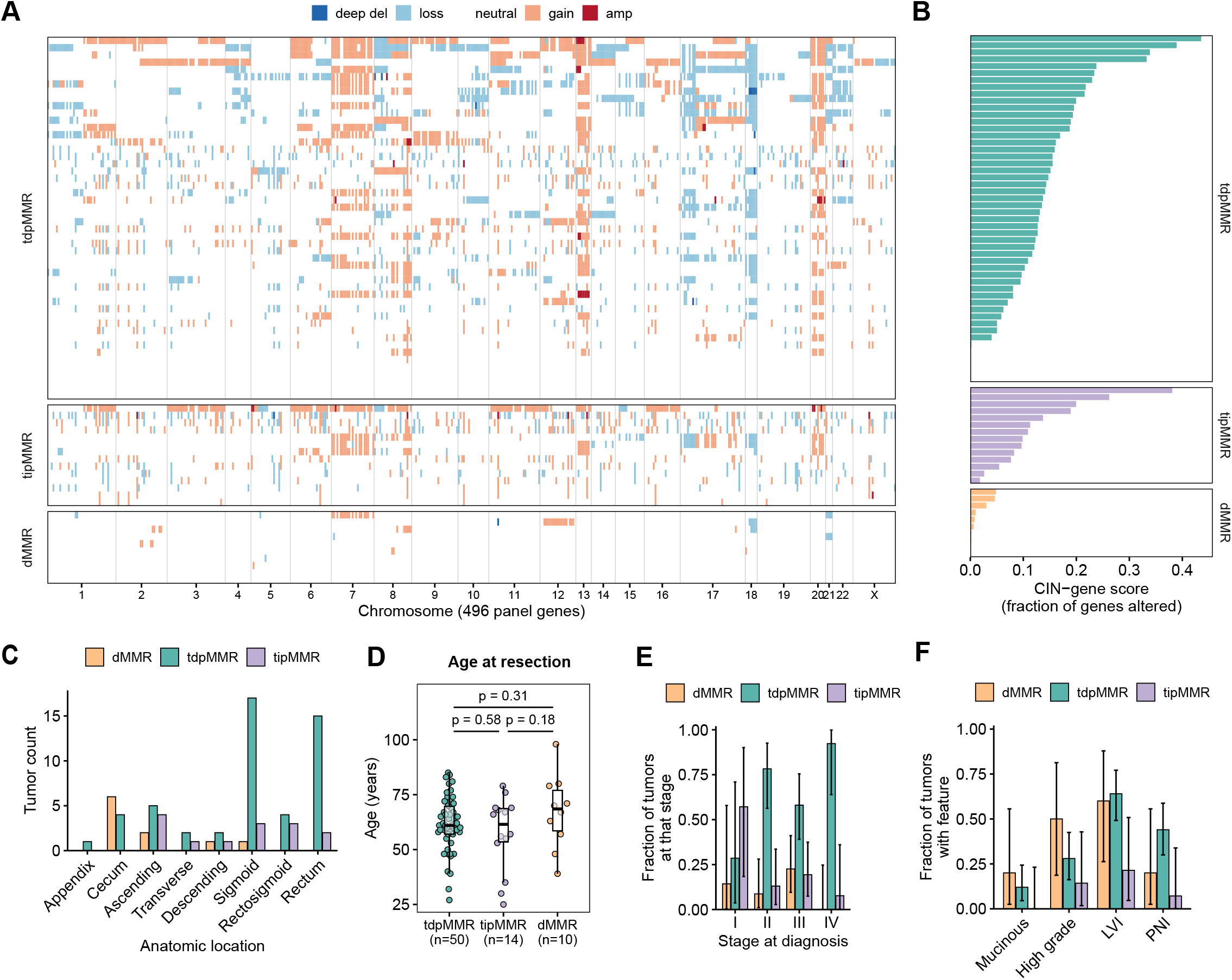
**Copy-number landscape and demographic and anatomic features of CRCs in our cohort,** by MMR/T cell status. (A) Gene-level copy-number landscape across the 496 reported panel genes, ordered by chromosome, for all 74 tumors, grouped by T cell/MMR class and sorted within class by copy-number burden. Calls are colored as deep deletion, loss, neutral, gain or amplification. (B) Per-tumor copy-number instability (CIN) gene-level score, defined as the proportion of assessed panel genes carrying a copy-number alteration, by T cell/MMR class. (C) Tumor anatomical location, (D) age at time of resection (age of diagnosis), and (E) TNM stage at diagnosis. Stage I: n = 1/4/2, II: n = 2/3/18, III: n = 7/6/18, IV: 0/1/12 (dMMR/tipMMR/tdpMMR). (F) Frequency of various histopathological features among different classes of tumors, which differed among the three groups for LVI (p = 0.017) and PNI (p = 0.019) but not for mucinous histology (p = 0.21) or high grade (p = 0.19; Fisher’s exact test).

**Extended Figure 3:**
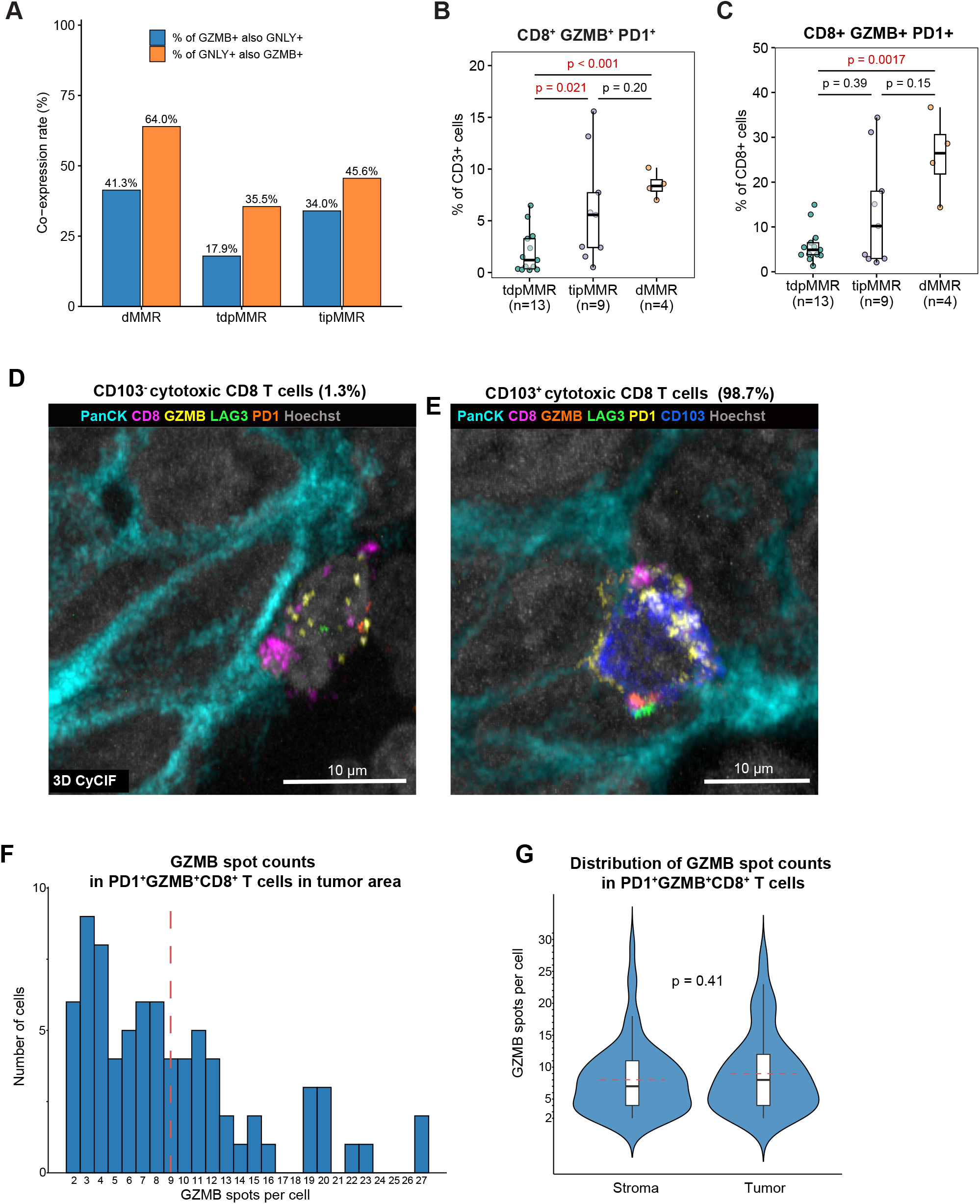
tipMMR tumors exhibit increased cytotoxic T cell activity. (A) Co-expression of Granulysin (GNLY) and Granzyme B (GZMB) among CD3^+^CD8^+^ cells in the tumor compartment across the different T cell / MMR classes, calculated as percentage of GZMB^+^ cells that are also GNLY^+^ (blue) and of GNLY^+^ cells that are also GZMB^+^ (orange), with cells pooled across the tumors of each class (2D CyCIF, n = 26; also in B, C). (B) Percentage of CD3^+^ or C) CD8^+^ that also express CD8^+^PD1^+^GZMB^+^ in CRC tumor regions. Wilcoxon rank-sum test. (D, E) Projections from 3D CyCIF identifying CD103^-^ cytotoxic T cells (1.3%, D) or CD103^+^ cytotoxic T cells (98.7%, E). Scale bar: 10 µm. (F) Distribution of GZMB spot counts per cell in cytotoxic CD8⁺ T cells (PD1⁺GZMB⁺) in the tumor area as measured in 3D CyCIF on a single tipMMR sample. Dashed red lines indicate the mean number of spots. (G) Distribution of GZMB spot counts per cell in cytotoxic CD8⁺ T cells across stromal and tumor regions as measured in 3D CyCIF on a single tipMMR sample. Box plots indicate the median (black line) and interquartile range, with dashed red lines showing the mean number of spots. Wilcoxon rank-sum test, n = 77 cells in the tumor area, n = 66 cells in the stromal area.

**Extended Figure 4.**
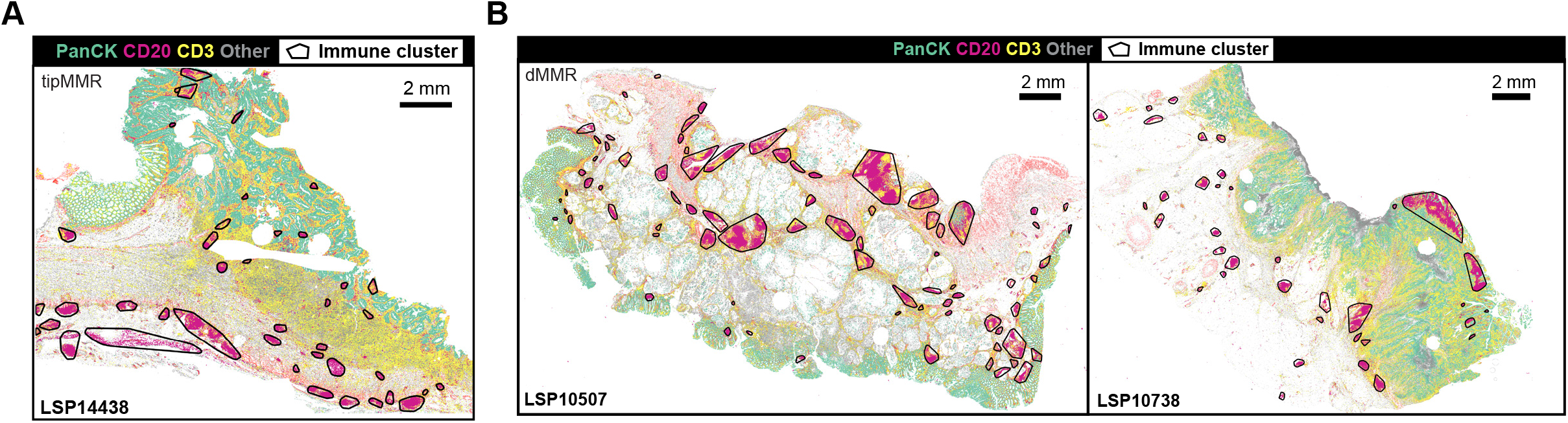
Immune clusters and TCF1^+^ T cells in tipMMR. Pseudoimages derived from spatial single-cell Orion multiplex image data showing PanCK cells (green), CD20 B cells (pink), CD3 T cells (yellow) or all other cells (gray), with approximate immune cluster boundaries superimposed as the convex hull lines of cells labeled as belonging to a distinct immune cluster; for (A) a tipMMR tumor and (B) two dMMR tumors.

**Extended Figure 5:**
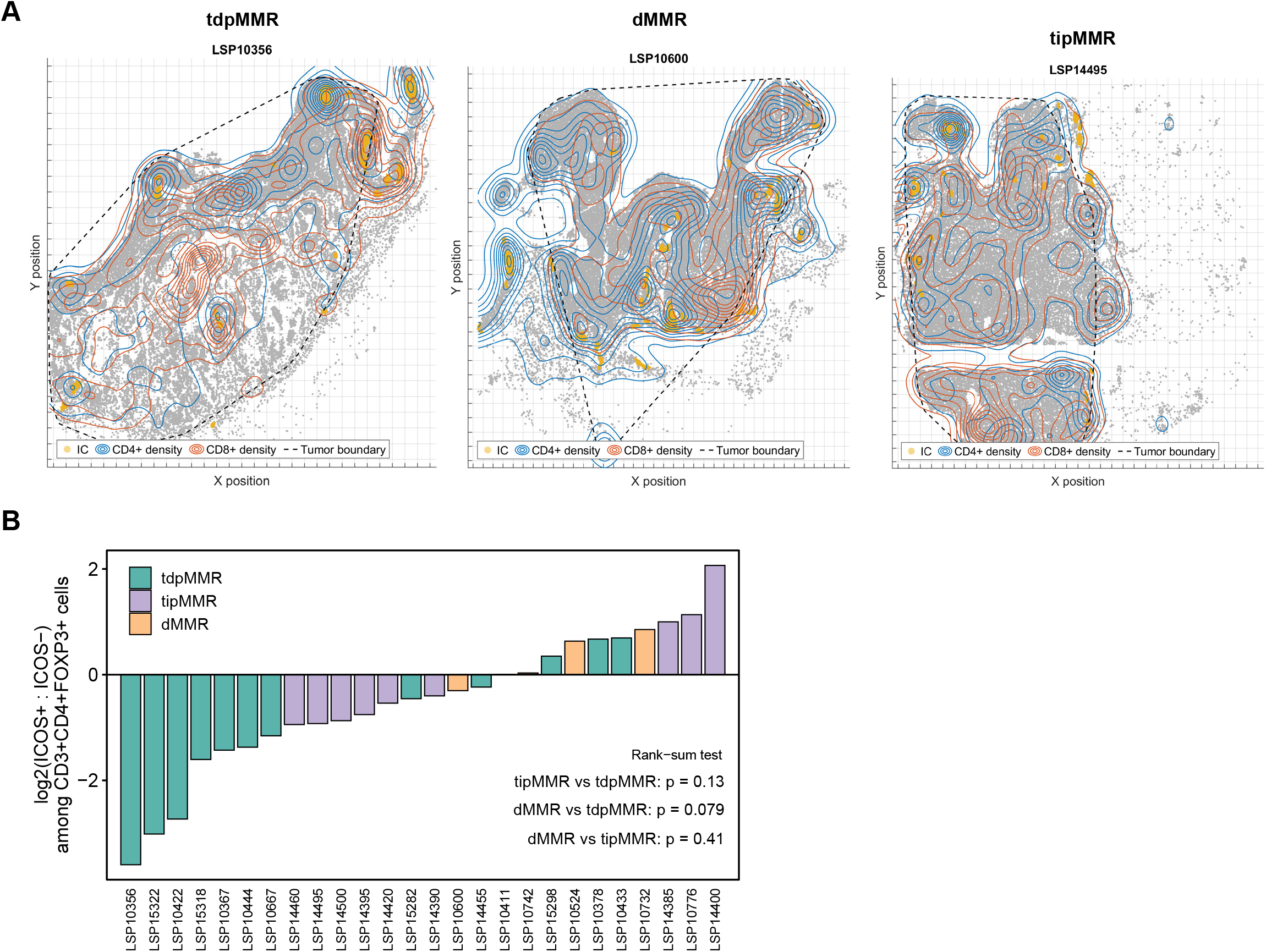
Spatial distribution of CD4^+^ T cells and ICOS expression on Tregs. (A) Digital visualization of representative tdpMMR, dMMR, and tipMMR samples with density contours for CD4+ (blue) and CD8+ (orange). Immune clusters (IC) are depicted in yellow, tumor boundaries by the dashed lines, and gray points represent individual cells. (B) Ranked ratios of ICOS positive to ICOS negative CD3+CD4+FOXP3+ Treg cells among the different T cell / MMR classes. P values derived from Wilcoxon rank-sum testing.

