## Supplementary Tables for "T cell infiltration stratifies mismatch repair-proficient colorectal cancer independently of tumor genotype"

**Supplementary Table 1**: Clinical and abbreviated genomic data. All genes: 0 = not altered, 1 = protein-altering Tier 1-3 mutation (any protein-altering variant in the three tumors without tiers), or deep deletion or amplification

| Specimen_ID | CRC_ID | HTMA402ID | Age | Gender | Primary | Histology | Grade | Location | TNM_At_Diagno | Stage_At_Diagnosis | stage_num | LVI |
|---|---|---|---|---|---|---|---|---|---|---|---|---|
| C01 | CRC2 | 97 | 69 | M | 1 | Adenocarcinoma, with | Low | Rectum | pT3 N0 cM0 | IIA | 2 | 0 |
| C02 | CRC3 | 101 | 40 | F | 1 | Adenocarcinoma | Low | Sigmoid | pT3 N2a cM0 | IIIB | 3 | 1 |
| C03 | CRC4 | 121 | 58 | M | 1 | Adenocarcinoma | Low | Sigmoid | pT3 N0 cM0 | IIA | 2 | 0 |
| C04 | CRC5 | 129 | 56 | M | 1 | Adenocarcinoma | High | Rectosigmoid | pT3 N2a cM0 | IIIB | 3 | 1 |
| C05 | CRC6 | 257 | 73 | F | 1 | Adenocarcinoma | High | Sigmoid | ypT4a N1b cM | IIIB | 3 | 1 |
| C06 | CRC7 | 181 | 58 | M | 1 | Adenocarcinoma | Low | Rectosigmoid | pT4a N2b cM1 | IVA | 4 | 1 |
| C07 | CRC9 | 349 | 67 | M | 1 | Adenocarcinoma | Low | Transverse | pT4a N1a M1c | IVC | 4 | 1 |
| C08 | CRC10 | 73 | 58 | F | 1 | Adenocarcinoma, with | High | Rectum | pT3 N2b cM1b | IVB | 4 | 1 |
| C09 | CRC11 | 81 | 62 | M | 1 | Adenocarcinoma | Low | Transverse | pT4a N1b cM1 | IVA | 4 | 1 |
| C10 | CRC12 | 117 | 25 | M | 1 | Adenocarcinoma | Low | Transverse | pT3 N1b M1a | IIIB | 3 | 0 |
| C11 | CRC13 | 153 | 70 | F | 1 | Adenocarcinoma, Ex-G | High | Appendix | T4a N2a M1c | IVC | 4 | 1 |
| C12 | CRC14 | 245 | 64 | F | 1 | Adenocarcinoma, with | High | Rectum | ypT4a N1b cM | IVC | 4 | 1 |
| C13 | CRC15 | 281 | 39 | F | 1 | Adenocarcinoma | Low | Sigmoid | pT3 N2b cM1a | IIIC | 3 | 1 |
| C14 | CRC16 | 93 | 60 | F | 1 | Adenocarcinoma, with | Low | Sigmoid | pT3 N1b cM0 | IIIB | 3 | 0 |
| C15 | CRC17 | 321 | 39 | F | 0 | Adenocarcinoma, with | Low | Cecum | pT4a N2a cM0 | IIIC | 3 | 1 |
| C16 | CRC18 | 253 | 80 | F | 1 | Adenocarcinoma | Low | Cecum | pT4a N2a cM0 | IIIC | 3 | 1 |
| C17 | CRC35 | 9 | 57 | M | 1 | Adenocarcinoma | Low | Sigmoid | pT4a N0 cM1a | IVA | 4 | 0 |
| C18 | CRC52 | 17 | 71 | M | 1 | Adenocarcinoma | Low | Ascending | pT4a N2a cM1 | IVA | 4 | 1 |
| C19 | CRC81 | 29 | 32 | F | 1 | Adenocarcinoma | High | Sigmoid | pT4a N2b cM0 | IIIC | 3 | 1 |
| C20 | CRC20 | 37 | 27 | F | 1 | Adenocarcinoma | Low | Sigmoid | pT3 N0 cM0 | IIA | 2 | 0 |
| C21 | CRC23 | 49 | 74 | M | 1 | Adenocarcinoma | Low | Rectum | pT3 N0 cM0 | IIA | 2 | 1 |
| C22 | CRC30 | 85 | 55 | M | 1 | Adenocarcinoma | Low | Cecum | ypT4b N1a cM | IVA | 4 | 1 |
| C23 | CRC31 | 89 | 98 | F | 1 | Adenocarcinoma | High | Cecum | yT3 N0 cM0 | IIA | 2 | 0 |
| C24 | CRC36 | 125 | 48 | M | 1 | Adenocarcinoma, with | Low | Sigmoid | pT4a N2b cM0 | IIIC | 3 | 1 |
| C25 | CRC40 | 145 | 65 | F | 1 | Adenocarcinoma | Low | Rectum | pT3 N1c cM0 | IIIB | 3 | 1 |
| C26 | CRC45 | 165 | 85 | F | 1 | Adenocarcinoma | High | Ascending | pT4a N2b cM0 | IIIC | 3 | 1 |
| C27 | CRC56 | 209 | 64 | M | 1 | Adenocarcinoma | Low | Rectum | pT3 N1a cM0 | IIIB | 3 | 0 |
| C28 | CRC59 | 221 | 77 | F | 1 | Adenocarcinoma | High | Rectosigmoid | ypT3 N0 cM0 | IIA | 2 | 0 |
| C29 | CRC68 | 265 | 48 | F | 1 | Adenocarcinoma | High | Cecum | pT4a N2b cM1 | IIIC | 3 | 1 |
| C30 | CRC83 | 325 | 47 | F | 1 | Adenocarcinoma | Low | Sigmoid | pT3 N0 cM0 | IIA | 2 | 1 |
| C31 | CRC85 | 333 | 57 | M | 1 | Adenocarcinoma | Low | Sigmoid | pT4b N1a cM0 | IIIC | 3 | 1 |
| C32 | CRC94 | 369 | 84 | M | 1 | Adenocarcinoma | High | Rectum | ypT4a N1b cM | IIIB | 3 | 0 |
| C33 | CRC28 | 69 | 63 | F | 1 | Adenocarcinoma | Low | Descending | pT1 N0 cM0 | I | 1 | 0 |
| C34 | CRC32 | 105 | 70 | F | 1 | Adenocarcinoma, with | Low | Cecum | pT3 N0 cM0 | IIA | 2 | 0 |
| C35 | CRC43 | 157 | 71 | M | 1 | Adenocarcinoma | High | Cecum | pT3 N1a cM0 | IIIB | 3 | 0 |
| C36 | CRC47 | 173 | 67 | M | 1 | Adenocarcinoma | Low | Cecum | pT3 N1b cM0 | IIIB | 3 | 1 |
| C37 | CRC38 | 137 | 48 | F | 1 | Adenocarcinoma | High | Sigmoid | pT3 N1b cM0 | IIIB | 3 | 1 |
| C38 | CRC71 | 25 | 57 | M | 1 | Adenocarcinoma | High | Ascending | pT3 N2b cM0 | IIIC | 3 | 1 |
| C39 | CRC26 | 61 | 67 | M | 1 | Adenocarcinoma | Low | Ascending | pT3 N0 cM0 | IIA | 2 | 0 |
| C40 | CRC62 | 233 | 59 | M | 1 | Adenocarcinoma | Low | Sigmoid | pT2 N1a cM0 | IIIA | 3 | 1 |
| C41 | CRC42 | 13 | 48 | F | 1 | Adenocarcinoma | Low | Sigmoid | pT4a N0 cM0 | IIB | 2 | 1 |
| C42 | CRC44 | 161 | 67 | M | 1 | Adenocarcinoma | Low | Descending | pT3 N0 cM0 | IIA | 2 | 1 |
| C43 | CRC50 | 189 | 76 | F | 1 | Adenocarcinoma | Low | Ascending | pT3 N2a cM0 | IIIB | 3 | 0 |
| C44 | CRC51 | 193 | 57 | M | 1 | Adenocarcinoma | Low | Ascending | pT3 N0 cM0 | IIA | 2 | 0 |
| C45 | CRC53 | 197 | 69 | F | 1 | Adenocarcinoma | Low | Rectum | pT3 N1b cM0 | IIIB | 3 | 0 |
| C46 | CRC54 | 201 | 55 | F | 1 | Adenocarcinoma | Low | Sigmoid | pT1 N0 cM0 | I | 1 | 0 |
| C47 | CRC55 | 205 | 61 | F | 1 | Adenocarcinoma | High | Rectum | pT2 N0 cM0 | I | 1 | 1 |
| C48 | CRC57 | 213 | 65 | M | 1 | Adenocarcinoma | High | Sigmoid | pT2N0 cM0 | I | 1 | 0 |
| C49 | CRC61 | 229 | 80 | F | 1 | Adenocarcinoma | Low | Rectosigmoid | pT3 N1a cM0 | IIIB | 3 | 0 |
| C50 | CRC63 | 21 | 66 | F | 1 | Adenocarcinoma | High | Ascending | pT3 N2b cM0 | IIIC | 3 | 1 |
| C51 | CRC64 | 237 | 66 | F | 1 | Adenocarcinoma | Low | Ascending | pT3N0 cM0 | IIA | 2 | 0 |
| C52 | CRC69 | 269 | 30 | F | 1 | Adenocarcinoma | Low | Sigmoid | pT3 N0 cM0 | IIA | 2 | 0 |
| C53 | CRC72 | 277 | 59 | M | 1 | Adenocarcinoma | Low | Rectosigmoid | pT3 N0 cM0 | IIA | 2 | 0 |
| C54 | CRC75 | 293 | 66 | M | 1 | Adenocarcinoma | Low | Rectum | ypT3 N0 cM0 | IIA | 2 | 0 |
| C55 | CRC76 | 297 | 62 | M | 1 | Adenocarcinoma | Low | Rectum | pT3 N0 cM0 | IIA | 2 | 0 |
| C56 | CRC77 | 301 | 35 | F | 1 | Adenocarcinoma | Low | Descending | pT2 N0 cM0 | I | 1 | 0 |
| C57 | CRC78 | 305 | 60 | M | 1 | Adenocarcinoma | Low | Rectum | ypT3 N0 cM0 | IIA | 2 | 0 |
| C58 | CRC79 | 309 | 53 | F | 1 | Adenocarcinoma | Low | Rectum | pT1 N0 cM0 | I | 1 | 0 |
| C59 | CRC80 | 313 | 74 | F | 1 | Adenocarcinoma | Low | Rectum | pT3 N0 cM0 | IIA | 2 | 1 |
| C60 | CRC84 | 329 | 48 | F | 1 | Adenocarcinoma | Low | Rectum | pT3 N0 cM0 | IIA | 2 | 0 |
| C61 | CRC90 | 33 | 68 | M | 1 | Adenocarcinoma | Low | Rectosigmoid | pT3 N2a M0 | IIIB | 3 | 0 |
| C62 | CRC91 | 357 | 69 | M | 1 | Adenocarcinoma | Low | Sigmoid | pT1 N0 cM0 | I | 1 | 1 |
| C63 | CRC92 | 361 | 50 | F | 1 | Adenocarcinoma | Low | Sigmoid | pT3 N1b cM0 | IIIB | 3 | 0 |
| C64 | CRC95 | NA | 72 | F | 1 | Adenocarcinoma | High | Cecum | pT4a N1a M0 | IIIC | 3 | 1 |
| C65 | CRC96 | NA | 81 | F | 0 | Recurrent adenocarcin | Low | Sigmoid | pT3 N0 M0 | IIA | 2 | 0 |
| C66 | CRC97 | NA | 79 | M | 1 | Adenocarcinoma | High | Ascending | pT4a N2b M0 | IIIC | 3 | 1 |
| C67 | CRC98 | NA | 83 | M | 1 | Adenocarcinoma | Low | Rectum | pT4a N0 M0 | IIB | 2 | 1 |
| C68 | CRC100 | NA | 61 | M | 1 | Adenocarcinoma | High | Cecum | pT3 N2a M1a | IVA | 4 | 1 |
| C69 | CRC101 | NA | 60 | M | 1 | Adenocarcinoma | Low | Descending | pT4a N2a M0 | IIIC | 3 | 1 |
| C70 | CRC102 | NA | 79 | M | 1 | Adenocarcinoma | Low | Rectosigmoid | pT2 N0 M1a | IVA | 4 | 0 |
| C71 | CRC103 | NA | 61 | M | 1 | Adenocarcinoma | Low | Rectum | pT3 N0 M0 | IIA | 2 | 1 |
| C72 | CRC104 | NA | 76 | F | 1 | Adenocarcinoma | Low | Ascending | pT3 N1b M0 | IIIA | 3 | 1 |
| C73 | CRC105 | NA | 53 | F | 1 | Adenocarcinoma | Low | Ascending | pT3 N2b M1a | IVA | 4 | 1 |
| C74 | CRC106 | NA | 59 | F | 1 | Adenocarcinoma | High | Sigmoid | pT4a N1b pM1 | IVA | 4 | 1 |

| PNI | Deposits | Border | TIL | MMRIHC | MMR_molecula | nt_Mets_At_Diag | Recurrence | Location_Of_Re | Death | PFSCensor | OSCensor | PFSDays | OSDays | OP_TMB | panel_version |
|---|---|---|---|---|---|---|---|---|---|---|---|---|---|---|---|
| 1 | 0 | NA | Absent | Intact | MSS | 0 | 1 | Lung | 0 | 0 | 1 | 397 | 1965 | 15.73531744 | 2 |
| 1 | 0 | Infiltrating | Absent | Intact | MSS | 0 | 1 | Ovaries | 0 | 0 | 1 | 448 | 1932 | 10.8936813 | 2 |
| 1 | 1 | Infiltrating | Absent | Intact | MSS | 0 | 1 | Bowel (Primary | 0 | 0 | 1 | 445 | 1887 | 9.683272268 | 2 |
| 0 | 1 | Infiltrating | Moderate | Intact | MSS | 0 | 1 | Pre-sacral, lung | 1 | 0 | 0 | 459 | 778 | 8.472863235 | 2 |
| 1 | 0 | Infiltrating | Mild | Intact | MSS | 0 | 1 | Liver | 0 | 0 | 1 | 56 | 1534 | 8.472863235 | 2 |
| 1 | 1 | Infiltrating | Absent | Intact | MSS | 0 | 1 | Liver, Peritoneu | 1 | 0 | 0 | 195 | 425 | 10.8936813 | 2 |
| 1 | 1 | Infiltrating | Moderate | Intact | MSS | 1 | 1 | Liver, Periteon | 1 | 0 | 0 | 26 | 673 | 8.472863235 | 3 |
| 1 | 0 | Infiltrating | Moderate | intact | MSS | 1 | 1 | Liver | 0 | 0 | 1 | 681 | 1991 | 13.31449937 | 2 |
| 1 | 0 | NA | Absent | intact | MSS | 1 | 1 | Liver | 1 | 0 | 0 | 275 | 1006 | 8.472863235 | 2 |
| 1 | 0 | Infiltrating | Mild | Intact | MSS | 1 | 1 | Liver | 1 | 0 | 0 | 2068 | 2830 | 8.472863235 | 2 |
| 1 | 1 | Infiltrating | Mild | Intact | MSS | 1 | 1 | Ovaries, Perito | 1 | 0 | 0 | 67 | 618 | 6.052045168 | 2 |
| 0 | 1 | Infiltrating | Absent | Intact | MSS | 1 | 1 | Liver, Lung | 1 | 0 | 0 | 147 | 278 | 6.052045168 | 2 |
| 1 | 1 | Infiltrating | Mild | Intact | MSS | 1 | 1 | Anastomosis, L | 0 | 0 | 1 | 91 | 1612 | 9.683272268 | 2 |
| 1 | 0 | Pushing | Mild | Intact | MSS | 0 | 0 | NA | 0 | 1 | 1 | 1792 | 3309 | 14.5249084 | 2 |
| 1 | 0 | Infiltrating | Absent | LH1/PMS2 abs | MSI-H | 0 | 0 | NA | 0 | 1 | 1 | 780 | 790 | 52.04758844 | 2 |
| 0 | 1 | Infiltrating | Marked | LH1/PMS2 abs | MSI-H | 0 | 0 | NA | 0 | 1 | 1 | 2872 | 3035 | 45.99554327 | 2 |
| 0 | 0 | Infiltrating | Moderate | Intact | MSS | 1 | 0 | NA | 1 | 1 | 0 | 98 | 109 | 10.61945963 | 1 |
| 1 | 0 | Infiltrating | Mild | Intact | MSS | 1 | 1 | Lung | 0 | 0 | 1 | 412 | 1311 | 12.10409034 | 2 |
| 1 | 1 | Infiltrating | Mild | Intact | MSS | 0 | 1 | RP LN | 0 | 0 | 1 | 529 | 2233 | 8.472863235 | 2 |
| 0 | 0 | Infiltrating | Moderate | Intact | MSS | 0 | 1 | Lung | 0 | 0 | 1 | 208 | 2024 | 9.683272268 | 2 |
| 0 | 0 | Infiltrating | Mild | Intact | MSS | 0 | 1 | Lung | 0 | 0 | 1 | 314 | 2121 | 6.052045168 | 2 |
| 1 | 0 | Infiltrating | Absent | Intact | MSS | 1 | 1 | Lung, Bone, Liv | 1 | 0 | 0 | 76 | 918 | 14.5249084 | 2 |
| 0 | 0 | Pushing | Mild | LH1/PMS2 Abs | MSI-H | 0 | 0 | NA | 1 | 1 | 0 | 337 | 337 | 44.78513424 | 2 |
| 1 | 1 | Infiltrating | Mild | Intact | MSS | 0 | 1 | RP LN | 0 | 0 | 1 | 1115 | 1879 | 2.420818067 | 2 |
| 0 | 1 | Infiltrating | Absent | Intact | MSS | 0 | 1 | Lung | 0 | 0 | 1 | 653 | 1819 | 9.683272268 | 2 |
| 0 | 0 | Pushing | Marked | Intact | MSS | 0 | 1 | Peritoneum, Ar | 1 | 0 | 0 | 87 | 291 | 12.10409034 | 2 |
| 0 | 0 | Infiltrating | Absent | Intact | MSS | 0 | 0 | NA | 1 | 1 | 0 | 777 | 1295 | 10.8936813 | 2 |
| 0 | 0 | Infiltrating | Absent | Intact | MSS | 0 | 0 | NA | 1 | 1 | 0 | 84 | 86 | 6.052045168 | 2 |
| 0 | 1 | Infiltrating | Absent | Intact | MSS | 1 | 1 | Lungs, LN, boc | 1 | 0 | 0 | 181 | 205 | 16.94572647 | 2 |
| 0 | 0 | Infiltrating | Mild | Intact | MSS | 0 | 0 | NA | 1 | 1 | 0 | 704 | 707 | 6.083289356 | 3 |
| 1 | 0 | Infiltrating | Mild | Intact | MSS | 0 | 1 | LN, Peritoneum | 0 | 0 | 1 | 1481 | 1532 | 8.364522865 | 3 |
| 0 | 1 | Infiltrating | Mild | Intact | MSS | 0 | 1 | Lung, LN, Bone | 1 | 0 | 0 | 311 | 1064 | 13.31449937 | 2 |
| 0 | 0 | Pushing | Moderate | LH1/PMS2 Abs | MSI-H | 0 | 0 | NA | 0 | 1 | 1 | 1757 | 3498 | 26.62899874 | 2 |
| 0 | 0 | Infiltrating | Moderate | LH1/PMS2 Abs | MSI-H | 0 | 0 | NA | 0 | 1 | 1 | 3000 | 3306 | 64.15167878 | 2 |
| 0 | 0 | Infiltrating | Moderate | PMS2 Absent | MSI-H | 0 | 0 | NA | 0 | 1 | 1 | 1741 | 3224 | 68.99331491 | 2 |
| 0 | 0 | Infiltrating | Mild | LH1/PMS2 Abs | MSI-H | 0 | 0 | NA | 0 | 1 | 1 | 19 | 19 | 59.31004264 | 2 |
| 0 | 0 | NA | Moderate | Intact | MSI-H | 0 | 0 | NA | 0 | 1 | 1 | 2226 | 3202 | 48.41636134 | 2 |
| 0 | 1 | Pushing | Marked | LH1/PMS2 Abs | MSI-H | 0 | 0 | NA | 0 | 1 | 1 | 1875 | 3634 | 54.46840651 | 2 |
| 0 | 0 | Infiltrating | Marked | Intact | MSS | 0 | 0 | NA | 1 | 0 | 0 | 2318 | 3079 | 10.8936813 | 2 |
| 0 | 1 | Infiltrating | Mild | Intact | MSS | 0 | 0 | NA | 0 | 1 | 1 | 204 | 3014 | 3.631227101 | 2 |
| 1 | 0 | Infiltrating | Absent | Intact | MSS | 0 | 0 | NA | 0 | 1 | 1 | 2118 | 3755 | 9.292027175 | 1 |
| 0 | 0 | Infiltrating | Mild | Intact | MSS | 0 | 0 | NA | 0 | 1 | 1 | 1840 | 3143 | 12.10409034 | 2 |
| 0 | 0 | Infiltrating | Mild | Intact | MSS | 0 | 0 | NA | 0 | 1 | 1 | 2941 | 3079 | 7.262454201 | 2 |
| 0 | 0 | Infiltrating | Absent | Intact | MSS | 0 | 0 | NA | 0 | 1 | 1 | 2131 | 2133 | 10.8936813 | 2 |
| 0 | 0 | Infiltrating | Moderate | Intact | MSS | 0 | 0 | NA | 0 | 1 | 1 | 2459 | 2975 | 8.472863235 | 2 |
| 0 | 0 | Infiltrating | Mild | Intact | MSS | 0 | 0 | NA | 0 | 1 | 1 | 3008 | 3114 | 10.8936813 | 2 |
| 1 | 0 | Infiltrating | Absent | Intact | MSS | 0 | 0 | NA | 0 | 1 | 1 | 2876 | 2909 | 12.10409034 | 2 |
| 0 | 0 | Pushing | Absent | Intact | MSS | 0 | 0 | NA | 1 | 1 | 0 | 1526 | 2572 | 9.683272268 | 2 |
| 0 | 0 | Infiltrating | Mild | Intact | MSS | 0 | 0 | NA | 0 | 1 | 1 | 1658 | 1660 | 7.262454201 | 2 |
| 0 | 1 | Infiltrating | Mild | Intact | MSS | 0 | 0 | NA | 0 | 1 | 1 | 3531 | 3650 | 7.964594722 | 1 |
| 0 | 0 | Infiltrating | Moderate | Intact | MSS | 0 | 0 | NA | 0 | 1 | 1 | 3002 | 3002 | 13.31449937 | 2 |
| 0 | 0 | Infiltrating | Absent | Intact | MSS | 0 | 0 | NA | 0 | 1 | 1 | 1558 | 3012 | 8.472863235 | 2 |
| 0 | 0 | Infiltrating | Mild | Intact | MSS | 0 | 0 | NA | 0 | 1 | 1 | 1558 | 1571 | 7.262454201 | 2 |
| 0 | 0 | Infiltrating | Absent | Intact | MSS | 0 | 0 | NA | 0 | 1 | 1 | 2718 | 2760 | 15.73531744 | 2 |
| 0 | 0 | Infiltrating | Mild | Intact | MSS | 0 | 0 | NA | 0 | 1 | 1 | 1737 | 2783 | 9.683272268 | 2 |
| 0 | 0 | Infiltrating | Mild | Intact | MSS | 0 | 0 | NA | 0 | 1 | 1 | 2182 | 2791 | 3.631227101 | 2 |
| 0 | 0 | Infiltrating | Absent | Intact | MSS | 0 | 0 | NA | 0 | 1 | 1 | 2374 | 2891 | 7.262454201 | 2 |
| 0 | 0 | Infiltrating | Absent | Intact | MSS | 0 | 0 | NA | 0 | 1 | 1 | 2605 | 2605 | 10.8936813 | 2 |
| 0 | 0 | Pushing | Mild | Intact | MSS | 0 | 0 | NA | 0 | 1 | 1 | 2614 | 2621 | 8.472863235 | 2 |
| 0 | 0 | Pushing | Mild | Intact | MSS | 0 | 0 | NA | 0 | 1 | 1 | 251 | 251 | 6.052045168 | 2 |
| 0 | 0 | Pushing | Moderate | Intact | MSS | 0 | 0 | NA | 0 | 1 | 1 | 1793 | 3648 | 7.262454201 | 2 |
| 0 | 0 | Infiltrating | Moderate | Intact | MSS | 0 | 0 | NA | 0 | 1 | 1 | 1119 | 2761 | 5.322878187 | 3 |
| 0 | 0 | Infiltrating | Mild | Intact | MSS | 0 | 0 | NA | 0 | 1 | 1 | 1478 | 2825 | 7.604111695 | 3 |
| 0 | NA | Infiltrating | Moderate | Intact | MSS | 0 | 1 | Liver | 0 | 0 | 1 | 893 | 4436 | 9.885345204 | 3 |
| 0 | 0 | NA | Absent | Intact | MSS | 0 | 1 | NA | 1 | 0 | 0 | 2199 | 3810 | 9.124934034 | 3 |
| 1 | NA | Infiltrating | Mild | LH1/PMS2 abs | MSI-H | 1 | 1 | NA | 1 | 0 | 0 | 18 | 66 | 83.64522865 | 3 |
| 0 | 0 | Pushing | Mild | Intact | MSS | 0 | 1 | Liver | 1 | 0 | 0 | 179 | 271 | 7.604111695 | 3 |
| 0 | NA | Infiltrating | Mild | Intact | MSS | 1 | 1 | Liver | 0 | 0 | 1 | 568 | 1963 | 10.64575637 | 3 |
| 1 | 0 | Infiltrating | Absent | Intact | MSS | 0 | 1 | NA | 1 | 0 | 0 | 693 | 1905 | 4.562467017 | 3 |
| 0 | 0 | NA | NA | Intact | MSS | 1 | 1 | NA | 0 | 1 | 1 | 1588 | 1943 | 13.68740105 | 3 |
| 1 | 0 | NA | NA | Intact | MSS | 0 | 1 | Brain | 1 | 0 | 0 | 228 | 1238 | 4.562467017 | 3 |
| 0 | 0 | NA | NA | Intact | MSS | 0 | 1 | NA | 0 | 0 | 1 | 697 | 1555 | 13.68740105 | 3 |
| 1 | 0 | NA | NA | Intact | MSS | 1 | 1 | NA | 1 | 0 | 0 | 308 | 970 | 5.322878187 | 3 |
| 1 | 1 | NA | NA | Intact | MSS | 1 | 1 | NA | 1 | 0 | 0 | 286 | 642 | 6.083289356 | 3 |

| KRAS | TP53 | NRAS | BRAF | PIK3CA | APC | FBXW7 | PTEN | TCF7L2 | SOX9 | CTNNB1 | Slide_ID | HTAN Particip | HTAN Parent I | HTAN Biospecimen ID** |
|---|---|---|---|---|---|---|---|---|---|---|---|---|---|---|
| 1 | 0 | 0 | 0 | 0 | 1 | 0 | 0 | 0 | 1 | 0 | LSP10353 | HTA7_926 | HTA7_926_1 | HTA7_926_8 |
| 1 | 0 | 0 | 0 | 0 | 1 | 0 | 0 | 0 | 1 | 0 | LSP10364 | HTA7_927 | HTA7_927_1 | HTA7_927_9 |
| 0 | 1 | 0 | 0 | 0 | 1 | 0 | 0 | 0 | 0 | 0 | LSP10375 | HTA7_932 | HTA7_932_1 | HTA7_932_9 |
| 0 | 1 | 0 | 1 | 0 | 0 | 0 | 0 | 0 | 0 | 0 | LSP10388 | HTA7_934 | HTA7_934_1 | HTA7_934_9 |
| 0 | 1 | 0 | 0 | 0 | 1 | 0 | 0 | 0 | 0 | 0 | LSP10397 | HTA7_966 | HTA7_966_1 | HTA7_966_7 |
| 0 | 1 | 0 | 1 | 0 | 0 | 0 | 0 | 0 | 0 | 0 | LSP10408 | HTA7_947 | HTA7_947_1 | HTA7_947_9 |
| 1 | 0 | 0 | 0 | 1 | 1 | 0 | 0 | 0 | 0 | 0 | LSP10419 | HTA7_989 | HTA7_989_1 | HTA7_989_9 |
| 1 | 1 | 0 | 0 | 0 | 1 | 0 | 0 | 0 | 0 | 0 | LSP10431 | HTA7_920 | HTA7_920_1 | HTA7_920_9 |
| 0 | 1 | 0 | 0 | 0 | 1 | 0 | 1 | 0 | 0 | 0 | LSP10441 | HTA7_922 | HTA7_922_1 | HTA7_922_9 |
| 0 | 1 | 0 | 0 | 0 | 1 | 0 | 0 | 0 | 0 | 0 | LSP10452 | HTA7_931 | HTA7_931_1 | HTA7_931_9 |
| 0 | 0 | 0 | 0 | 0 | 0 | 0 | 0 | 0 | 0 | 0 | LSP10463 | HTA7_940 | HTA7_940_1 | HTA7_940_9 |
| 1 | 0 | 0 | 0 | 0 | 1 | 0 | 0 | 0 | 0 | 0 | LSP10474 | HTA7_963 | HTA7_963_1 | HTA7_963_8 |
| 0 | 1 | 0 | 0 | 1 | 1 | 0 | 0 | 0 | 0 | 0 | LSP10485 | HTA7_972 | HTA7_972_1 | HTA7_972_9 |
| 1 | 0 | 0 | 0 | 1 | 1 | 0 | 0 | 0 | 1 | 0 | LSP10496 | HTA7_925 | HTA7_925_1 | HTA7_925_9 |
| 0 | 0 | 0 | 0 | 1 | 0 | 0 | 1 | 0 | 0 | 1 | LSP10507 | HTA7_982 | HTA7_982_1 | HTA7_982_9 |
| 0 | 0 | 0 | 1 | 0 | 1 | 0 | 0 | 0 | 1 | 0 | LSP10518 | HTA7_965 | HTA7_965_1 | HTA7_965_9 |
| 0 | 1 | 0 | 0 | 0 | 0 | 0 | 0 | 1 | 0 | 0 | LSP10529 | HTA7_904 | HTA7_904_1 | HTA7_904_6 |
| 1 | 1 | 0 | 0 | 0 | 1 | 0 | 0 | 0 | 0 | 0 | LSP10540 | HTA7_906 | HTA7_906_1 | HTA7_906_6 |
| 1 | 1 | 0 | 0 | 0 | 0 | 0 | 0 | 0 | 0 | 0 | LSP10551 | HTA7_909 | HTA7_909_1 | HTA7_909_6 |
| 1 | 0 | 0 | 0 | 0 | 1 | 0 | 0 | 0 | 0 | 0 | LSP10562 | HTA7_911 | HTA7_911_1 | HTA7_911_6 |
| 0 | 1 | 0 | 0 | 0 | 1 | 0 | 0 | 0 | 0 | 0 | LSP10573 | HTA7_914 | HTA7_914_1 | HTA7_914_6 |
| 1 | 1 | 0 | 0 | 1 | 1 | 1 | 0 | 0 | 0 | 0 | LSP10584 | HTA7_923 | HTA7_923_1 | HTA7_923_6 |
| 1 | 0 | 0 | 0 | 0 | 0 | 0 | 0 | 0 | 0 | 0 | LSP10595 | HTA7_924 | HTA7_924_1 | HTA7_924_6 |
| 0 | 1 | 0 | 0 | 0 | 0 | 0 | 0 | 0 | 0 | 0 | LSP10606 | HTA7_933 | HTA7_933_1 | HTA7_933_6 |
| 0 | 1 | 1 | 0 | 0 | 1 | 0 | 0 | 0 | 0 | 0 | LSP10617 | HTA7_938 | HTA7_938_1 | HTA7_938_6 |
| 0 | 1 | 0 | 1 | 0 | 0 | 0 | 0 | 0 | 0 | 0 | LSP10628 | HTA7_943 | HTA7_943_1 | HTA7_943_6 |
| 0 | 1 | 0 | 0 | 0 | 1 | 1 | 0 | 0 | 0 | 0 | LSP10639 | HTA7_954 | HTA7_954_1 | HTA7_954_6 |
| 1 | 1 | 0 | 0 | 0 | 0 | 1 | 0 | 1 | 0 | 0 | LSP10650 | HTA7_957 | HTA7_957_1 | HTA7_957_6 |
| 0 | 1 | 0 | 1 | 0 | 1 | 0 | 1 | 0 | 1 | 0 | LSP10661 | HTA7_968 | HTA7_968_1 | HTA7_968_6 |
| 1 | 0 | 0 | 0 | 0 | 0 | 0 | 0 | 0 | 0 | 0 | LSP10672 | HTA7_983 | HTA7_983_1 | HTA7_983_6 |
| 0 | 1 | 0 | 0 | 0 | 1 | 0 | 0 | 0 | 0 | 0 | LSP10683 | HTA7_985 | HTA7_985_1 | HTA7_985_6 |
| 0 | 1 | 0 | 0 | 0 | 1 | 0 | 0 | 0 | 0 | 0 | LSP10696 | HTA7_994 | HTA7_994_1 | HTA7_994_6 |
| 0 | 0 | 0 | 0 | 0 | 1 | 0 | 0 | 0 | 0 | 0 | LSP10705 | HTA7_919 | HTA7_919_1 | HTA7_919_6 |
| 0 | 1 | 0 | 1 | 0 | 1 | 0 | 0 | 0 | 0 | 0 | LSP10716 | HTA7_928 | HTA7_928_1 | HTA7_928_6 |
| 0 | 1 | 0 | 1 | 0 | 1 | 0 | 0 | 0 | 0 | 0 | LSP10727 | HTA7_941 | HTA7_941_1 | HTA7_941_6 |
| 0 | 0 | 0 | 1 | 0 | 0 | 0 | 0 | 0 | 0 | 0 | LSP10738 | HTA7_945 | HTA7_945_1 | HTA7_945_6 |
| 1 | 0 | 0 | 0 | 1 | 1 | 1 | 0 | 0 | 0 | 0 | LSP10749 | HTA7_936 | HTA7_936_1 | HTA7_936_6 |
| 0 | 0 | 0 | 0 | 0 | 0 | 0 | 0 | 0 | 1 | 0 | LSP10760 | HTA7_908 | HTA7_908_1 | HTA7_908_6 |
| 1 | 1 | 0 | 0 | 0 | 1 | 0 | 0 | 0 | 0 | 0 | LSP10771 | HTA7_917 | HTA7_917_1 | HTA7_917_6 |
| 0 | 1 | 0 | 1 | 0 | 1 | 0 | 0 | 0 | 0 | 0 | LSP10786 | HTA7_960 | HTA7_960_1 | HTA7_960_6 |
| 1 | 1 | 0 | 0 | 0 | 1 | 1 | 0 | 0 | 0 | 0 | LSP14363 | HTA7_905 | HTA7_905_1 | HTA7_905_6 |
| 1 | 1 | 0 | 0 | 0 | 1 | 0 | 0 | 0 | 0 | 0 | LSP14373 | HTA7_942 | HTA7_942_1 | HTA7_942_6 |
| 0 | 1 | 0 | 0 | 0 | 0 | 0 | 0 | 0 | 0 | 1 | LSP14383 | HTA7_949 | HTA7_949_1 | HTA7_949_6 |
| 0 | 0 | 0 | 1 | 1 | 1 | 0 | 1 | 0 | 0 | 0 | LSP14388 | HTA7_950 | HTA7_950_1 | HTA7_950_6 |
| 0 | 0 | 0 | 0 | 0 | 1 | 0 | 0 | 0 | 0 | 0 | LSP14393 | HTA7_951 | HTA7_951_1 | HTA7_951_6 |
| 0 | 1 | 1 | 0 | 0 | 1 | 0 | 0 | 0 | 1 | 0 | LSP14398 | HTA7_952 | HTA7_952_1 | HTA7_952_6 |
| 0 | 1 | 0 | 0 | 0 | 0 | 0 | 1 | 0 | 0 | 0 | LSP14403 | HTA7_953 | HTA7_953_1 | HTA7_953_6 |
| 0 | 1 | 0 | 0 | 0 | 1 | 0 | 0 | 0 | 0 | 0 | LSP14408 | HTA7_955 | HTA7_955_1 | HTA7_955_6 |
| 0 | 1 | 0 | 0 | 0 | 1 | 0 | 0 | 0 | 0 | 0 | LSP14413 | HTA7_959 | HTA7_959_1 | HTA7_959_6 |
| 0 | 1 | 0 | 1 | 0 | 0 | 0 | 0 | 0 | 0 | 0 | LSP14418 | HTA7_907 | HTA7_907_1 | HTA7_907_6 |
| 1 | 1 | 0 | 0 | 0 | 1 | 0 | 0 | 0 | 0 | 0 | LSP14423 | HTA7_961 | HTA7_961_1 | HTA7_961_6 |
| 0 | 1 | 0 | 0 | 0 | 1 | 0 | 0 | 0 | 0 | 0 | LSP14438 | HTA7_969 | HTA7_969_1 | HTA7_969_6 |
| 0 | 1 | 0 | 0 | 1 | 1 | 0 | 0 | 0 | 0 | 0 | LSP14443 | HTA7_971 | HTA7_971_1 | HTA7_971_6 |
| 1 | 1 | 0 | 0 | 0 | 1 | 0 | 0 | 0 | 0 | 0 | LSP14448 | HTA7_975 | HTA7_975_1 | HTA7_975_6 |
| 0 | 1 | 1 | 0 | 0 | 1 | 0 | 0 | 0 | 1 | 0 | LSP14453 | HTA7_976 | HTA7_976_1 | HTA7_976_6 |
| 1 | 0 | 0 | 0 | 0 | 1 | 0 | 0 | 0 | 0 | 0 | LSP14458 | HTA7_977 | HTA7_977_1 | HTA7_977_6 |
| 1 | 0 | 0 | 0 | 0 | 1 | 0 | 0 | 0 | 0 | 0 | LSP14463 | HTA7_978 | HTA7_978_1 | HTA7_978_6 |
| 1 | 0 | 0 | 0 | 0 | 1 | 1 | 0 | 0 | 0 | 0 | LSP14468 | HTA7_979 | HTA7_979_1 | HTA7_979_6 |
| 0 | 1 | 0 | 0 | 0 | 1 | 0 | 0 | 0 | 0 | 0 | LSP14473 | HTA7_980 | HTA7_980_1 | HTA7_980_6 |
| 0 | 1 | 0 | 0 | 0 | 1 | 0 | 0 | 0 | 0 | 0 | LSP14483 | HTA7_984 | HTA7_984_1 | HTA7_984_6 |
| 0 | 0 | 0 | 0 | 0 | 1 | 0 | 0 | 0 | 0 | 0 | LSP14493 | HTA7_910 | HTA7_910_1 | HTA7_910_5 |
| 0 | 0 | 1 | 1 | 0 | 0 | 0 | 0 | 0 | 0 | 0 | LSP14498 | HTA7_991 | HTA7_991_1 | HTA7_991_6 |
| 0 | 1 | 0 | 0 | 0 | 1 | 0 | 0 | 0 | 0 | 0 | LSP14503 | HTA7_992 | HTA7_992_1 | HTA7_992_6 |
| 1 | 0 | 0 | 0 | 1 | 1 | 0 | 0 | 0 | 0 | 0 | LSP15280 | HTA7_996 | HTA7_996_1 | HTA7_996_2 |
| 0 | 1 | 0 | 0 | 0 | 1 | 0 | 0 | 0 | 0 | 0 | LSP15284 | HTA7_997 | HTA7_997_1 | HTA7_997_2 |
| 0 | 0 | 0 | 1 | 0 | 0 | 0 | 0 | 0 | 0 | 0 | LSP15288 | HTA7_998 | HTA7_998_1 | HTA7_998_2 |
| 1 | 1 | 0 | 0 | 0 | 1 | 0 | 0 | 0 | 0 | 0 | LSP15292 | HTA7_999 | HTA7_999_1 | HTA7_999_2 |
| 1 | 0 | 0 | 0 | 0 | 1 | 0 | 0 | 0 | 0 | 0 | LSP15300 | HTA7_1000 | HTA7_1000_1 | HTA7_1000_2 |
| 1 | 1 | 0 | 0 | 0 | 1 | 0 | 0 | 0 | 0 | 0 | LSP15304 | HTA7_1001 | HTA7_1001_1 | HTA7_1001_2 |
| 0 | 1 | 0 | 0 | 1 | 1 | 0 | 0 | 0 | 1 | 0 | LSP15308 | HTA7_1002 | HTA7_1002_1 | HTA7_1002_2 |
| 1 | 1 | 0 | 0 | 1 | 1 | 0 | 0 | 0 | 0 | 0 | LSP15312 | HTA7_1003 | HTA7_1003_1 | HTA7_1003_2 |
| 1 | 1 | 0 | 0 | 0 | 1 | 0 | 0 | 0 | 0 | 0 | LSP15316 | HTA7_1004 | HTA7_1004_1 | HTA7_1004_2 |
| 1 | 0 | 0 | 0 | 0 | 1 | 0 | 0 | 0 | 0 | 0 | LSP15320 | HTA7_1005 | HTA7_1005_1 | HTA7_1005_2 |
| 1 | 1 | 0 | 0 | 0 | 1 | 0 | 0 | 0 | 1 | 0 | LSP15324 | HTA7_1006 | HTA7_1006_1 | HTA7_1006_2 |

**Supplementary Table 2**: *_N: Number of dMMR/tdpMMR/tipMMR tumors with mutation; *_fraction: Fraction of dMMR/tdpMMR/tipMMR tumors with mutation; pvalue_3way: p-value (Fisher's exact test) for enrichment in mutation frequency across all three T cell/MMR groups; pvalue_tip_vs_tdp: p-value (Fisher's exact test) for enrichment in mutation frequency between tdpMMR versus tipMMR tumors; pvalue_pmmr_vs_dmmr: p-value (Fisher's exact test) for enrichment in mutation frequency between all pMMR versus dMMR tumors; qvalue_*: Corresponding q-value after false discovery rate (FDR) correction; gene: Hugo gene identifier. Mutation = any [illegible]

| dMMR_N | dMMR_fractior | tdpMMR_N | tdpMMR_fraction | tipMMR_N | tipMMR_fraction | pvalue_3way | value_tip_vs_tc | e_pmmr_vs_c | gene | qvalue_3way | qvalue_tip_vs_tdp | e_pmmr_vs_dmmr |
|---|---|---|---|---|---|---|---|---|---|---|---|---|
| 1 | 0.1 | 0 | 0 | 1 | 0.071428571 | 0.102184376 | 0.21875 | 0.253609774 | ABL1 | 0.244220124 | 1 | 0.371136255 |
| 0 | 0 | 0 | 0 | 1 | 0.071428571 | 0.324324324 | 0.21875 | 1 | AKT1 | 0.43631075 | 1 | 1 |
| 0 | 0 | 1 | 0.02 | 0 | 0 | 1 | 1 | 1 | AKT3 | 1 | 1 | 1 |
| 3 | 0.3 | 2 | 0.04 | 0 | 0 | 0.020737407 | 1 | 0.015867884 | ALK | 0.109144249 | 1 | 0.090875433 |
| 1 | 0.1 | 1 | 0.02 | 0 | 0 | 0.287301 | 1 | 0.253609774 | ALOX12B | 0.408484834 | 1 | 0.371136255 |
| 6 | 0.6 | 41 | 0.82 | 10 | 0.714285714 | 0.231423443 | 0.456620522 | 0.224046783 | APC | 0.399005937 | 1 | 0.371136255 |
| 2 | 0.2 | 2 | 0.04 | 0 | 0 | 0.1450958 | 1 | 0.085701175 | AR | 0.256051412 | 1 | 0.254971953 |
| 2 | 0.2 | 0 | 0 | 0 | 0 | 0.016660496 | 1 | 0.016660496 | ARAF | 0.096118247 | 1 | 0.090875433 |
| 5 | 0.5 | 1 | 0.02 | 0 | 0 | 0.000104404 | 1 | 8.8194E-05 | ARID1A | 0.010440442 | 1 | 0.008819396 |
| 5 | 0.5 | 4 | 0.08 | 2 | 0.142857143 | 0.00654172 | 0.60436292 | 0.004921939 | ARID1B | 0.096118247 | 1 | 0.073469324 |
| 1 | 0.1 | 3 | 0.06 | 0 | 0 | 0.561369203 | 1 | 0.447799719 | ARID2 | 0.695912235 | 1 | 0.599731767 |
| 4 | 0.4 | 3 | 0.06 | 1 | 0.071428571 | 0.022100437 | 1 | 0.009578318 | ASXL1 | 0.114312603 | 1 | 0.090875433 |
| 5 | 0.5 | 4 | 0.08 | 2 | 0.142857143 | 0.00654172 | 0.60436292 | 0.004921939 | ATM | 0.096118247 | 1 | 0.073469324 |
| 1 | 0.1 | 0 | 0 | 0 | 0 | 0.135135135 | 1 | 0.135135135 | ATR | 0.244220124 | 1 | 0.254971953 |
| 0 | 0 | 2 | 0.04 | 0 | 0 | 1 | 1 | 1 | ATRX | 1 | 1 | 1 |
| 0 | 0 | 1 | 0.02 | 0 | 0 | 1 | 1 | 1 | AURKB | 1 | 1 | 1 |
| 0 | 0 | 2 | 0.04 | 0 | 0 | 1 | 1 | 1 | AXL | 1 | 1 | 1 |
| 3 | 0.3 | 0 | 0 | 0 | 0 | 0.001851166 | 1 | 0.001851166 | B2M | 0.042719221 | 1 | 0.042719221 |
| 0 | 0 | 1 | 0.02 | 0 | 0 | 1 | 1 | 1 | BAP1 | 1 | 1 | 1 |
| 0 | 0 | 1 | 0.02 | 0 | 0 | 1 | 1 | 1 | BCL11B | 1 | 1 | 1 |
| 0 | 0 | 1 | 0.02 | 0 | 0 | 1 | 1 | 1 | BCL6 | 1 | 1 | 1 |
| 2 | 0.2 | 0 | 0 | 0 | 0 | 0.016660496 | 1 | 0.016660496 | BCOR | 0.096118247 | 1 | 0.090875433 |
| 2 | 0.2 | 2 | 0.04 | 0 | 0 | 0.1450958 | 1 | 0.085701175 | BCORL1 | 0.256051412 | 1 | 0.254971953 |
| 4 | 0.4 | 0 | 0 | 1 | 0.071428571 | 0.000322433 | 0.21875 | 0.000849972 | BLM | 0.019345991 | 1 | 0.042498605 |
| 1 | 0.1 | 2 | 0.04 | 0 | 0 | 0.433080341 | 1 | 0.357275083 | BMPR1A | 0.560017682 | 1 | 0.491662959 |
| 5 | 0.5 | 6 | 0.12 | 4 | 0.285714286 | 0.015250212 | 0.205866524 | 0.024404812 | BRAF | 0.096118247 | 1 | 0.128446379 |
| 2 | 0.2 | 1 | 0.02 | 1 | 0.071428571 | 0.057643405 | 0.392361111 | 0.085701175 | BRCA1 | 0.244220124 | 1 | 0.254971953 |
| 4 | 0.4 | 2 | 0.04 | 2 | 0.142857143 | 0.003625879 | 0.205667195 | 0.009578318 | BRCA2 | 0.063986103 | 1 | 0.090875433 |
| 1 | 0.1 | 0 | 0 | 0 | 0 | 0.135135135 | 1 | 0.135135135 | BRD4 | 0.244220124 | 1 | 0.254971953 |
| 0 | 0 | 3 | 0.06 | 0 | 0 | 1 | 1 | 1 | BRIP1 | 1 | 1 | 1 |
| 0 | 0 | 0 | 0 | 1 | 0.071428571 | 0.324324324 | 0.21875 | 1 | BUB1B | 0.43631075 | 1 | 1 |
| 1 | 0.1 | 0 | 0 | 0 | 0 | 0.135135135 | 1 | 0.135135135 | CADM2 | 0.244220124 | 1 | 0.254971953 |
| 2 | 0.2 | 1 | 0.02 | 0 | 0 | 0.065932371 | 1 | 0.046279156 | CARD11 | 0.244220124 | 1 | 0.201213721 |
| 0 | 0 | 1 | 0.02 | 0 | 0 | 1 | 1 | 1 | CBFA2T3 | 1 | 1 | 1 |
| 1 | 0.1 | 1 | 0.02 | 0 | 0 | 0.287301 | 1 | 0.253609774 | CBL | 0.408484834 | 1 | 0.371136255 |
| 2 | 0.2 | 0 | 0 | 0 | 0 | 0.016660496 | 1 | 0.016660496 | CBLB | 0.096118247 | 1 | 0.090875433 |
| 1 | 0.1 | 0 | 0 | 0 | 0 | 0.135135135 | 1 | 0.135135135 | CCND2 | 0.244220124 | 1 | 0.254971953 |
| 1 | 0.1 | 0 | 0 | 0 | 0 | 0.135135135 | 1 | 0.135135135 | CCNE1 | 0.244220124 | 1 | 0.254971953 |
| 1 | 0.1 | 0 | 0 | 0 | 0 | 0.135135135 | 1 | 0.135135135 | CD274 | 0.244220124 | 1 | 0.254971953 |
| 1 | 0.1 | 0 | 0 | 0 | 0 | 0.135135135 | 1 | 0.135135135 | CD58 | 0.244220124 | 1 | 0.254971953 |
| 1 | 0.1 | 0 | 0 | 0 | 0 | 0.135135135 | 1 | 0.135135135 | CDC73 | 0.244220124 | 1 | 0.254971953 |
| 0 | 0 | 1 | 0.02 | 0 | 0 | 1 | 1 | 1 | CDH1 | 1 | 1 | 1 |
| 1 | 0.1 | 1 | 0.02 | 0 | 0 | 0.287301 | 1 | 0.253609774 | CDH4 | 0.408484834 | 1 | 0.371136255 |
| 1 | 0.1 | 0 | 0 | 0 | 0 | 0.135135135 | 1 | 0.135135135 | CDK12 | 0.244220124 | 1 | 0.254971953 |
| 1 | 0.1 | 0 | 0 | 1 | 0.071428571 | 0.102184376 | 0.21875 | 0.253609774 | CDK9 | 0.244220124 | 1 | 0.371136255 |
| 0 | 0 | 2 | 0.04 | 0 | 0 | 1 | 1 | 1 | CDKN1B | 1 | 1 | 1 |
| 1 | 0.1 | 0 | 0 | 0 | 0 | 0.135135135 | 1 | 0.135135135 | CDKN2B | 0.244220124 | 1 | 0.254971953 |
| 0 | 0 | 1 | 0.02 | 0 | 0 | 1 | 1 | 1 | CDKN2C | 1 | 1 | 1 |
| 1 | 0.1 | 2 | 0.04 | 0 | 0 | 0.433080341 | 1 | 0.357275083 | CEBPA | 0.560017682 | 1 | 0.491662959 |
| 1 | 0.1 | 1 | 0.02 | 0 | 0 | 0.287301 | 1 | 0.253609774 | CHEK2 | 0.408484834 | 1 | 0.371136255 |
| 1 | 0.1 | 0 | 0 | 0 | 0 | 0.135135135 | 1 | 0.135135135 | CIC | 0.244220124 | 1 | 0.254971953 |
| 3 | 0.3 | 2 | 0.04 | 0 | 0 | 0.020737407 | 1 | 0.015867884 | CIITA | 0.109144249 | 1 | 0.090875433 |
| 0 | 0 | 1 | 0.02 | 0 | 0 | 1 | 1 | 1 | COL7A1 | 1 | 1 | 1 |
| 5 | 0.5 | 4 | 0.08 | 1 | 0.071428571 | 0.005874061 | 1 | 0.002867312 | CREBBP | 0.096118247 | 1 | 0.053762103 |
| 1 | 0.1 | 0 | 0 | 0 | 0 | 0.135135135 | 1 | 0.135135135 | CRTC1 | 0.244220124 | 1 | 0.254971953 |
| 2 | 0.2 | 0 | 0 | 0 | 0 | 0.016660496 | 1 | 0.016660496 | CRTC2 | 0.096118247 | 1 | 0.090875433 |
| 0 | 0 | 0 | 0 | 1 | 0.071428571 | 0.324324324 | 0.21875 | 1 | CRTC3 | 0.43631075 | 1 | 1 |
| 1 | 0.1 | 0 | 0 | 1 | 0.071428571 | 0.102184376 | 0.21875 | 0.253609774 | CSF1R | 0.244220124 | 1 | 0.371136255 |
| 0 | 0 | 1 | 0.02 | 0 | 0 | 1 | 1 | 1 | CSF3R | 1 | 1 | 1 |
| 1 | 0.1 | 0 | 0 | 0 | 0 | 0.135135135 | 1 | 0.135135135 | CTCF | 0.244220124 | 1 | 0.254971953 |
| 2 | 0.2 | 2 | 0.04 | 1 | 0.071428571 | 0.120335365 | 0.529569892 | 0.132256702 | CTNNB1 | 0.244220124 | 1 | 0.254971953 |
| 0 | 0 | 3 | 0.06 | 0 | 0 | 1 | 1 | 1 | CUX1 | 1 | 1 | 1 |
| 2 | 0.2 | 0 | 0 | 1 | 0.071428571 | 0.017184993 | 0.21875 | 0.046279156 | CYLD | 0.097273546 | 1 | 0.201213721 |
| 1 | 0.1 | 1 | 0.02 | 0 | 0 | 0.287301 | 1 | 0.253609774 | DDB1 | 0.408484834 | 1 | 0.371136255 |
| 1 | 0.1 | 0 | 0 | 0 | 0 | 0.135135135 | 1 | 0.135135135 | DDB2 | 0.244220124 | 1 | 0.254971953 |
| 1 | 0.1 | 0 | 0 | 0 | 0 | 0.135135135 | 1 | 0.135135135 | DDR2 | 0.244220124 | 1 | 0.254971953 |
| 1 | 0.1 | 0 | 0 | 0 | 0 | 0.135135135 | 1 | 0.135135135 | DEPDC5 | 0.244220124 | 1 | 0.254971953 |
| 3 | 0.3 | 1 | 0.02 | 0 | 0 | 0.014449526 | 1 | 0.006857137 | DICER1 | 0.096118247 | 1 | 0.073469324 |
| 1 | 0.1 | 0 | 0 | 0 | 0 | 0.135135135 | 1 | 0.135135135 | DIS3 | 0.244220124 | 1 | 0.254971953 |
| 0 | 0 | 1 | 0.02 | 0 | 0 | 1 | 1 | 1 | DIS3L2 | 1 | 1 | 1 |
| 1 | 0.1 | 1 | 0.02 | 0 | 0 | 0.287301 | 1 | 0.253609774 | DMD | 0.408484834 | 1 | 0.371136255 |
| 3 | 0.3 | 2 | 0.04 | 0 | 0 | 0.020737407 | 1 | 0.015867884 | DNMT3A | 0.109144249 | 1 | 0.090875433 |
| 0 | 0 | 0 | 0 | 1 | 0.071428571 | 0.324324324 | 0.21875 | 1 | DOCK8 | 0.43631075 | 1 | 1 |
| 0 | 0 | 1 | 0.02 | 0 | 0 | 1 | 1 | 1 | EED | 1 | 1 | 1 |
| 4 | 0.4 | 0 | 0 | 1 | 0.071428571 | 0.000322433 | 0.21875 | 0.000849972 | EGFR | 0.019345991 | 1 | 0.042498605 |
| 1 | 0.1 | 0 | 0 | 0 | 0 | 0.135135135 | 1 | 0.135135135 | ELANE | 0.244220124 | 1 | 0.254971953 |
| 3 | 0.3 | 2 | 0.04 | 1 | 0.071428571 | 0.03131577 | 0.529569892 | 0.02936224 | EP300 | 0.159232729 | 1 | 0.149299524 |
| 2 | 0.2 | 2 | 0.04 | 1 | 0.071428571 | 0.120335365 | 0.529569892 | 0.132256702 | EPHA3 | 0.244220124 | 1 | 0.254971953 |
| 3 | 0.3 | 0 | 0 | 0 | 0 | 0.001851166 | 1 | 0.001851166 | EPHA5 | 0.042719221 | 1 | 0.042719221 |
| 1 | 0.1 | 0 | 0 | 1 | 0.071428571 | 0.102184376 | 0.21875 | 0.253609774 | EPHA7 | 0.244220124 | 1 | 0.371136255 |
| 1 | 0.1 | 2 | 0.04 | 0 | 0 | 0.433080341 | 1 | 0.357275083 | ERBB2 | 0.560017682 | 1 | 0.491662959 |
| 1 | 0.1 | 1 | 0.02 | 0 | 0 | 0.287301 | 1 | 0.253609774 | ERBB3 | 0.408484834 | 1 | 0.371136255 |
| 2 | 0.2 | 5 | 0.1 | 1 | 0.071428571 | 0.5357319 | 1 | 0.29414158 | ERBB4 | 0.681015128 | 1 | 0.426292145 |
| 0 | 0 | 1 | 0.02 | 0 | 0 | 1 | 1 | 1 | ERCC2 | 1 | 1 | 1 |
| 0 | 0 | 1 | 0.02 | 0 | 0 | 1 | 1 | 1 | ERCC3 | 1 | 1 | 1 |
| 1 | 0.1 | 1 | 0.02 | 0 | 0 | 0.287301 | 1 | 0.253609774 | ERCC4 | 0.408484834 | 1 | 0.371136255 |
| 2 | 0.2 | 1 | 0.02 | 0 | 0 | 0.065932371 | 1 | 0.046279156 | ERCC5 | 0.244220124 | 1 | 0.201213721 |
| 3 | 0.3 | 1 | 0.02 | 0 | 0 | 0.014449526 | 1 | 0.006857137 | ESR1 | 0.096118247 | 1 | 0.073469324 |
| 0 | 0 | 0 | 0 | 1 | 0.071428571 | 0.324324324 | 0.21875 | 1 | ETV1 | 0.43631075 | 1 | 1 |
| 1 | 0.1 | 0 | 0 | 0 | 0 | 0.135135135 | 1 | 0.135135135 | ETV4 | 0.244220124 | 1 | 0.254971953 |
| 1 | 0.1 | 0 | 0 | 0 | 0 | 0.135135135 | 1 | 0.135135135 | ETV5 | 0.244220124 | 1 | 0.254971953 |

| | | | | | | | | | | | | |
|---|---|---|---|---|---|---|---|---|---|---|---|---|
| 1 | 0.1 | 4 | 0.08 | 0 | 0 | 0.629506025 | 0.568129737 | 0.526685474 | ETV6 | 0.773982818 | 1 | 0.696060098 |
| 1 | 0.1 | 1 | 0.02 | 0 | 0 | 0.287301 | 1 | 0.253609774 | EWSR1 | 0.408484834 | 1 | 0.371136255 |
| 1 | 0.1 | 0 | 0 | 0 | 0 | 0.135135135 | 1 | 0.135135135 | EXT1 | 0.244220124 | 1 | 0.254971953 |
| 1 | 0.1 | 1 | 0.02 | 0 | 0 | 0.287301 | 1 | 0.253609774 | EXT2 | 0.408484834 | 1 | 0.371136255 |
| 1 | 0.1 | 0 | 0 | 0 | 0 | 0.135135135 | 1 | 0.135135135 | EZH2 | 0.244220124 | 1 | 0.254971953 |
| 0 | 0 | 1 | 0.02 | 0 | 0 | 1 | 1 | 1 | FAN1 | 1 | 1 | 1 |
| 1 | 0.1 | 0 | 0 | 1 | 0.071428571 | 0.102184376 | 0.21875 | 0.253609774 | FANCA | 0.244220124 | 1 | 0.371136255 |
| 1 | 0.1 | 2 | 0.04 | 0 | 0 | 0.433080341 | 1 | 0.357275083 | FANCC | 0.560017682 | 1 | 0.491662959 |
| 1 | 0.1 | 1 | 0.02 | 0 | 0 | 0.287301 | 1 | 0.253609774 | FANCD2 | 0.408484834 | 1 | 0.371136255 |
| 2 | 0.2 | 0 | 0 | 0 | 0 | 0.016660496 | 1 | 0.016660496 | FANCG | 0.096118247 | 1 | 0.090875433 |
| 1 | 0.1 | 0 | 0 | 0 | 0 | 0.135135135 | 1 | 0.135135135 | FANCM | 0.244220124 | 1 | 0.254971953 |
| 1 | 0.1 | 0 | 0 | 0 | 0 | 0.135135135 | 1 | 0.135135135 | FAS | 0.244220124 | 1 | 0.254971953 |
| 1 | 0.1 | 1 | 0.02 | 0 | 0 | 0.287301 | 1 | 0.253609774 | FAT1 | 0.408484834 | 1 | 0.371136255 |
| 2 | 0.2 | 5 | 0.1 | 1 | 0.071428571 | 0.5357319 | 1 | 0.29414158 | FBXW7 | 0.681015128 | 1 | 0.426292145 |
| 1 | 0.1 | 0 | 0 | 0 | 0 | 0.135135135 | 1 | 0.135135135 | FGFR2 | 0.244220124 | 1 | 0.254971953 |
| 0 | 0 | 1 | 0.02 | 1 | 0.071428571 | 0.546464272 | 0.392361111 | 1 | FGFR3 | 0.685938417 | 1 | 1 |
| 1 | 0.1 | 1 | 0.02 | 0 | 0 | 0.287301 | 1 | 0.253609774 | FGFR4 | 0.408484834 | 1 | 0.371136255 |
| 1 | 0.1 | 1 | 0.02 | 0 | 0 | 0.287301 | 1 | 0.253609774 | FH | 0.408484834 | 1 | 0.371136255 |
| 1 | 0.1 | 0 | 0 | 1 | 0.071428571 | 0.102184376 | 0.21875 | 0.253609774 | FKBP9 | 0.244220124 | 1 | 0.371136255 |
| 0 | 0 | 2 | 0.04 | 0 | 0 | 1 | 1 | 1 | FLCN | 1 | 1 | 1 |
| 1 | 0.1 | 3 | 0.06 | 0 | 0 | 0.561369203 | 1 | 0.447799719 | FLT1 | 0.695912235 | 1 | 0.599731767 |
| 2 | 0.2 | 0 | 0 | 0 | 0 | 0.016660496 | 1 | 0.016660496 | FLT3 | 0.096118247 | 1 | 0.090875433 |
| 1 | 0.1 | 4 | 0.08 | 0 | 0 | 0.629506025 | 0.568129737 | 0.526685474 | FLT4 | 0.773982818 | 1 | 0.696060098 |
| 2 | 0.2 | 0 | 0 | 0 | 0 | 0.016660496 | 1 | 0.016660496 | FUS | 0.096118247 | 1 | 0.090875433 |
| 0 | 0 | 0 | 0 | 1 | 0.071428571 | 0.324324324 | 0.21875 | 1 | GALNT12 | 0.43631075 | 1 | 1 |
| 1 | 0.1 | 0 | 0 | 0 | 0 | 0.135135135 | 1 | 0.135135135 | GATA3 | 0.244220124 | 1 | 0.254971953 |
| 2 | 0.2 | 0 | 0 | 0 | 0 | 0.016660496 | 1 | 0.016660496 | GATA4 | 0.096118247 | 1 | 0.090875433 |
| 1 | 0.1 | 0 | 0 | 0 | 0 | 0.135135135 | 1 | 0.135135135 | GATA6 | 0.244220124 | 1 | 0.254971953 |
| 1 | 0.1 | 0 | 0 | 0 | 0 | 0.135135135 | 1 | 0.135135135 | GBA | 0.244220124 | 1 | 0.254971953 |
| 0 | 0 | 0 | 0 | 1 | 0.071428571 | 0.324324324 | 0.21875 | 1 | GEN1 | 0.43631075 | 1 | 1 |
| 2 | 0.2 | 1 | 0.02 | 0 | 0 | 0.065932371 | 1 | 0.046279156 | GLI1 | 0.244220124 | 1 | 0.201213721 |
| 4 | 0.4 | 2 | 0.04 | 1 | 0.071428571 | 0.00746761 | 0.529569892 | 0.005151778 | GLI2 | 0.096118247 | 1 | 0.073469324 |
| 4 | 0.4 | 1 | 0.02 | 0 | 0 | 0.001595653 | 1 | 0.000849972 | GLI3 | 0.042719221 | 1 | 0.042498605 |
| 1 | 0.1 | 0 | 0 | 0 | 0 | 0.135135135 | 1 | 0.135135135 | GNA11 | 0.244220124 | 1 | 0.254971953 |
| 1 | 0.1 | 0 | 0 | 0 | 0 | 0.135135135 | 1 | 0.135135135 | GNAS | 0.244220124 | 1 | 0.254971953 |
| 1 | 0.1 | 0 | 0 | 0 | 0 | 0.135135135 | 1 | 0.135135135 | GNB2L1 | 0.244220124 | 1 | 0.254971953 |
| 0 | 0 | 1 | 0.02 | 0 | 0 | 1 | 1 | 1 | HABP2 | 1 | 1 | 1 |
| 1 | 0.1 | 0 | 0 | 0 | 0 | 0.135135135 | 1 | 0.135135135 | HELQ | 0.244220124 | 1 | 0.254971953 |
| 1 | 0.1 | 0 | 0 | 0 | 0 | 0.135135135 | 1 | 0.135135135 | HIST1H3C | 0.244220124 | 1 | 0.254971953 |
| 1 | 0.1 | 0 | 0 | 0 | 0 | 0.135135135 | 1 | 0.135135135 | HNF1A | 0.244220124 | 1 | 0.254971953 |
| 1 | 0.1 | 0 | 0 | 1 | 0.071428571 | 0.102184376 | 0.21875 | 0.253609774 | IGF1R | 0.244220124 | 1 | 0.371136255 |
| 0 | 0 | 1 | 0.02 | 0 | 0 | 1 | 1 | 1 | IGF2 | 1 | 1 | 1 |
| 2 | 0.2 | 0 | 0 | 0 | 0 | 0.016660496 | 1 | 0.016660496 | IKZF1 | 0.096118247 | 1 | 0.090875433 |
| 1 | 0.1 | 3 | 0.06 | 0 | 0 | 0.561369203 | 1 | 0.447799719 | IKZF3 | 0.695912235 | 1 | 0.599731767 |
| 3 | 0.3 | 0 | 0 | 0 | 0 | 0.001851166 | 1 | 0.001851166 | JAK2 | 0.042719221 | 1 | 0.042719221 |
| 0 | 0 | 1 | 0.02 | 0 | 0 | 1 | 1 | 1 | JAK3 | 1 | 1 | 1 |
| 0 | 0 | 2 | 0.04 | 0 | 0 | 1 | 1 | 1 | KAT6A | 1 | 1 | 1 |
| 0 | 0 | 1 | 0.02 | 0 | 0 | 1 | 1 | 1 | KAT6B | 1 | 1 | 1 |
| 0 | 0 | 1 | 0.02 | 0 | 0 | 1 | 1 | 1 | KDM5A | 1 | 1 | 1 |
| 1 | 0.1 | 1 | 0.02 | 1 | 0.071428571 | 0.244107121 | 0.392361111 | 0.357275083 | KDM5C | 0.408484834 | 1 | 0.491662959 |
| 0 | 0 | 0 | 0 | 1 | 0.071428571 | 0.324324324 | 0.21875 | 1 | KDM6A | 0.43631075 | 1 | 1 |
| 2 | 0.2 | 1 | 0.02 | 1 | 0.071428571 | 0.057643405 | 0.392361111 | 0.085701175 | KDM6B | 0.244220124 | 1 | 0.254971953 |
| 2 | 0.2 | 0 | 0 | 0 | 0 | 0.016660496 | 1 | 0.016660496 | KDR | 0.096118247 | 1 | 0.090875433 |
| 2 | 0.2 | 2 | 0.04 | 1 | 0.071428571 | 0.120335365 | 0.529569892 | 0.132256702 | KEAP1 | 0.244220124 | 1 | 0.254971953 |
| 1 | 0.1 | 0 | 0 | 0 | 0 | 0.135135135 | 1 | 0.135135135 | KIF1B | 0.244220124 | 1 | 0.254971953 |
| 2 | 0.2 | 1 | 0.02 | 0 | 0 | 0.065932371 | 1 | 0.046279156 | KIT | 0.244220124 | 1 | 0.201213721 |
| 1 | 0.1 | 0 | 0 | 0 | 0 | 0.135135135 | 1 | 0.135135135 | KLF4 | 0.244220124 | 1 | 0.254971953 |
| 1 | 0.1 | 1 | 0.02 | 1 | 0.071428571 | 0.244107121 | 0.392361111 | 0.357275083 | KMT2A | 0.408484834 | 1 | 0.491662959 |
| 7 | 0.7 | 2 | 0.04 | 0 | 0 | 3.19642E-06 | 1 | 2.21499E-06 | KMT2D | 0.000479464 | 1 | 0.000332249 |
| 2 | 0.2 | 25 | 0.5 | 3 | 0.214285714 | 0.062917015 | 0.072233128 | 0.186529271 | KRAS | 0.244220124 | 1 | 0.345424577 |
| 0 | 0 | 1 | 0.02 | 0 | 0 | 1 | 1 | 1 | LMO2 | 1 | 1 | 1 |
| 0 | 0 | 2 | 0.04 | 0 | 0 | 1 | 1 | 1 | MAP2K1 | 1 | 1 | 1 |
| 1 | 0.1 | 2 | 0.04 | 1 | 0.071428571 | 0.391027145 | 0.529569892 | 0.447799719 | MAP2K4 | 0.519062582 | 1 | 0.599731767 |
| 2 | 0.2 | 2 | 0.04 | 0 | 0 | 0.1450958 | 1 | 0.085701175 | MAP3K1 | 0.256051412 | 1 | 0.254971953 |
| 1 | 0.1 | 0 | 0 | 0 | 0 | 0.135135135 | 1 | 0.135135135 | MCL1 | 0.244220124 | 1 | 0.254971953 |
| 0 | 0 | 1 | 0.02 | 0 | 0 | 1 | 1 | 1 | MCM8 | 1 | 1 | 1 |
| 1 | 0.1 | 0 | 0 | 0 | 0 | 0.135135135 | 1 | 0.135135135 | MDM2 | 0.244220124 | 1 | 0.254971953 |
| 1 | 0.1 | 0 | 0 | 0 | 0 | 0.135135135 | 1 | 0.135135135 | MECOM | 0.244220124 | 1 | 0.254971953 |
| 0 | 0 | 0 | 0 | 1 | 0.071428571 | 0.324324324 | 0.21875 | 1 | MED12 | 0.43631075 | 1 | 1 |
| 1 | 0.1 | 1 | 0.02 | 0 | 0 | 0.287301 | 1 | 0.253609774 | MEN1 | 0.408484834 | 1 | 0.371136255 |
| 1 | 0.1 | 0 | 0 | 0 | 0 | 0.135135135 | 1 | 0.135135135 | MGA | 0.244220124 | 1 | 0.254971953 |
| 2 | 0.2 | 1 | 0.02 | 1 | 0.071428571 | 0.057643405 | 0.392361111 | 0.085701175 | MITF | 0.244220124 | 1 | 0.254971953 |
| 2 | 0.2 | 0 | 0 | 0 | 0 | 0.016660496 | 1 | 0.016660496 | MLH1 | 0.096118247 | 1 | 0.090875433 |
| 1 | 0.1 | 0 | 0 | 0 | 0 | 0.135135135 | 1 | 0.135135135 | MLH3 | 0.244220124 | 1 | 0.254971953 |
| 1 | 0.1 | 0 | 0 | 0 | 0 | 0.135135135 | 1 | 0.135135135 | MPL | 0.244220124 | 1 | 0.254971953 |
| 0 | 0 | 0 | 0 | 1 | 0.071428571 | 0.324324324 | 0.21875 | 1 | MRE11A | 0.43631075 | 1 | 1 |
| 0 | 0 | 1 | 0.02 | 0 | 0 | 1 | 1 | 1 | MSH2 | 1 | 1 | 1 |
| 3 | 0.3 | 0 | 0 | 1 | 0.071428571 | 0.002512545 | 0.21875 | 0.006857137 | MSH6 | 0.047110225 | 1 | 0.073469324 |
| 2 | 0.2 | 0 | 0 | 0 | 0 | 0.016660496 | 1 | 0.016660496 | MTOR | 0.096118247 | 1 | 0.090875433 |
| 3 | 0.3 | 1 | 0.02 | 0 | 0 | 0.014449526 | 1 | 0.006857137 | MYBL1 | 0.096118247 | 1 | 0.073469324 |
| 0 | 0 | 1 | 0.02 | 0 | 0 | 1 | 1 | 1 | MYC | 1 | 1 | 1 |
| 1 | 0.1 | 0 | 0 | 1 | 0.071428571 | 0.102184376 | 0.21875 | 0.253609774 | MYCL | 0.244220124 | 1 | 0.371136255 |
| 0 | 0 | 1 | 0.02 | 0 | 0 | 1 | 1 | 1 | MYCN | 1 | 1 | 1 |
| 1 | 0.1 | 0 | 0 | 0 | 0 | 0.135135135 | 1 | 0.135135135 | MYD88 | 0.244220124 | 1 | 0.254971953 |
| 2 | 0.2 | 0 | 0 | 0 | 0 | 0.016660496 | 1 | 0.016660496 | NBN | 0.096118247 | 1 | 0.090875433 |
| 3 | 0.3 | 2 | 0.04 | 0 | 0 | 0.020737407 | 1 | 0.015867884 | NF1 | 0.109144249 | 1 | 0.090875433 |
| 1 | 0.1 | 0 | 0 | 0 | 0 | 0.135135135 | 1 | 0.135135135 | NF2 | 0.244220124 | 1 | 0.254971953 |
| 1 | 0.1 | 1 | 0.02 | 0 | 0 | 0.287301 | 1 | 0.253609774 | NFKBIA | 0.408484834 | 1 | 0.371136255 |
| 1 | 0.1 | 0 | 0 | 0 | 0 | 0.135135135 | 1 | 0.135135135 | NFKBIE | 0.244220124 | 1 | 0.254971953 |
| 2 | 0.2 | 0 | 0 | 0 | 0 | 0.016660496 | 1 | 0.016660496 | NFKBIZ | 0.096118247 | 1 | 0.090875433 |
| 2 | 0.2 | 4 | 0.08 | 0 | 0 | 0.189616178 | 0.568129737 | 0.183703933 | NOTCH1 | 0.328814181 | 1 | 0.342305465 |
| 1 | 0.1 | 2 | 0.04 | 0 | 0 | 0.433080341 | 1 | 0.357275083 | NOTCH2 | 0.560017682 | 1 | 0.491662959 |
| 1 | 0.1 | 1 | 0.02 | 1 | 0.071428571 | 0.244107121 | 0.392361111 | 0.357275083 | NOTCH3 | 0.408484834 | 1 | 0.491662959 |
| 1 | 0.1 | 0 | 0 | 0 | 0 | 0.135135135 | 1 | 0.135135135 | NPM1 | 0.244220124 | 1 | 0.254971953 |

| | | | | | | | | | | | | |
|---|---|---|---|---|---|---|---|---|---|---|---|---|
| 0 | 0 | 2 | 0.04 | 2 | 0.142857143 | 0.241977845 | 0.205667195 | 1 | NRAS | 0.408484834 | 1 | 1 |
| 0 | 0 | 0 | 0 | 1 | 0.071428571 | 0.324324324 | 0.21875 | 1 | NSD1 | 0.43631075 | 1 | 1 |
| 1 | 0.1 | 0 | 0 | 0 | 0 | 0.135135135 | 1 | 0.135135135 | NTHL1 | 0.244220124 | 1 | 0.254971953 |
| 1 | 0.1 | 1 | 0.02 | 0 | 0 | 0.287301 | 1 | 0.253609774 | NTRK1 | 0.408484834 | 1 | 0.371136255 |
| 2 | 0.2 | 2 | 0.04 | 0 | 0 | 0.1450958 | 1 | 0.085701175 | NTRK3 | 0.256051412 | 1 | 0.254971953 |
| 3 | 0.3 | 0 | 0 | 0 | 0 | 0.001851166 | 1 | 0.001851166 | PALB2 | 0.042719221 | 1 | 0.042719221 |
| 0 | 0 | 1 | 0.02 | 0 | 0 | 1 | 1 | 1 | PARK2 | 1 | 1 | 1 |
| 0 | 0 | 1 | 0.02 | 1 | 0.071428571 | 0.546464272 | 0.392361111 | 1 | PAX5 | 0.685938417 | 1 | 1 |
| 3 | 0.3 | 1 | 0.02 | 0 | 0 | 0.014449526 | 1 | 0.006857137 | PBRM1 | 0.096118247 | 1 | 0.073469324 |
| 1 | 0.1 | 0 | 0 | 0 | 0 | 0.135135135 | 1 | 0.135135135 | PDCD1LG2 | 0.244220124 | 1 | 0.254971953 |
| 0 | 0 | 0 | 0 | 2 | 0.142857143 | 0.050351722 | 0.045138889 | 1 | PDGFRA | 0.243637363 | 1 | 1 |
| 2 | 0.2 | 1 | 0.02 | 1 | 0.071428571 | 0.057643405 | 0.392361111 | 0.085701175 | PDGFRB | 0.244220124 | 1 | 0.254971953 |
| 1 | 0.1 | 0 | 0 | 0 | 0 | 0.135135135 | 1 | 0.135135135 | PHF6 | 0.244220124 | 1 | 0.254971953 |
| 0 | 0 | 2 | 0.04 | 0 | 0 | 1 | 1 | 1 | PHOX2B | 1 | 1 | 1 |
| 3 | 0.3 | 1 | 0.02 | 0 | 0 | 0.014449526 | 1 | 0.006857137 | PIK3C2B | 0.096118247 | 1 | 0.073469324 |
| 3 | 0.3 | 7 | 0.14 | 2 | 0.142857143 | 0.494836139 | 1 | 0.350519099 | PIK3CA | 0.634405306 | 1 | 0.491662959 |
| 2 | 0.2 | 3 | 0.06 | 0 | 0 | 0.175088169 | 1 | 0.132256702 | PIK3R1 | 0.307172227 | 1 | 0.254971953 |
| 0 | 0 | 1 | 0.02 | 0 | 0 | 1 | 1 | 1 | PML | 1 | 1 | 1 |
| 2 | 0.2 | 1 | 0.02 | 0 | 0 | 0.065932371 | 1 | 0.046279156 | PMS1 | 0.244220124 | 1 | 0.201213721 |
| 1 | 0.1 | 1 | 0.02 | 0 | 0 | 0.287301 | 1 | 0.253609774 | PMS2 | 0.408484834 | 1 | 0.371136255 |
| 1 | 0.1 | 0 | 0 | 0 | 0 | 0.135135135 | 1 | 0.135135135 | PNKP | 0.244220124 | 1 | 0.254971953 |
| 0 | 0 | 1 | 0.02 | 0 | 0 | 1 | 1 | 1 | POLD1 | 1 | 1 | 1 |
| 0 | 0 | 2 | 0.04 | 0 | 0 | 1 | 1 | 1 | POLE | 1 | 1 | 1 |
| 0 | 0 | 1 | 0.02 | 0 | 0 | 1 | 1 | 1 | POLQ | 1 | 1 | 1 |
| 1 | 0.1 | 0 | 0 | 0 | 0 | 0.135135135 | 1 | 0.135135135 | PPP2R1A | 0.244220124 | 1 | 0.254971953 |
| 2 | 0.2 | 0 | 0 | 0 | 0 | 0.016660496 | 1 | 0.016660496 | PRDM1 | 0.096118247 | 1 | 0.090875433 |
| 3 | 0.3 | 1 | 0.02 | 0 | 0 | 0.014449526 | 1 | 0.006857137 | PRF1 | 0.096118247 | 1 | 0.073469324 |
| 3 | 0.3 | 0 | 0 | 0 | 0 | 0.001851166 | 1 | 0.001851166 | PRKCI | 0.042719221 | 1 | 0.042719221 |
| 0 | 0 | 1 | 0.02 | 0 | 0 | 1 | 1 | 1 | PRKCZ | 1 | 1 | 1 |
| 5 | 0.5 | 3 | 0.06 | 2 | 0.142857143 | 0.00221791 | 0.299239086 | 0.002867312 | PRKDC | 0.047110225 | 1 | 0.053762103 |
| 1 | 0.1 | 1 | 0.02 | 0 | 0 | 0.287301 | 1 | 0.253609774 | PRPF40B | 0.408484834 | 1 | 0.371136255 |
| 3 | 0.3 | 0 | 0 | 0 | 0 | 0.001851166 | 1 | 0.001851166 | PRPF8 | 0.042719221 | 1 | 0.042719221 |
| 3 | 0.3 | 3 | 0.06 | 0 | 0 | 0.044012072 | 1 | 0.02936224 | PTCH1 | 0.216452815 | 1 | 0.149299524 |
| 1 | 0.1 | 3 | 0.06 | 1 | 0.071428571 | 0.799848083 | 1 | 0.526685474 | PTEN | 0.979405816 | 1 | 0.696060098 |
| 3 | 0.3 | 0 | 0 | 0 | 0 | 0.001851166 | 1 | 0.001851166 | PTK2 | 0.042719221 | 1 | 0.042719221 |
| 1 | 0.1 | 0 | 0 | 1 | 0.071428571 | 0.102184376 | 0.21875 | 0.253609774 | PTK2B | 0.244220124 | 1 | 0.371136255 |
| 1 | 0.1 | 0 | 0 | 0 | 0 | 0.135135135 | 1 | 0.135135135 | PTPN11 | 0.244220124 | 1 | 0.254971953 |
| 0 | 0 | 1 | 0.02 | 0 | 0 | 1 | 1 | 1 | PTPN14 | 1 | 1 | 1 |
| 2 | 0.2 | 1 | 0.02 | 0 | 0 | 0.065932371 | 1 | 0.046279156 | PTPRD | 0.244220124 | 1 | 0.201213721 |
| 1 | 0.1 | 0 | 0 | 0 | 0 | 0.135135135 | 1 | 0.135135135 | RAD51 | 0.244220124 | 1 | 0.254971953 |
| 1 | 0.1 | 0 | 0 | 0 | 0 | 0.135135135 | 1 | 0.135135135 | RAF1 | 0.244220124 | 1 | 0.254971953 |
| 0 | 0 | 1 | 0.02 | 0 | 0 | 1 | 1 | 1 | RARA | 1 | 1 | 1 |
| 1 | 0.1 | 0 | 0 | 0 | 0 | 0.135135135 | 1 | 0.135135135 | RASA1 | 0.244220124 | 1 | 0.254971953 |
| 1 | 0.1 | 1 | 0.02 | 0 | 0 | 0.287301 | 1 | 0.253609774 | RB1 | 0.408484834 | 1 | 0.371136255 |
| 0 | 0 | 1 | 0.02 | 0 | 0 | 1 | 1 | 1 | RBBP8 | 1 | 1 | 1 |
| 1 | 0.1 | 0 | 0 | 0 | 0 | 0.135135135 | 1 | 0.135135135 | RBM10 | 0.244220124 | 1 | 0.254971953 |
| 1 | 0.1 | 1 | 0.02 | 0 | 0 | 0.287301 | 1 | 0.253609774 | RECQL4 | 0.408484834 | 1 | 0.371136255 |
| 1 | 0.1 | 0 | 0 | 0 | 0 | 0.135135135 | 1 | 0.135135135 | REL | 0.244220124 | 1 | 0.254971953 |
| 1 | 0.1 | 3 | 0.06 | 2 | 0.142857143 | 0.438331452 | 0.299239086 | 1 | RET | 0.56437526 | 1 | 1 |
| 1 | 0.1 | 0 | 0 | 0 | 0 | 0.135135135 | 1 | 0.135135135 | RFWD2 | 0.244220124 | 1 | 0.254971953 |
| 1 | 0.1 | 1 | 0.02 | 0 | 0 | 0.287301 | 1 | 0.253609774 | RHBDF2 | 0.408484834 | 1 | 0.371136255 |
| 1 | 0.1 | 0 | 0 | 0 | 0 | 0.135135135 | 1 | 0.135135135 | RHOA | 0.244220124 | 1 | 0.254971953 |
| 0 | 0 | 1 | 0.02 | 0 | 0 | 1 | 1 | 1 | RHOH | 1 | 1 | 1 |
| 1 | 0.1 | 0 | 0 | 0 | 0 | 0.135135135 | 1 | 0.135135135 | RHPN2 | 0.244220124 | 1 | 0.254971953 |
| 0 | 0 | 0 | 0 | 1 | 0.071428571 | 0.324324324 | 0.21875 | 1 | RIF1 | 0.43631075 | 1 | 1 |
| 1 | 0.1 | 1 | 0.02 | 0 | 0 | 0.287301 | 1 | 0.253609774 | RNF43 | 0.408484834 | 1 | 0.371136255 |
| 0 | 0 | 1 | 0.02 | 0 | 0 | 1 | 1 | 1 | ROS1 | 1 | 1 | 1 |
| 0 | 0 | 1 | 0.02 | 0 | 0 | 1 | 1 | 1 | RSPO3 | 1 | 1 | 1 |
| 1 | 0.1 | 0 | 0 | 0 | 0 | 0.135135135 | 1 | 0.135135135 | RUNX1 | 0.244220124 | 1 | 0.254971953 |
| 1 | 0.1 | 2 | 0.04 | 0 | 0 | 0.433080341 | 1 | 0.357275083 | SDHA | 0.560017682 | 1 | 0.491662959 |
| 1 | 0.1 | 0 | 0 | 0 | 0 | 0.135135135 | 1 | 0.135135135 | SDHAF2 | 0.244220124 | 1 | 0.254971953 |
| 0 | 0 | 1 | 0.02 | 0 | 0 | 1 | 1 | 1 | SDHD | 1 | 1 | 1 |
| 3 | 0.3 | 1 | 0.02 | 1 | 0.071428571 | 0.01161194 | 0.392361111 | 0.015867884 | SETBP1 | 0.096118247 | 1 | 0.090875433 |
| 1 | 0.1 | 2 | 0.04 | 1 | 0.071428571 | 0.391027145 | 0.529569892 | 0.447799719 | SETD2 | 0.519062582 | 1 | 0.599731767 |
| 1 | 0.1 | 1 | 0.02 | 0 | 0 | 0.287301 | 1 | 0.253609774 | SF1 | 0.408484834 | 1 | 0.371136255 |
| 2 | 0.2 | 0 | 0 | 0 | 0 | 0.016660496 | 1 | 0.016660496 | SF3B1 | 0.096118247 | 1 | 0.090875433 |
| 2 | 0.2 | 1 | 0.02 | 0 | 0 | 0.065932371 | 1 | 0.046279156 | SH2B3 | 0.244220124 | 1 | 0.201213721 |
| 1 | 0.1 | 0 | 0 | 0 | 0 | 0.135135135 | 1 | 0.135135135 | SLC25A13 | 0.244220124 | 1 | 0.254971953 |
| 1 | 0.1 | 0 | 0 | 0 | 0 | 0.135135135 | 1 | 0.135135135 | SLC34A2 | 0.244220124 | 1 | 0.254971953 |
| 2 | 0.2 | 0 | 0 | 0 | 0 | 0.016660496 | 1 | 0.016660496 | SLITRK6 | 0.096118247 | 1 | 0.090875433 |
| 1 | 0.1 | 1 | 0.02 | 0 | 0 | 0.287301 | 1 | 0.253609774 | SLX4 | 0.408484834 | 1 | 0.371136255 |
| 0 | 0 | 1 | 0.02 | 0 | 0 | 1 | 1 | 1 | SMAD2 | 1 | 1 | 1 |
| 0 | 0 | 11 | 0.22 | 2 | 0.142857143 | 0.26168066 | 0.715372654 | 0.192779347 | SMAD4 | 0.408484834 | 1 | 0.354808614 |
| 6 | 0.6 | 0 | 0 | 0 | 0 | 1.1336E-06 | 1 | 1.1336E-06 | SMARCA4 | 0.00034008 | 1 | 0.000332249 |
| 0 | 0 | 1 | 0.02 | 0 | 0 | 1 | 1 | 1 | SMARCE1 | 1 | 1 | 1 |
| 2 | 0.2 | 0 | 0 | 0 | 0 | 0.016660496 | 1 | 0.016660496 | SMC1A | 0.096118247 | 1 | 0.090875433 |
| 2 | 0.2 | 0 | 0 | 2 | 0.142857143 | 0.009234973 | 0.045138889 | 0.085701175 | SMC3 | 0.096118247 | 1 | 0.254971953 |
| 2 | 0.2 | 1 | 0.02 | 0 | 0 | 0.065932371 | 1 | 0.046279156 | SMO | 0.244220124 | 1 | 0.201213721 |
| 1 | 0.1 | 0 | 0 | 0 | 0 | 0.135135135 | 1 | 0.135135135 | SOCS1 | 0.244220124 | 1 | 0.254971953 |
| 0 | 0 | 2 | 0.04 | 0 | 0 | 1 | 1 | 1 | SOX2 | 1 | 1 | 1 |
| 5 | 0.5 | 7 | 0.14 | 2 | 0.142857143 | 0.039666451 | 1 | 0.017429111 | SOX9 | 0.198332255 | 1 | 0.093370236 |
| 1 | 0.1 | 0 | 0 | 0 | 0 | 0.135135135 | 1 | 0.135135135 | SQSTM1 | 0.244220124 | 1 | 0.254971953 |
| 2 | 0.2 | 1 | 0.02 | 0 | 0 | 0.065932371 | 1 | 0.046279156 | SRSF2 | 0.244220124 | 1 | 0.201213721 |
| 1 | 0.1 | 0 | 0 | 0 | 0 | 0.135135135 | 1 | 0.135135135 | STAG2 | 0.244220124 | 1 | 0.254971953 |
| 2 | 0.2 | 0 | 0 | 0 | 0 | 0.016660496 | 1 | 0.016660496 | STAT3 | 0.096118247 | 1 | 0.090875433 |
| 1 | 0.1 | 0 | 0 | 0 | 0 | 0.135135135 | 1 | 0.135135135 | STAT6 | 0.244220124 | 1 | 0.254971953 |
| 1 | 0.1 | 1 | 0.02 | 0 | 0 | 0.287301 | 1 | 0.253609774 | STK11 | 0.408484834 | 1 | 0.371136255 |
| 1 | 0.1 | 1 | 0.02 | 1 | 0.071428571 | 0.244107121 | 0.392361111 | 0.357275083 | SUFU | 0.408484834 | 1 | 0.491662959 |
| 0 | 0 | 1 | 0.02 | 1 | 0.071428571 | 0.546464272 | 0.392361111 | 1 | SUZ12 | 0.685938417 | 1 | 1 |
| 1 | 0.1 | 1 | 0.02 | 0 | 0 | 0.287301 | 1 | 0.253609774 | SYK | 0.408484834 | 1 | 0.371136255 |
| 1 | 0.1 | 0 | 0 | 0 | 0 | 0.135135135 | 1 | 0.135135135 | TCF3 | 0.244220124 | 1 | 0.254971953 |
| 1 | 0.1 | 0 | 0 | 0 | 0 | 0.135135135 | 1 | 0.135135135 | TCF7L1 | 0.244220124 | 1 | 0.254971953 |
| 0 | 0 | 6 | 0.12 | 1 | 0.071428571 | 0.835169009 | 1 | 0.583378305 | TCF7L2 | 1 | 1 | 0.767603033 |
| 1 | 0.1 | 2 | 0.04 | 1 | 0.071428571 | 0.391027145 | 0.529569892 | 0.447799719 | TERT | 0.519062582 | 1 | 0.599731767 |

| | | | | | | | | | | | | |
|---|---|---|---|---|---|---|---|---|---|---|---|---|
| 1 | 0.1 | 0 | 0 | 0 | 0 | 0.135135135 | 1 | 0.135135135 | TET1 | 0.244220124 | 1 | 0.254971953 |
| 2 | 0.2 | 4 | 0.08 | 0 | 0 | 0.189616178 | 0.568129737 | 0.183703933 | TET2 | 0.328814181 | 1 | 0.342305465 |
| 1 | 0.1 | 0 | 0 | 0 | 0 | 0.135135135 | 1 | 0.135135135 | TMPRSS2 | 0.244220124 | 1 | 0.254971953 |
| 1 | 0.1 | 1 | 0.02 | 0 | 0 | 0.287301 | 1 | 0.253609774 | TNFAIP3 | 0.408484834 | 1 | 0.371136255 |
| 2 | 0.2 | 38 | 0.76 | 8 | 0.571428571 | 0.00243318 | 0.190220989 | 0.002669151 | TP53 | 0.047110225 | 1 | 0.053762103 |
| 1 | 0.1 | 0 | 0 | 0 | 0 | 0.135135135 | 1 | 0.135135135 | TP53BP1 | 0.244220124 | 1 | 0.254971953 |
| 1 | 0.1 | 0 | 0 | 0 | 0 | 0.135135135 | 1 | 0.135135135 | TRAF3 | 0.244220124 | 1 | 0.254971953 |
| 0 | 0 | 2 | 0.04 | 0 | 0 | 1 | 1 | 1 | TRBC2 | 1 | 1 | 1 |
| 0 | 0 | 1 | 0.02 | 0 | 0 | 1 | 1 | 1 | TRIM37 | 1 | 1 | 1 |
| 2 | 0.2 | 0 | 0 | 0 | 0 | 0.016660496 | 1 | 0.016660496 | TSC1 | 0.096118247 | 1 | 0.090875433 |
| 3 | 0.3 | 1 | 0.02 | 0 | 0 | 0.014449526 | 1 | 0.006857137 | TSC2 | 0.096118247 | 1 | 0.073469324 |
| 1 | 0.1 | 1 | 0.02 | 0 | 0 | 0.287301 | 1 | 0.253609774 | U2AF1 | 0.408484834 | 1 | 0.371136255 |
| 1 | 0.1 | 0 | 0 | 0 | 0 | 0.135135135 | 1 | 0.135135135 | VHL | 0.244220124 | 1 | 0.254971953 |
| 0 | 0 | 2 | 0.04 | 0 | 0 | 1 | 1 | 1 | WRN | 1 | 1 | 1 |
| 1 | 0.1 | 1 | 0.02 | 0 | 0 | 0.287301 | 1 | 0.253609774 | WT1 | 0.408484834 | 1 | 0.371136255 |
| 1 | 0.1 | 0 | 0 | 0 | 0 | 0.135135135 | 1 | 0.135135135 | XPC | 0.244220124 | 1 | 0.254971953 |
| 1 | 0.1 | 0 | 0 | 0 | 0 | 0.135135135 | 1 | 0.135135135 | XRCC1 | 0.244220124 | 1 | 0.254971953 |
| 3 | 0.3 | 1 | 0.02 | 0 | 0 | 0.014449526 | 1 | 0.006857137 | ZNF217 | 0.096118247 | 1 | 0.073469324 |
| 0 | 0 | 1 | 0.02 | 0 | 0 | 1 | 1 | 1 | ZNF708 | 1 | 1 | 1 |
| 1 | 0.1 | 1 | 0.02 | 0 | 0 | 0.287301 | 1 | 0.253609774 | ZNRF3 | 0.408484834 | 1 | 0.371136255 |
| 1 | 0.1 | 0 | 0 | 1 | 0.071428571 | 0.102184376 | 0.21875 | 0.253609774 | ZRSR2 | 0.244220124 | 1 | 0.371136255 |

**Supplementary Table 3**: *_N: Number of dMMR/tdpMMR/tipMMR tumors with SCNA; *_fraction: Fraction of dMMR/tdpMMR/tipMMR tumors with SCNA; pvalue_3way: p-value (Fisher's exact test) for enrichment in SCNA frequency across all three T cell/MMR groups; pvalue_tip_vs_tdp: p-value (Fisher's exact test) for enrichment in SCNA frequency between tdpMMR versus tipMMR tumors; pvalue_pmmr_vs_dmmr: p-value (Fisher's exact test) for enrichment in SCNA frequency between all pMMR versus dMMR tumors; qvalue_*:

| dMMR_N | dMMR fraction | tdpMMR_N | tdpMMR fraction | tipMMR_N | tipMMR fraction | pvalue_3way | pvalue tip_vs_tdp | pmmr_vs_d mmr | gene | qvalue 3way | qvalue tip_vs_tdp | qvalue pmmr_vs_dmmr |
|---|---|---|---|---|---|---|---|---|---|---|---|---|
| 0 | 0 | 2 | 0.04 | 1 | 0.07143 | 0.697642848 | 0.5295699 | 1 | ABCB11 | 1 | 1 | 1 |
| 0 | 0 | 8 | 0.16 | 1 | 0.07143 | 0.421541949 | 0.6699605 | 0.346600866 | ABL1 | 1 | 1 | 0.94492303 |
| 0 | 0 | 5 | 0.1 | 0 | 0 | 0.48654037 | 0.577127 | 1 | ABRAXAS1 | 1 | 1 | 1 |
| 0 | 0 | 3 | 0.06 | 0 | 0 | 1 | 1 | 1 | ACVR1 | 1 | 1 | 1 |
| 0 | 0 | 7 | 0.14 | 1 | 0.07143 | 0.62851523 | 0.6732942 | 0.587819901 | AKT1 | 1 | 1 | 1 |
| 0 | 0 | 3 | 0.06 | 1 | 0.07143 | 1 | 1 | 1 | AKT2 | 1 | 1 | 1 |
| 0 | 0 | 7 | 0.14 | 1 | 0.07143 | 0.62851523 | 0.6732942 | 0.587819901 | AKT3 | 1 | 1 | 1 |
| 0 | 0 | 2 | 0.04 | 0 | 0 | 1 | 1 | 1 | ALK | 1 | 1 | 1 |
| 0 | 0 | 12 | 0.24 | 4 | 0.28571 | 0.181669295 | 0.7355945 | 0.105743209 | ALOX12B | 0.8607244 | 1 | 0.94492303 |
| 1 | 0.1 | 5 | 0.1 | 1 | 0.07143 | 1 | 1 | 1 | APC | 1 | 1 | 1 |
| 0 | 0 | 2 | 0.04 | 0 | 0 | 1 | 1 | 1 | AR | 1 | 1 | 1 |
| 0 | 0 | 2 | 0.04 | 0 | 0 | 1 | 1 | 1 | ARAF | 1 | 1 | 1 |
| 0 | 0 | 4 | 0.08 | 1 | 0.07143 | 1 | 1 | 1 | ARHGAP35 | 1 | 1 | 1 |
| 0 | 0 | 2 | 0.04 | 1 | 0.07143 | 0.697642848 | 0.5295699 | 1 | ARHGEF12 | 1 | 1 | 1 |
| 0 | 0 | 7 | 0.14 | 1 | 0.07143 | 0.62851523 | 0.6732942 | 0.587819901 | ARID1A | 1 | 1 | 1 |
| 0 | 0 | 7 | 0.14 | 3 | 0.21429 | 0.442160876 | 0.6774614 | 0.33939492 | ARID1B | 1 | 1 | 0.94492303 |
| 1 | 0.1 | 10 | 0.2 | 5 | 0.35714 | 0.358128075 | 0.2861485 | 0.679825377 | ARID2 | 0.9763492 | 1 | 1 |
| 0 | 0 | 27 | 0.54 | 5 | 0.35714 | 0.003275383 | 0.3649203 | 0.00377781 | ASXL1 | 0.1730076 | 1 | 0.248677373 |
| 0 | 0 | 7 | 0.14 | 1 | 0.07143 | 0.62851523 | 0.6732942 | 0.587819901 | ATM | 1 | 1 | 1 |
| 0 | 0 | 4 | 0.08 | 1 | 0.07143 | 1 | 1 | 1 | ATR | 1 | 1 | 1 |
| 0 | 0 | 1 | 0.02 | 0 | 0 | 1 | 1 | 1 | ATRX | 1 | 1 | 1 |
| 0 | 0 | 25 | 0.5 | 6 | 0.42857 | 0.008508571 | 0.7652595 | 0.003937281 | AURKA | 0.1730076 | 1 | 0.248677373 |
| 0 | 0 | 19 | 0.38 | 3 | 0.21429 | 0.033911962 | 0.3456933 | 0.02789446 | AURKB | 0.3761145 | 1 | 0.618749838 |
| 0 | 0 | 7 | 0.14 | 3 | 0.21429 | 0.442160876 | 0.6774614 | 0.33939492 | AXIN2 | 1 | 1 | 0.94492303 |
| 0 | 0 | 1 | 0.02 | 0 | 0 | 1 | 1 | 1 | AXL | 1 | 1 | 1 |
| 0 | 0 | 7 | 0.14 | 2 | 0.14286 | 0.652062514 | 1 | 0.346600866 | B2M | 1 | 1 | 0.94492303 |
| 0 | 0 | 1 | 0.02 | 0 | 0 | 1 | 1 | 1 | BABAM2 | 1 | 1 | 1 |
| 0 | 0 | 5 | 0.1 | 2 | 0.14286 | 0.595089088 | 0.6418802 | 0.583378305 | BAP1 | 1 | 1 | 1 |
| 0 | 0 | 2 | 0.04 | 0 | 0 | 1 | 1 | 1 | BARD1 | 1 | 1 | 1 |
| 0 | 0 | 6 | 0.12 | 0 | 0 | 0.342051159 | 0.3248367 | 0.588422151 | BCL11B | 0.9763492 | 1 | 1 |
| 2 | 0.2 | 25 | 0.5 | 3 | 0.21429 | 0.062917015 | 0.0722331 | 0.186529271 | BCL2 | 0.4797422 | 1 | 0.94492303 |
| 0 | 0 | 26 | 0.52 | 4 | 0.28571 | 0.002882635 | 0.1420079 | 0.004371918 | BCL2L1 | 0.1730076 | 1 | 0.248677373 |
| 0 | 0 | 2 | 0.04 | 2 | 0.14286 | 0.241977845 | 0.2056672 | 1 | BCL2L12 | 0.9091937 | 1 | 1 |
| 0 | 0 | 2 | 0.04 | 1 | 0.07143 | 0.697642848 | 0.5295699 | 1 | BCL6 | 1 | 1 | 1 |
| 0 | 0 | 2 | 0.04 | 0 | 0 | 1 | 1 | 1 | BCOR | 1 | 1 | 1 |
| 0 | 0 | 3 | 0.06 | 5 | 0.35714 | 0.007856633 | 0.0097358 | 0.587819901 | BCORL1 | 0.1730076 | 1 | 1 |
| 0 | 0 | 7 | 0.14 | 2 | 0.14286 | 0.652062514 | 1 | 0.346600866 | BLM | 1 | 1 | 0.94492303 |
| 0 | 0 | 10 | 0.2 | 0 | 0 | 0.073759985 | 0.100854 | 0.33939492 | BMPR1A | 0.50697 | 1 | 0.94492303 |
| 1 | 0.1 | 16 | 0.32 | 4 | 0.28571 | 0.4626501 | 1 | 0.264290836 | BRAF | 1 | 1 | 0.94492303 |
| 0 | 0 | 10 | 0.2 | 2 | 0.14286 | 0.399567541 | 1 | 0.199914832 | BRCA1 | 1 | 1 | 0.94492303 |
| 0 | 0 | 26 | 0.52 | 4 | 0.28571 | 0.002882635 | 0.1420079 | 0.004371918 | BRCA2 | 0.1730076 | 1 | 0.248677373 |
| 0 | 0 | 3 | 0.06 | 2 | 0.14286 | 0.355011967 | 0.2992391 | 1 | BRCC3 | 0.9763492 | 1 | 1 |
| 0 | 0 | 2 | 0.04 | 0 | 0 | 1 | 1 | 1 | BRD3 | 1 | 1 | 1 |
| 0 | 0 | 2 | 0.04 | 0 | 0 | 1 | 1 | 1 | BRD4 | 1 | 1 | 1 |
| 0 | 0 | 9 | 0.18 | 2 | 0.14286 | 0.471161887 | 1 | 0.341106332 | BRIP1 | 1 | 1 | 0.94492303 |
| 0 | 0 | 9 | 0.18 | 5 | 0.35714 | 0.101447848 | 0.2701668 | 0.192132251 | BUB1B | 0.6519717 | 1 | 0.94492303 |
| 0 | 0 | 1 | 0.02 | 0 | 0 | 1 | 1 | 1 | CADM2 | 1 | 1 | 1 |
| 1 | 0.1 | 16 | 0.32 | 4 | 0.28571 | 0.4626501 | 1 | 0.264290836 | CARD11 | 1 | 1 | 0.94492303 |
| 0 | 0 | 2 | 0.04 | 0 | 0 | 1 | 1 | 1 | CASP8 | 1 | 1 | 1 |
| 0 | 0 | 4 | 0.08 | 3 | 0.21429 | 0.21053707 | 0.1707213 | 0.583378305 | CBFA2T3 | 0.8958294 | 1 | 1 |
| 0 | 0 | 1 | 0.02 | 1 | 0.07143 | 0.546464272 | 0.3923611 | 1 | CBFB | 1 | 1 | 1 |
| 0 | 0 | 10 | 0.2 | 3 | 0.21429 | 0.297408165 | 1 | 0.192779347 | CBL | 0.9763492 | 1 | 0.94492303 |
| 0 | 0 | 3 | 0.06 | 1 | 0.07143 | 1 | 1 | 1 | CBLB | 1 | 1 | 1 |
| 0 | 0 | 5 | 0.1 | 2 | 0.14286 | 0.595089088 | 0.6418802 | 0.583378305 | CCND1 | 1 | 1 | 1 |
| 1 | 0.1 | 9 | 0.18 | 2 | 0.14286 | 1 | 1 | 1 | CCND2 | 1 | 1 | 1 |
| 0 | 0 | 8 | 0.16 | 2 | 0.14286 | 0.510166839 | 1 | 0.33939492 | CCND3 | 1 | 1 | 0.94492303 |
| 0 | 0 | 11 | 0.22 | 5 | 0.35714 | 0.106620585 | 0.312148 | 0.105743209 | CCNE1 | 0.6519717 | 1 | 0.94492303 |
| 0 | 0 | 7 | 0.14 | 1 | 0.07143 | 0.62851523 | 0.6732942 | 0.587819901 | CD274 | 1 | 1 | 1 |
| 0 | 0 | 4 | 0.08 | 0 | 0 | 0.76152112 | 0.5681297 | 1 | CD58 | 1 | 1 | 1 |
| 0 | 0 | 9 | 0.18 | 2 | 0.14286 | 0.471161887 | 1 | 0.341106332 | CD79B | 1 | 1 | 0.94492303 |
| 0 | 0 | 14 | 0.28 | 6 | 0.42857 | 0.051998521 | 0.3364219 | 0.05331369 | CDC73 | 0.4375048 | 1 | 0.847858997 |
| 0 | 0 | 3 | 0.06 | 1 | 0.07143 | 1 | 1 | 1 | CDH1 | 1 | 1 | 1 |
| 0 | 0 | 10 | 0.2 | 2 | 0.14286 | 0.399567541 | 1 | 0.199914832 | CDH4 | 1 | 1 | 0.94492303 |
| 0 | 0 | 5 | 0.1 | 0 | 0 | 0.48654037 | 0.577127 | 1 | CDK1 | 1 | 1 | 1 |
| 0 | 0 | 7 | 0.14 | 2 | 0.14286 | 0.652062514 | 1 | 0.346600866 | CDK12 | 1 | 1 | 0.94492303 |
| 1 | 0.1 | 2 | 0.04 | 1 | 0.07143 | 0.391027145 | 0.5295699 | 0.447799719 | CDK2 | 1 | 1 | 1 |
| 1 | 0.1 | 8 | 0.16 | 0 | 0 | 0.301201587 | 0.1831159 | 1 | CDK4 | 0.9763492 | 1 | 1 |
| 1 | 0.1 | 10 | 0.2 | 4 | 0.28571 | 0.505412483 | 0.4850325 | 0.675901668 | CDK5 | 1 | 1 | 1 |
| 1 | 0.1 | 15 | 0.3 | 5 | 0.35714 | 0.398445876 | 0.748793 | 0.264290836 | CDK6 | 1 | 1 | 0.94492303 |
| 0 | 0 | 14 | 0.28 | 1 | 0.07143 | 0.059436703 | 0.1572291 | 0.197470798 | CDK8 | 0.4678244 | 1 | 0.94492303 |
| 0 | 0 | 6 | 0.12 | 1 | 0.07143 | 0.835169009 | 1 | 0.583378305 | CDK9 | 1 | 1 | 1 |
| 0 | 0 | 8 | 0.16 | 2 | 0.14286 | 0.510166839 | 1 | 0.33939492 | CDKN1A | 1 | 1 | 0.94492303 |
| 1 | 0.1 | 10 | 0.2 | 1 | 0.07143 | 0.5603718 | 0.43058 | 1 | CDKN1B | 1 | 1 | 1 |
| 1 | 0.1 | 17 | 0.34 | 9 | 0.64286 | 0.02549746 | 0.0640089 | 0.082122838 | CDKN1C | 0.3148752 | 1 | 0.94492303 |

| | | | | | | | | | | | | |
|---|---|---|---|---|---|---|---|---|---|---|---|---|
| 0 | 0 | 5 | 0.1 | 1 | 0.07143 | 0.825954855 | 1 | 0.588422151 | CDKN2A | 1 | 1 | 1 |
| 0 | 0 | 4 | 0.08 | 0 | 0 | 0.76152112 | 0.5681297 | 1 | CDKN2B | 1 | 1 | 1 |
| 0 | 0 | 5 | 0.1 | 0 | 0 | 0.48654037 | 0.577127 | 1 | CDKN2C | 1 | 1 | 1 |
| 0 | 0 | 2 | 0.04 | 0 | 0 | 1 | 1 | 1 | CEBPA | 1 | 1 | 1 |
| 0 | 0 | 6 | 0.12 | 6 | 0.42857 | 0.008374935 | 0.016967 | 0.199914832 | CHEK1 | 0.1730076 | 1 | 0.94492303 |
| 0 | 0 | 9 | 0.18 | 0 | 0 | 0.122866881 | 0.1870181 | 0.346600866 | CHEK2 | 0.6919557 | 1 | 0.94492303 |
| 0 | 0 | 11 | 0.22 | 5 | 0.35714 | 0.106620585 | 0.312148 | 0.105743209 | CIC | 0.6519717 | 1 | 0.94492303 |
| 0 | 0 | 13 | 0.26 | 7 | 0.5 | 0.023889615 | 0.1091393 | 0.05331369 | CIITA | 0.3148752 | 1 | 0.847858997 |
| 0 | 0 | 3 | 0.06 | 2 | 0.14286 | 0.355011967 | 0.2992391 | 1 | COL7A1 | 0.9763492 | 1 | 1 |
| 0 | 0 | 4 | 0.08 | 1 | 0.07143 | 1 | 1 | 1 | COP1 | 1 | 1 | 1 |
| 0 | 0 | 7 | 0.14 | 4 | 0.28571 | 0.164188161 | 0.2371794 | 0.341106332 | CREBBP | 0.8012382 | 1 | 0.94492303 |
| 0 | 0 | 15 | 0.3 | 0 | 0 | 0.008408547 | 0.0277545 | 0.197470798 | CRKL | 0.1730076 | 1 | 0.94492303 |
| 0 | 0 | 2 | 0.04 | 0 | 0 | 1 | 1 | 1 | CRLF2 | 1 | 1 | 1 |
| 0 | 0 | 2 | 0.04 | 0 | 0 | 1 | 1 | 1 | CRTC1 | 1 | 1 | 1 |
| 0 | 0 | 3 | 0.06 | 2 | 0.14286 | 0.355011967 | 0.2992391 | 1 | CRTC2 | 0.9763492 | 1 | 1 |
| 0 | 0 | 1 | 0.02 | 0 | 0 | 1 | 1 | 1 | CSF1R | 1 | 1 | 1 |
| 0 | 0 | 6 | 0.12 | 4 | 0.28571 | 0.12702587 | 0.2058665 | 0.33939492 | CSF3R | 0.6919557 | 1 | 0.94492303 |
| 0 | 0 | 1 | 0.02 | 1 | 0.07143 | 0.546464272 | 0.3923611 | 1 | CTCF | 1 | 1 | 1 |
| 0 | 0 | 2 | 0.04 | 0 | 0 | 1 | 1 | 1 | CTLA4 | 1 | 1 | 1 |
| 0 | 0 | 7 | 0.14 | 1 | 0.07143 | 0.62851523 | 0.6732942 | 0.587819901 | CTNNA1 | 1 | 1 | 1 |
| 0 | 0 | 5 | 0.1 | 3 | 0.21429 | 0.26269735 | 0.357359 | 0.587819901 | CTNNB1 | 0.9091937 | 1 | 1 |
| 1 | 0.1 | 17 | 0.34 | 5 | 0.35714 | 0.357763426 | 1 | 0.158337716 | CUX1 | 0.9763492 | 1 | 0.94492303 |
| 0 | 0 | 14 | 0.28 | 5 | 0.35714 | 0.1068806 | 0.7416156 | 0.055597311 | CXCR4 | 0.6519717 | 1 | 0.847858997 |
| 0 | 0 | 6 | 0.12 | 3 | 0.21429 | 0.353535986 | 0.3970432 | 0.346600866 | CYLD | 0.9763492 | 1 | 0.94492303 |
| 0 | 0 | 9 | 0.18 | 4 | 0.28571 | 0.229201109 | 0.4566205 | 0.192779347 | DAXX | 0.9091937 | 1 | 0.94492303 |
| 0 | 0 | 2 | 0.04 | 2 | 0.14286 | 0.241977845 | 0.2056672 | 1 | DCLRE1C | 0.9091937 | 1 | 1 |
| 0 | 0 | 7 | 0.14 | 2 | 0.14286 | 0.652062514 | 1 | 0.346600866 | DDB1 | 1 | 1 | 0.94492303 |
| 0 | 0 | 8 | 0.16 | 3 | 0.21429 | 0.416615438 | 0.6933892 | 0.341106332 | DDB2 | 1 | 1 | 0.94492303 |
| 0 | 0 | 8 | 0.16 | 4 | 0.28571 | 0.210057837 | 0.4376874 | 0.199914832 | DDR2 | 0.8958294 | 1 | 0.94492303 |
| 0 | 0 | 4 | 0.08 | 0 | 0 | 0.76152112 | 0.5681297 | 1 | DEPDC5 | 1 | 1 | 1 |
| 0 | 0 | 6 | 0.12 | 1 | 0.07143 | 0.835169009 | 1 | 0.583378305 | DICER1 | 1 | 1 | 1 |
| 0 | 0 | 25 | 0.5 | 3 | 0.21429 | 0.002747671 | 0.0722331 | 0.01061248 | DIS3 | 0.1730076 | 1 | 0.383035181 |
| 0 | 0 | 5 | 0.1 | 0 | 0 | 0.48654037 | 0.577127 | 1 | DIS3L2 | 1 | 1 | 1 |
| 0 | 0 | 2 | 0.04 | 0 | 0 | 1 | 1 | 1 | DKC1 | 1 | 1 | 1 |
| 0 | 0 | 5 | 0.1 | 0 | 0 | 0.48654037 | 0.577127 | 1 | DMC1 | 1 | 1 | 1 |
| 0 | 0 | 0 | 0 | 1 | 0.07143 | 0.324324324 | 0.21875 | 1 | DMD | 0.9763492 | 1 | 1 |
| 0 | 0 | 4 | 0.08 | 0 | 0 | 0.76152112 | 0.5681297 | 1 | DNMT3A | 1 | 1 | 1 |
| 0 | 0 | 2 | 0.04 | 1 | 0.07143 | 0.697642848 | 0.5295699 | 1 | DOCK8 | 1 | 1 | 1 |
| 0 | 0 | 4 | 0.08 | 0 | 0 | 0.76152112 | 0.5681297 | 1 | EED | 1 | 1 | 1 |
| 1 | 0.1 | 16 | 0.32 | 4 | 0.28571 | 0.4626501 | 1 | 0.264290836 | EGFR | 1 | 1 | 0.94492303 |
| 0 | 0 | 4 | 0.08 | 1 | 0.07143 | 1 | 1 | 1 | EGLN1 | 1 | 1 | 1 |
| 0 | 0 | 5 | 0.1 | 2 | 0.14286 | 0.595089088 | 0.6418802 | 0.583378305 | ELOC | 1 | 1 | 1 |
| 0 | 0 | 5 | 0.1 | 1 | 0.07143 | 0.825954855 | 1 | 0.588422151 | EME1 | 1 | 1 | 1 |
| 0 | 0 | 2 | 0.04 | 0 | 0 | 1 | 1 | 1 | ENG | 1 | 1 | 1 |
| 0 | 0 | 14 | 0.28 | 0 | 0 | 0.014317702 | 0.0278514 | 0.192132251 | EP300 | 0.2743679 | 1 | 0.94492303 |
| 0 | 0 | 3 | 0.06 | 0 | 0 | 1 | 1 | 1 | EPCAM | 1 | 1 | 1 |
| 0 | 0 | 0 | 0 | 1 | 0.07143 | 0.324324324 | 0.21875 | 1 | EPHA3 | 0.9763492 | 1 | 1 |
| 0 | 0 | 5 | 0.1 | 1 | 0.07143 | 0.825954855 | 1 | 0.588422151 | EPHA5 | 1 | 1 | 1 |
| 0 | 0 | 7 | 0.14 | 3 | 0.21429 | 0.442160876 | 0.6774614 | 0.33939492 | EPHA7 | 1 | 1 | 0.94492303 |
| 0 | 0 | 10 | 0.2 | 2 | 0.14286 | 0.399567541 | 1 | 0.199914832 | ERBB2 | 1 | 1 | 0.94492303 |
| 1 | 0.1 | 4 | 0.08 | 1 | 0.07143 | 1 | 1 | 1 | ERBB3 | 1 | 1 | 1 |
| 1 | 0.1 | 3 | 0.06 | 0 | 0 | 0.561369203 | 1 | 0.447799719 | ERBB4 | 1 | 1 | 1 |
| 0 | 0 | 9 | 0.18 | 3 | 0.21429 | 0.356969362 | 0.7153727 | 0.199914832 | ERCC1 | 0.9763492 | 1 | 0.94492303 |
| 0 | 0 | 2 | 0.04 | 2 | 0.14286 | 0.241977845 | 0.2056672 | 1 | ERCC2 | 0.9091937 | 1 | 1 |
| 1 | 0.1 | 7 | 0.14 | 2 | 0.14286 | 1 | 1 | 1 | ERCC3 | 1 | 1 | 1 |
| 0 | 0 | 2 | 0.04 | 1 | 0.07143 | 0.697642848 | 0.5295699 | 1 | ERCC4 | 1 | 1 | 1 |
| 0 | 0 | 25 | 0.5 | 5 | 0.35714 | 0.007213584 | 0.3813873 | 0.004371918 | ERCC5 | 0.1730076 | 1 | 0.248677373 |
| 0 | 0 | 14 | 0.28 | 7 | 0.5 | 0.022479606 | 0.1961374 | 0.053464503 | ERCC6 | 0.3148752 | 1 | 0.847858997 |
| 2 | 0.2 | 8 | 0.16 | 0 | 0 | 0.222382945 | 0.1831159 | 0.616643684 | ERG | 0.9091937 | 1 | 1 |
| 0 | 0 | 9 | 0.18 | 6 | 0.42857 | 0.026177212 | 0.0749201 | 0.197470798 | ESR1 | 0.3148752 | 1 | 0.94492303 |
| 1 | 0.1 | 18 | 0.36 | 4 | 0.28571 | 0.287153715 | 0.754635 | 0.158337716 | ETV1 | 0.9763492 | 1 | 0.94492303 |
| 0 | 0 | 11 | 0.22 | 3 | 0.21429 | 0.315033422 | 1 | 0.192132251 | ETV4 | 0.9763492 | 1 | 0.94492303 |
| 0 | 0 | 6 | 0.12 | 5 | 0.35714 | 0.039930204 | 0.0524849 | 0.341106332 | ETV5 | 0.4059571 | 1 | 0.94492303 |
| 1 | 0.1 | 13 | 0.26 | 3 | 0.21429 | 0.647825713 | 1 | 0.436934283 | ETV6 | 1 | 1 | 1 |
| 0 | 0 | 10 | 0.2 | 1 | 0.07143 | 0.250143775 | 0.43058 | 0.341106332 | EWSR1 | 0.9091937 | 1 | 0.94492303 |
| 0 | 0 | 3 | 0.06 | 1 | 0.07143 | 1 | 1 | 1 | EXO1 | 1 | 1 | 1 |
| 1 | 0.1 | 18 | 0.36 | 5 | 0.35714 | 0.31353033 | 1 | 0.151936269 | EXT1 | 0.9763492 | 1 | 0.94492303 |
| 0 | 0 | 5 | 0.1 | 1 | 0.07143 | 0.825954855 | 1 | 0.588422151 | EXT2 | 1 | 1 | 1 |
| 1 | 0.1 | 17 | 0.34 | 5 | 0.35714 | 0.357763426 | 1 | 0.158337716 | EZH2 | 0.9763492 | 1 | 0.94492303 |
| 0 | 0 | 6 | 0.12 | 1 | 0.07143 | 0.835169009 | 1 | 0.583378305 | FAAP100 | 1 | 1 | 1 |
| 0 | 0 | 2 | 0.04 | 0 | 0 | 1 | 1 | 1 | FAAP20 | 1 | 1 | 1 |
| 0 | 0 | 8 | 0.16 | 5 | 0.35714 | 0.053606445 | 0.1363359 | 0.192779347 | FAH | 0.4380924 | 1 | 0.94492303 |
| 0 | 0 | 2 | 0.04 | 1 | 0.07143 | 0.697642848 | 0.5295699 | 1 | FAN1 | 1 | 1 | 1 |
| 0 | 0 | 4 | 0.08 | 1 | 0.07143 | 1 | 1 | 1 | FANCA | 1 | 1 | 1 |
| 0 | 0 | 1 | 0.02 | 0 | 0 | 1 | 1 | 1 | FANCB | 1 | 1 | 1 |
| 0 | 0 | 9 | 0.18 | 3 | 0.21429 | 0.356969362 | 0.7153727 | 0.199914832 | FANCC | 0.9763492 | 1 | 0.94492303 |
| 0 | 0 | 6 | 0.12 | 1 | 0.07143 | 0.835169009 | 1 | 0.583378305 | FANCD2 | 1 | 1 | 1 |
| 0 | 0 | 8 | 0.16 | 3 | 0.21429 | 0.416615438 | 0.6933892 | 0.341106332 | FANCE | 1 | 1 | 0.94492303 |
| 0 | 0 | 8 | 0.16 | 1 | 0.07143 | 0.421541949 | 0.6699605 | 0.346600866 | FANCF | 1 | 1 | 0.94492303 |

| | | | | | | | | | | | | |
|---|---|---|---|---|---|---|---|---|---|---|---|---|
| 0 | 0 | 6 | 0.12 | 0 | 0 | 0.342051159 | 0.3248367 | 0.588422151 | FANCG | 0.9763492 | 1 | 1 |
| 0 | 0 | 1 | 0.02 | 0 | 0 | 1 | 1 | 1 | FANCI | 1 | 1 | 1 |
| 0 | 0 | 1 | 0.02 | 0 | 0 | 1 | 1 | 1 | FANCL | 1 | 1 | 1 |
| 0 | 0 | 6 | 0.12 | 2 | 0.14286 | 0.724461627 | 1 | 0.587819901 | FANCM | 1 | 1 | 1 |
| 0 | 0 | 11 | 0.22 | 0 | 0 | 0.048866885 | 0.1027189 | 0.341106332 | FAS | 0.4375048 | 1 | 0.94492303 |
| 0 | 0 | 4 | 0.08 | 0 | 0 | 0.76152112 | 0.5681297 | 1 | FAT1 | 1 | 1 | 1 |
| 0 | 0 | 7 | 0.14 | 0 | 0 | 0.319022773 | 0.3315098 | 0.583378305 | FBXW7 | 0.9763492 | 1 | 1 |
| 0 | 0 | 18 | 0.36 | 2 | 0.14286 | 0.02903574 | 0.192875 | 0.05331369 | FGFR1 | 0.3373676 | 1 | 0.847858997 |
| 0 | 0 | 10 | 0.2 | 0 | 0 | 0.073759985 | 0.100854 | 0.33939492 | FGFR2 | 0.50697 | 1 | 0.94492303 |
| 0 | 0 | 5 | 0.1 | 0 | 0 | 0.48654037 | 0.577127 | 1 | FGFR3 | 1 | 1 | 1 |
| 0 | 0 | 2 | 0.04 | 0 | 0 | 1 | 1 | 1 | FGFR4 | 1 | 1 | 1 |
| 0 | 0 | 5 | 0.1 | 2 | 0.14286 | 0.595089088 | 0.6418802 | 0.583378305 | FH | 1 | 1 | 1 |
| 1 | 0.1 | 13 | 0.26 | 4 | 0.28571 | 0.603863443 | 1 | 0.434053099 | FKBP9 | 1 | 1 | 1 |
| 0 | 0 | 21 | 0.42 | 2 | 0.14286 | 0.00674495 | 0.066506 | 0.025822519 | FLCN | 0.1730076 | 1 | 0.618749838 |
| 0 | 0 | 23 | 0.46 | 3 | 0.21429 | 0.006656455 | 0.1293853 | 0.011773622 | FLT1 | 0.1730076 | 1 | 0.383035181 |
| 0 | 0 | 24 | 0.48 | 4 | 0.28571 | 0.006669232 | 0.235397 | 0.01061248 | FLT3 | 0.1730076 | 1 | 0.383035181 |
| 0 | 0 | 2 | 0.04 | 0 | 0 | 1 | 1 | 1 | FLT4 | 1 | 1 | 1 |
| 0 | 0 | 3 | 0.06 | 0 | 0 | 1 | 1 | 1 | FOXA1 | 1 | 1 | 1 |
| 0 | 0 | 2 | 0.04 | 2 | 0.14286 | 0.241977845 | 0.2056672 | 1 | FOXL2 | 0.9091937 | 1 | 1 |
| 0 | 0 | 3 | 0.06 | 3 | 0.21429 | 0.133673095 | 0.1128883 | 0.588422151 | FUS | 0.6919557 | 1 | 1 |
| 0 | 0 | 2 | 0.04 | 0 | 0 | 1 | 1 | 1 | GALNT12 | 1 | 1 | 1 |
| 0 | 0 | 2 | 0.04 | 1 | 0.07143 | 0.697642848 | 0.5295699 | 1 | GATA2 | 1 | 1 | 1 |
| 0 | 0 | 10 | 0.2 | 5 | 0.35714 | 0.103692008 | 0.2861485 | 0.197470798 | GATA3 | 0.6519717 | 1 | 0.94492303 |
| 0 | 0 | 20 | 0.4 | 3 | 0.21429 | 0.026454681 | 0.3446684 | 0.025822519 | GATA4 | 0.3148752 | 1 | 0.618749838 |
| 1 | 0.1 | 22 | 0.44 | 2 | 0.14286 | 0.024890864 | 0.0610259 | 0.149405814 | GATA6 | 0.3148752 | 1 | 0.94492303 |
| 0 | 0 | 3 | 0.06 | 1 | 0.07143 | 1 | 1 | 1 | GBA | 1 | 1 | 1 |
| 0 | 0 | 9 | 0.18 | 6 | 0.42857 | 0.026177212 | 0.0749201 | 0.197470798 | GEN1 | 0.3148752 | 1 | 0.94492303 |
| 1 | 0.1 | 7 | 0.14 | 3 | 0.21429 | 0.785257333 | 0.6774614 | 1 | GLI1 | 1 | 1 | 1 |
| 1 | 0.1 | 4 | 0.08 | 0 | 0 | 0.629506025 | 0.5681297 | 0.526685474 | GLI2 | 1 | 1 | 1 |
| 1 | 0.1 | 12 | 0.24 | 3 | 0.21429 | 0.699616049 | 1 | 0.679825377 | GLI3 | 1 | 1 | 1 |
| 0 | 0 | 2 | 0.04 | 1 | 0.07143 | 0.697642848 | 0.5295699 | 1 | GNA11 | 1 | 1 | 1 |
| 0 | 0 | 7 | 0.14 | 0 | 0 | 0.319022773 | 0.3315098 | 0.583378305 | GNAQ | 0.9763492 | 1 | 1 |
| 0 | 0 | 25 | 0.5 | 4 | 0.28571 | 0.005449123 | 0.2262717 | 0.005095848 | GNAS | 0.1730076 | 1 | 0.248677373 |
| 0 | 0 | 7 | 0.14 | 1 | 0.07143 | 0.62851523 | 0.6732942 | 0.587819901 | GPC3 | 1 | 1 | 1 |
| 0 | 0 | 2 | 0.04 | 0 | 0 | 1 | 1 | 1 | GREM1 | 1 | 1 | 1 |
| 1 | 0.1 | 13 | 0.26 | 5 | 0.35714 | 0.36729587 | 0.5118719 | 0.436474262 | GSTM5 | 0.9957799 | 1 | 1 |
| 0 | 0 | 4 | 0.08 | 1 | 0.07143 | 1 | 1 | 1 | H19 | 1 | 1 | 1 |
| 0 | 0 | 11 | 0.22 | 7 | 0.5 | 0.015855952 | 0.0507235 | 0.105937061 | H3F3A | 0.2865816 | 1 | 0.94492303 |
| 0 | 0 | 4 | 0.08 | 1 | 0.07143 | 1 | 1 | 1 | H3F3B | 1 | 1 | 1 |
| 0 | 0 | 3 | 0.06 | 0 | 0 | 1 | 1 | 1 | HABP2 | 1 | 1 | 1 |
| 0 | 0 | 5 | 0.1 | 0 | 0 | 0.48654037 | 0.577127 | 1 | HELQ | 1 | 1 | 1 |
| 0 | 0 | 5 | 0.1 | 1 | 0.07143 | 0.825954855 | 1 | 0.588422151 | HFE | 1 | 1 | 1 |
| 0 | 0 | 5 | 0.1 | 1 | 0.07143 | 0.825954855 | 1 | 0.588422151 | HIST1H3B | 1 | 1 | 1 |
| 0 | 0 | 6 | 0.12 | 4 | 0.28571 | 0.12702587 | 0.2058665 | 0.33939492 | HIST1H3C | 0.6919557 | 1 | 0.94492303 |
| 0 | 0 | 3 | 0.06 | 3 | 0.21429 | 0.133673095 | 0.1128883 | 0.588422151 | HMBS | 0.6919557 | 1 | 1 |
| 1 | 0.1 | 8 | 0.16 | 3 | 0.21429 | 0.795914644 | 0.6933892 | 1 | HNF1A | 1 | 1 | 1 |
| 0 | 0 | 6 | 0.12 | 2 | 0.14286 | 0.724461627 | 1 | 0.587819901 | HOXB13 | 1 | 1 | 1 |
| 0 | 0 | 7 | 0.14 | 3 | 0.21429 | 0.442160876 | 0.6774614 | 0.33939492 | HRAS | 1 | 1 | 0.94492303 |
| 0 | 0 | 8 | 0.16 | 1 | 0.07143 | 0.421541949 | 0.6699605 | 0.346600866 | ID3 | 1 | 1 | 0.94492303 |
| 0 | 0 | 4 | 0.08 | 1 | 0.07143 | 1 | 1 | 1 | ID4 | 1 | 1 | 1 |
| 1 | 0.1 | 4 | 0.08 | 0 | 0 | 0.629506025 | 0.5681297 | 0.526685474 | IDH1 | 1 | 1 | 1 |
| 0 | 0 | 6 | 0.12 | 1 | 0.07143 | 0.835169009 | 1 | 0.583378305 | IDH2 | 1 | 1 | 1 |
| 0 | 0 | 8 | 0.16 | 1 | 0.07143 | 0.421541949 | 0.6699605 | 0.346600866 | IGF1R | 1 | 1 | 0.94492303 |
| 0 | 0 | 6 | 0.12 | 2 | 0.14286 | 0.724461627 | 1 | 0.587819901 | IGF2 | 1 | 1 | 1 |
| 1 | 0.1 | 16 | 0.32 | 4 | 0.28571 | 0.4626501 | 1 | 0.264290836 | IKZF1 | 1 | 1 | 0.94492303 |
| 0 | 0 | 12 | 0.24 | 5 | 0.35714 | 0.120634886 | 0.4949933 | 0.103827959 | IKZF3 | 0.6919557 | 1 | 0.94492303 |
| 0 | 0 | 1 | 0.02 | 1 | 0.07143 | 0.546464272 | 0.3923611 | 1 | IL7R | 1 | 1 | 1 |
| 1 | 0.1 | 11 | 0.22 | 3 | 0.21429 | 0.823891618 | 1 | 0.675901668 | INSIG1 | 1 | 1 | 1 |
| 0 | 0 | 13 | 0.26 | 5 | 0.35714 | 0.099207853 | 0.5118719 | 0.105937061 | ITK | 0.6519717 | 1 | 0.94492303 |
| 0 | 0 | 4 | 0.08 | 2 | 0.14286 | 0.551460797 | 0.6043629 | 0.588422151 | JAK1 | 1 | 1 | 1 |
| 0 | 0 | 9 | 0.18 | 2 | 0.14286 | 0.471161887 | 1 | 0.341106332 | JAK2 | 1 | 1 | 0.94492303 |
| 0 | 0 | 1 | 0.02 | 0 | 0 | 1 | 1 | 1 | JAK3 | 1 | 1 | 1 |
| 0 | 0 | 5 | 0.1 | 1 | 0.07143 | 0.825954855 | 1 | 0.588422151 | JAZF1 | 1 | 1 | 1 |
| 0 | 0 | 6 | 0.12 | 1 | 0.07143 | 0.835169009 | 1 | 0.583378305 | KAT6A | 1 | 1 | 1 |
| 0 | 0 | 2 | 0.04 | 0 | 0 | 1 | 1 | 1 | KAT6B | 1 | 1 | 1 |
| 0 | 0 | 1 | 0.02 | 0 | 0 | 1 | 1 | 1 | KCNIP1 | 1 | 1 | 1 |
| 0 | 0 | 4 | 0.08 | 1 | 0.07143 | 1 | 1 | 1 | KCNQ1 | 1 | 1 | 1 |
| 0 | 0 | 2 | 0.04 | 1 | 0.07143 | 0.697642848 | 0.5295699 | 1 | KDM5A | 1 | 1 | 1 |
| 0 | 0 | 8 | 0.16 | 4 | 0.28571 | 0.210057837 | 0.4376874 | 0.199914832 | KDM5C | 0.8958294 | 1 | 0.94492303 |
| 0 | 0 | 2 | 0.04 | 0 | 0 | 1 | 1 | 1 | KDM6A | 1 | 1 | 1 |
| 0 | 0 | 11 | 0.22 | 2 | 0.14286 | 0.26168066 | 0.7153727 | 0.192779347 | KDM6B | 0.9091937 | 1 | 0.94492303 |
| 0 | 0 | 6 | 0.12 | 0 | 0 | 0.342051159 | 0.3248367 | 0.588422151 | KDR | 0.9763492 | 1 | 1 |
| 0 | 0 | 1 | 0.02 | 1 | 0.07143 | 0.546464272 | 0.3923611 | 1 | KEAP1 | 1 | 1 | 1 |
| 0 | 0 | 2 | 0.04 | 1 | 0.07143 | 0.697642848 | 0.5295699 | 1 | KIF1B | 1 | 1 | 1 |
| 0 | 0 | 6 | 0.12 | 0 | 0 | 0.342051159 | 0.3248367 | 0.588422151 | KIT | 0.9763492 | 1 | 1 |
| 0 | 0 | 2 | 0.04 | 0 | 0 | 1 | 1 | 1 | KLF4 | 1 | 1 | 1 |
| 0 | 0 | 3 | 0.06 | 0 | 0 | 1 | 1 | 1 | KLLN | 1 | 1 | 1 |
| 0 | 0 | 7 | 0.14 | 1 | 0.07143 | 0.62851523 | 0.6732942 | 0.587819901 | KMT2A | 1 | 1 | 1 |
| 1 | 0.1 | 5 | 0.1 | 2 | 0.14286 | 0.852390158 | 0.6418802 | 1 | KMT2D | 1 | 1 | 1 |

| | | | | | | | | | | | | |
|---|---|---|---|---|---|---|---|---|---|---|---|---|
| 1 | 0.1 | 9 | 0.18 | 2 | 0.14286 | 1 | 1 | 1 | KRAS | 1 | 1 | 1 |
| 0 | 0 | 8 | 0.16 | 1 | 0.07143 | 0.421541949 | 0.6699605 | 0.346600866 | LIG4 | 1 | 1 | 0.94492303 |
| 0 | 0 | 1 | 0.02 | 0 | 0 | 1 | 1 | 1 | LINC00894 | 1 | 1 | 1 |
| 0 | 0 | 7 | 0.14 | 3 | 0.21429 | 0.442160876 | 0.6774614 | 0.33939492 | LMO1 | 1 | 1 | 0.94492303 |
| 0 | 0 | 8 | 0.16 | 3 | 0.21429 | 0.416615438 | 0.6933892 | 0.341106332 | LMO2 | 1 | 1 | 0.94492303 |
| 1 | 0.1 | 7 | 0.14 | 2 | 0.14286 | 1 | 1 | 1 | LMO3 | 1 | 1 | 1 |
| 0 | 0 | 1 | 0.02 | 2 | 0.14286 | 0.136122424 | 0.1179435 | 1 | MAF | 0.6919557 | 1 | 1 |
| 0 | 0 | 15 | 0.3 | 6 | 0.42857 | 0.045127575 | 0.5204229 | 0.053464503 | MAFB | 0.431809 | 1 | 0.847858997 |
| 0 | 0 | 6 | 0.12 | 1 | 0.07143 | 0.835169009 | 1 | 0.583378305 | MAP2K1 | 1 | 1 | 1 |
| 0 | 0 | 21 | 0.42 | 2 | 0.14286 | 0.00674495 | 0.066506 | 0.025822519 | MAP2K4 | 0.1730076 | 1 | 0.618749838 |
| 0 | 0 | 5 | 0.1 | 2 | 0.14286 | 0.595089088 | 0.6418802 | 0.583378305 | MAP3K1 | 1 | 1 | 1 |
| 0 | 0 | 10 | 0.2 | 0 | 0 | 0.073759985 | 0.100854 | 0.33939492 | MAPK1 | 0.50697 | 1 | 0.94492303 |
| 0 | 0 | 6 | 0.12 | 1 | 0.07143 | 0.835169009 | 1 | 0.583378305 | MAX | 1 | 1 | 1 |
| 0 | 0 | 5 | 0.1 | 3 | 0.21429 | 0.26269735 | 0.357359 | 0.587819901 | MBD4 | 0.9091937 | 1 | 1 |
| 0 | 0 | 5 | 0.1 | 1 | 0.07143 | 0.825954855 | 1 | 0.588422151 | MCL1 | 1 | 1 | 1 |
| 0 | 0 | 9 | 0.18 | 2 | 0.14286 | 0.471161887 | 1 | 0.341106332 | MCM8 | 1 | 1 | 0.94492303 |
| 1 | 0.1 | 11 | 0.22 | 1 | 0.07143 | 0.472274771 | 0.2740515 | 0.68026957 | MDM2 | 1 | 1 | 1 |
| 0 | 0 | 3 | 0.06 | 1 | 0.07143 | 1 | 1 | 1 | MDM4 | 1 | 1 | 1 |
| 0 | 0 | 2 | 0.04 | 1 | 0.07143 | 0.697642848 | 0.5295699 | 1 | MECOM | 1 | 1 | 1 |
| 0 | 0 | 1 | 0.02 | 0 | 0 | 1 | 1 | 1 | MED12 | 1 | 1 | 1 |
| 0 | 0 | 2 | 0.04 | 1 | 0.07143 | 0.697642848 | 0.5295699 | 1 | MEF2B | 1 | 1 | 1 |
| 0 | 0 | 5 | 0.1 | 1 | 0.07143 | 0.825954855 | 1 | 0.588422151 | MEN1 | 1 | 1 | 1 |
| 1 | 0.1 | 16 | 0.32 | 4 | 0.28571 | 0.4626501 | 1 | 0.264290836 | MET | 1 | 1 | 0.94492303 |
| 0 | 0 | 2 | 0.04 | 1 | 0.07143 | 0.697642848 | 0.5295699 | 1 | MGA | 1 | 1 | 1 |
| 0 | 0 | 5 | 0.1 | 4 | 0.28571 | 0.079340314 | 0.0960457 | 0.346600866 | MITF | 0.5303846 | 1 | 0.94492303 |
| 0 | 0 | 8 | 0.16 | 3 | 0.21429 | 0.416615438 | 0.6933892 | 0.341106332 | MLH1 | 1 | 1 | 0.94492303 |
| 0 | 0 | 12 | 0.24 | 6 | 0.42857 | 0.053863824 | 0.190221 | 0.105937061 | MLH3 | 0.4380924 | 1 | 0.94492303 |
| 0 | 0 | 4 | 0.08 | 0 | 0 | 0.76152112 | 0.5681297 | 1 | MPL | 1 | 1 | 1 |
| 0 | 0 | 3 | 0.06 | 1 | 0.07143 | 1 | 1 | 1 | MRE11 | 1 | 1 | 1 |
| 0 | 0 | 9 | 0.18 | 6 | 0.42857 | 0.026177212 | 0.0749201 | 0.197470798 | MSH2 | 0.3148752 | 1 | 0.94492303 |
| 0 | 0 | 2 | 0.04 | 0 | 0 | 1 | 1 | 1 | MSH6 | 1 | 1 | 1 |
| 0 | 0 | 6 | 0.12 | 1 | 0.07143 | 0.835169009 | 1 | 0.583378305 | MTA1 | 1 | 1 | 1 |
| 0 | 0 | 1 | 0.02 | 1 | 0.07143 | 0.546464272 | 0.3923611 | 1 | MTAP | 1 | 1 | 1 |
| 0 | 0 | 9 | 0.18 | 2 | 0.14286 | 0.471161887 | 1 | 0.341106332 | MTOR | 1 | 1 | 0.94492303 |
| 0 | 0 | 2 | 0.04 | 1 | 0.07143 | 0.697642848 | 0.5295699 | 1 | MUS81 | 1 | 1 | 1 |
| 0 | 0 | 4 | 0.08 | 0 | 0 | 0.76152112 | 0.5681297 | 1 | MUTYH | 1 | 1 | 1 |
| 0 | 0 | 9 | 0.18 | 4 | 0.28571 | 0.229201109 | 0.4566205 | 0.192779347 | MYB | 0.9091937 | 1 | 0.94492303 |
| 1 | 0.1 | 17 | 0.34 | 2 | 0.14286 | 0.193259725 | 0.1985534 | 0.269612096 | MYBL1 | 0.8814088 | 1 | 0.94492303 |
| 1 | 0.1 | 32 | 0.64 | 11 | 0.78571 | 0.001520915 | 0.3559965 | 0.000918086 | MYC | 0.1730076 | 1 | 0.248677373 |
| 0 | 0 | 4 | 0.08 | 3 | 0.21429 | 0.21053707 | 0.1707213 | 0.583378305 | MYCL | 0.8958294 | 1 | 1 |
| 0 | 0 | 5 | 0.1 | 1 | 0.07143 | 0.825954855 | 1 | 0.588422151 | MYCN | 1 | 1 | 1 |
| 0 | 0 | 4 | 0.08 | 1 | 0.07143 | 1 | 1 | 1 | MYD88 | 1 | 1 | 1 |
| 1 | 0.1 | 17 | 0.34 | 2 | 0.14286 | 0.193259725 | 0.1985534 | 0.269612096 | NBN | 0.8814088 | 1 | 0.94492303 |
| 0 | 0 | 5 | 0.1 | 0 | 0 | 0.48654037 | 0.577127 | 1 | NECTIN4 | 1 | 1 | 1 |
| 0 | 0 | 5 | 0.1 | 0 | 0 | 0.48654037 | 0.577127 | 1 | NEGR1 | 1 | 1 | 1 |
| 0 | 0 | 3 | 0.06 | 0 | 0 | 1 | 1 | 1 | NEIL1 | 1 | 1 | 1 |
| 0 | 0 | 9 | 0.18 | 2 | 0.14286 | 0.471161887 | 1 | 0.341106332 | NEIL2 | 1 | 1 | 0.94492303 |
| 0 | 0 | 5 | 0.1 | 3 | 0.21429 | 0.26269735 | 0.357359 | 0.587819901 | NEIL3 | 0.9091937 | 1 | 1 |
| 0 | 0 | 13 | 0.26 | 2 | 0.14286 | 0.162620262 | 0.488584 | 0.197470798 | NF1 | 0.8012382 | 1 | 0.94492303 |
| 0 | 0 | 9 | 0.18 | 2 | 0.14286 | 0.471161887 | 1 | 0.341106332 | NF2 | 1 | 1 | 0.94492303 |
| 1 | 0.1 | 4 | 0.08 | 1 | 0.07143 | 1 | 1 | 1 | NFE2L2 | 1 | 1 | 1 |
| 0 | 0 | 4 | 0.08 | 1 | 0.07143 | 1 | 1 | 1 | NFKBIA | 1 | 1 | 1 |
| 0 | 0 | 5 | 0.1 | 3 | 0.21429 | 0.26269735 | 0.357359 | 0.587819901 | NFKBIE | 0.9091937 | 1 | 1 |
| 0 | 0 | 7 | 0.14 | 2 | 0.14286 | 0.652062514 | 1 | 0.346600866 | NFKBIZ | 1 | 1 | 0.94492303 |
| 0 | 0 | 6 | 0.12 | 1 | 0.07143 | 0.835169009 | 1 | 0.583378305 | NKX2-1 | 1 | 1 | 1 |
| 0 | 0 | 9 | 0.18 | 1 | 0.07143 | 0.391551787 | 0.4374327 | 0.33939492 | NKX3-1 | 1 | 1 | 0.94492303 |
| 0 | 0 | 9 | 0.18 | 1 | 0.07143 | 0.391551787 | 0.4374327 | 0.33939492 | NOTCH1 | 1 | 1 | 0.94492303 |
| 0 | 0 | 4 | 0.08 | 0 | 0 | 0.76152112 | 0.5681297 | 1 | NOTCH2 | 1 | 1 | 1 |
| 0 | 0 | 3 | 0.06 | 2 | 0.14286 | 0.355011967 | 0.2992391 | 1 | NOTCH3 | 0.9763492 | 1 | 1 |
| 0 | 0 | 9 | 0.18 | 6 | 0.42857 | 0.026177212 | 0.0749201 | 0.197470798 | NPM1 | 0.3148752 | 1 | 0.94492303 |
| 0 | 0 | 5 | 0.1 | 2 | 0.14286 | 0.595089088 | 0.6418802 | 0.583378305 | NPRL2 | 1 | 1 | 1 |
| 0 | 0 | 3 | 0.06 | 0 | 0 | 1 | 1 | 1 | NPRL3 | 1 | 1 | 1 |
| 0 | 0 | 1 | 0.02 | 0 | 0 | 1 | 1 | 1 | NR0B1 | 1 | 1 | 1 |
| 1 | 0.1 | 6 | 0.12 | 0 | 0 | 0.487948944 | 0.3248367 | 1 | NRAS | 1 | 1 | 1 |
| 0 | 0 | 9 | 0.18 | 1 | 0.07143 | 0.391551787 | 0.4374327 | 0.33939492 | NRG1 | 1 | 1 | 0.94492303 |
| 0 | 0 | 1 | 0.02 | 0 | 0 | 1 | 1 | 1 | NSD1 | 1 | 1 | 1 |
| 0 | 0 | 5 | 0.1 | 0 | 0 | 0.48654037 | 0.577127 | 1 | NSD2 | 1 | 1 | 1 |
| 0 | 0 | 7 | 0.14 | 0 | 0 | 0.319022773 | 0.3315098 | 0.583378305 | NSD3 | 0.9763492 | 1 | 1 |
| 0 | 0 | 3 | 0.06 | 0 | 0 | 1 | 1 | 1 | NT5C2 | 1 | 1 | 1 |
| 0 | 0 | 1 | 0.02 | 2 | 0.14286 | 0.136122424 | 0.1179435 | 1 | NTHL1 | 0.6919557 | 1 | 1 |
| 0 | 0 | 5 | 0.1 | 1 | 0.07143 | 0.825954855 | 1 | 0.588422151 | NTRK1 | 1 | 1 | 1 |
| 0 | 0 | 6 | 0.12 | 0 | 0 | 0.342051159 | 0.3248367 | 0.588422151 | NTRK2 | 0.9763492 | 1 | 1 |
| 0 | 0 | 7 | 0.14 | 1 | 0.07143 | 0.62851523 | 0.6732942 | 0.587819901 | NTRK3 | 1 | 1 | 1 |
| 0 | 0 | 2 | 0.04 | 2 | 0.14286 | 0.241977845 | 0.2056672 | 1 | OGG1 | 0.9091937 | 1 | 1 |
| 0 | 0 | 3 | 0.06 | 1 | 0.07143 | 1 | 1 | 1 | PALB2 | 1 | 1 | 1 |
| 0 | 0 | 9 | 0.18 | 0 | 0 | 0.122866881 | 0.1870181 | 0.346600866 | PAX5 | 0.6919557 | 1 | 0.94492303 |
| 0 | 0 | 6 | 0.12 | 1 | 0.07143 | 0.835169009 | 1 | 0.583378305 | PAXIP1 | 1 | 1 | 1 |
| 0 | 0 | 4 | 0.08 | 1 | 0.07143 | 1 | 1 | 1 | PBRM1 | 1 | 1 | 1 |
| 0 | 0 | 9 | 0.18 | 1 | 0.07143 | 0.391551787 | 0.4374327 | 0.33939492 | PDCD1LG2 | 1 | 1 | 0.94492303 |

| | | | | | | | | | | | | |
|---|---|---|---|---|---|---|---|---|---|---|---|---|
| 0 | 0 | 6 | 0.12 | 0 | 0 | 0.342051159 | 0.3248367 | 0.588422151 | PDGFRA | 0.9763492 | 1 | 1 |
| 0 | 0 | 2 | 0.04 | 3 | 0.21429 | 0.072426662 | 0.0653094 | 1 | PDGFRB | 0.50697 | 1 | 1 |
| 0 | 0 | 2 | 0.04 | 0 | 0 | 1 | 1 | 1 | PHF6 | 1 | 1 | 1 |
| 0 | 0 | 10 | 0.2 | 3 | 0.21429 | 0.297408165 | 1 | 0.192779347 | PHOX2B | 0.9763492 | 1 | 0.94492303 |
| 0 | 0 | 14 | 0.28 | 6 | 0.42857 | 0.051998521 | 0.3364219 | 0.05331369 | PIK3C2B | 0.4375048 | 1 | 0.847858997 |
| 0 | 0 | 2 | 0.04 | 1 | 0.07143 | 0.697642848 | 0.5295699 | 1 | PIK3CA | 1 | 1 | 1 |
| 1 | 0.1 | 3 | 0.06 | 1 | 0.07143 | 0.799848083 | 1 | 0.526685474 | PIK3R1 | 1 | 1 | 1 |
| 0 | 0 | 9 | 0.18 | 2 | 0.14286 | 0.471161887 | 1 | 0.341106332 | PIM1 | 1 | 1 | 0.94492303 |
| 0 | 0 | 2 | 0.04 | 1 | 0.07143 | 0.697642848 | 0.5295699 | 1 | PML | 1 | 1 | 1 |
| 2 | 0.2 | 3 | 0.06 | 0 | 0 | 0.175088169 | 1 | 0.132256702 | PMS1 | 0.8459706 | 1 | 0.94492303 |
| 2 | 0.2 | 18 | 0.36 | 4 | 0.28571 | 0.701407187 | 0.754635 | 0.483550933 | PMS2 | 1 | 1 | 1 |
| 0 | 0 | 3 | 0.06 | 0 | 0 | 1 | 1 | 1 | PNKP | 1 | 1 | 1 |
| 0 | 0 | 5 | 0.1 | 1 | 0.07143 | 0.825954855 | 1 | 0.588422151 | PNRC1 | 1 | 1 | 1 |
| 0 | 0 | 7 | 0.14 | 2 | 0.14286 | 0.652062514 | 1 | 0.346600866 | POLB | 1 | 1 | 0.94492303 |
| 0 | 0 | 2 | 0.04 | 0 | 0 | 1 | 1 | 1 | POLD1 | 1 | 1 | 1 |
| 0 | 0 | 4 | 0.08 | 0 | 0 | 0.76152112 | 0.5681297 | 1 | POLE | 1 | 1 | 1 |
| 0 | 0 | 4 | 0.08 | 1 | 0.07143 | 1 | 1 | 1 | POLH | 1 | 1 | 1 |
| 0 | 0 | 2 | 0.04 | 1 | 0.07143 | 0.697642848 | 0.5295699 | 1 | POLQ | 1 | 1 | 1 |
| 0 | 0 | 10 | 0.2 | 4 | 0.28571 | 0.211107347 | 0.4850325 | 0.192132251 | POT1 | 0.8958294 | 1 | 0.94492303 |
| 0 | 0 | 4 | 0.08 | 1 | 0.07143 | 1 | 1 | 1 | PPARG | 1 | 1 | 1 |
| 0 | 0 | 5 | 0.1 | 5 | 0.35714 | 0.024244074 | 0.0330382 | 0.33939492 | PPM1D | 0.3148752 | 1 | 0.94492303 |
| 0 | 0 | 3 | 0.06 | 0 | 0 | 1 | 1 | 1 | PPP2R1A | 1 | 1 | 1 |
| 0 | 0 | 6 | 0.12 | 2 | 0.14286 | 0.724461627 | 1 | 0.587819901 | PRAME | 1 | 1 | 1 |
| 0 | 0 | 9 | 0.18 | 4 | 0.28571 | 0.229201109 | 0.4566205 | 0.192779347 | PRDM1 | 0.9091937 | 1 | 0.94492303 |
| 0 | 0 | 6 | 0.12 | 2 | 0.14286 | 0.724461627 | 1 | 0.587819901 | PRF1 | 1 | 1 | 1 |
| 0 | 0 | 16 | 0.32 | 6 | 0.42857 | 0.04165357 | 0.529717 | 0.02789446 | PRKAR1A | 0.4119566 | 1 | 0.618749838 |
| 0 | 0 | 5 | 0.1 | 4 | 0.28571 | 0.079340314 | 0.0960457 | 0.346600866 | PRKCI | 0.5303846 | 1 | 0.94492303 |
| 0 | 0 | 4 | 0.08 | 0 | 0 | 0.76152112 | 0.5681297 | 1 | PRKCZ | 1 | 1 | 1 |
| 1 | 0.1 | 16 | 0.32 | 2 | 0.14286 | 0.278895204 | 0.3146226 | 0.436474262 | PRKDC | 0.9584568 | 1 | 1 |
| 0 | 0 | 9 | 0.18 | 2 | 0.14286 | 0.471161887 | 1 | 0.341106332 | PRKN | 1 | 1 | 0.94492303 |
| 1 | 0.1 | 3 | 0.06 | 2 | 0.14286 | 0.438331452 | 0.2992391 | 1 | PRPF40B | 1 | 1 | 1 |
| 0 | 0 | 9 | 0.18 | 3 | 0.21429 | 0.356969362 | 0.7153727 | 0.199914832 | PRPF8 | 0.9763492 | 1 | 0.94492303 |
| 0 | 0 | 5 | 0.1 | 1 | 0.07143 | 0.825954855 | 1 | 0.588422151 | PRSS1 | 1 | 1 | 1 |
| 0 | 0 | 3 | 0.06 | 2 | 0.14286 | 0.355011967 | 0.2992391 | 1 | PSMD13 | 0.9763492 | 1 | 1 |
| 0 | 0 | 7 | 0.14 | 2 | 0.14286 | 0.652062514 | 1 | 0.346600866 | PTCH1 | 1 | 1 | 0.94492303 |
| 0 | 0 | 11 | 0.22 | 0 | 0 | 0.048866885 | 0.1027189 | 0.341106332 | PTEN | 0.4375048 | 1 | 0.94492303 |
| 1 | 0.1 | 14 | 0.28 | 3 | 0.21429 | 0.554981208 | 0.74248 | 0.434053099 | PTK2 | 1 | 1 | 1 |
| 0 | 0 | 12 | 0.24 | 3 | 0.21429 | 0.229845529 | 1 | 0.197470798 | PTK2B | 0.9091937 | 1 | 0.94492303 |
| 1 | 0.1 | 11 | 0.22 | 5 | 0.35714 | 0.37307475 | 0.312148 | 0.436934283 | PTPN11 | 1 | 1 | 1 |
| 0 | 0 | 10 | 0.2 | 5 | 0.35714 | 0.103692008 | 0.2861485 | 0.197470798 | PTPN14 | 0.6519717 | 1 | 0.94492303 |
| 0 | 0 | 8 | 0.16 | 3 | 0.21429 | 0.416615438 | 0.6933892 | 0.341106332 | PTPRD | 1 | 1 | 0.94492303 |
| 0 | 0 | 6 | 0.12 | 1 | 0.07143 | 0.835169009 | 1 | 0.583378305 | QKI | 1 | 1 | 1 |
| 0 | 0 | 6 | 0.12 | 1 | 0.07143 | 0.835169009 | 1 | 0.583378305 | RAC1 | 1 | 1 | 1 |
| 0 | 0 | 3 | 0.06 | 0 | 0 | 1 | 1 | 1 | RACK1 | 1 | 1 | 1 |
| 1 | 0.1 | 23 | 0.46 | 3 | 0.21429 | 0.039592698 | 0.1293853 | 0.082122838 | RAD21 | 0.4059571 | 1 | 0.94492303 |
| 0 | 0 | 1 | 0.02 | 1 | 0.07143 | 0.546464272 | 0.3923611 | 1 | RAD50 | 1 | 1 | 1 |
| 0 | 0 | 3 | 0.06 | 1 | 0.07143 | 1 | 1 | 1 | RAD51 | 1 | 1 | 1 |
| 0 | 0 | 4 | 0.08 | 0 | 0 | 0.76152112 | 0.5681297 | 1 | RAD51C | 1 | 1 | 1 |
| 0 | 0 | 6 | 0.12 | 1 | 0.07143 | 0.835169009 | 1 | 0.583378305 | RAD51D | 1 | 1 | 1 |
| 0 | 0 | 2 | 0.04 | 2 | 0.14286 | 0.241977845 | 0.2056672 | 1 | RAD52 | 0.9091937 | 1 | 1 |
| 0 | 0 | 6 | 0.12 | 6 | 0.42857 | 0.008374935 | 0.016967 | 0.199914832 | RAD54B | 0.1730076 | 1 | 0.94492303 |
| 0 | 0 | 4 | 0.08 | 1 | 0.07143 | 1 | 1 | 1 | RAF1 | 1 | 1 | 1 |
| 0 | 0 | 11 | 0.22 | 3 | 0.21429 | 0.315033422 | 1 | 0.192132251 | RARA | 0.9763492 | 1 | 0.94492303 |
| 0 | 0 | 3 | 0.06 | 1 | 0.07143 | 1 | 1 | 1 | RASA1 | 1 | 1 | 1 |
| 0 | 0 | 25 | 0.5 | 4 | 0.28571 | 0.005449123 | 0.2262717 | 0.005095848 | RB1 | 0.1730076 | 1 | 0.248677373 |
| 0 | 0 | 9 | 0.18 | 0 | 0 | 0.122866881 | 0.1870181 | 0.346600866 | RBBP8 | 0.6919557 | 1 | 0.94492303 |
| 0 | 0 | 2 | 0.04 | 0 | 0 | 1 | 1 | 1 | RBL2 | 1 | 1 | 1 |
| 0 | 0 | 1 | 0.02 | 0 | 0 | 1 | 1 | 1 | RBM10 | 1 | 1 | 1 |
| 1 | 0.1 | 18 | 0.36 | 3 | 0.21429 | 0.215605768 | 0.3559965 | 0.263944896 | RECQL4 | 0.9070312 | 1 | 0.94492303 |
| 0 | 0 | 5 | 0.1 | 1 | 0.07143 | 0.825954855 | 1 | 0.588422151 | REL | 1 | 1 | 1 |
| 0 | 0 | 2 | 0.04 | 1 | 0.07143 | 0.697642848 | 0.5295699 | 1 | RELA | 1 | 1 | 1 |
| 0 | 0 | 5 | 0.1 | 1 | 0.07143 | 0.825954855 | 1 | 0.588422151 | RET | 1 | 1 | 1 |
| 0 | 0 | 11 | 0.22 | 6 | 0.42857 | 0.046022119 | 0.1703034 | 0.103827959 | RHBDF2 | 0.4318999 | 1 | 0.94492303 |
| 1 | 0.1 | 16 | 0.32 | 4 | 0.28571 | 0.4626501 | 1 | 0.264290836 | RHEB | 1 | 1 | 0.94492303 |
| 0 | 0 | 1 | 0.02 | 1 | 0.07143 | 0.546464272 | 0.3923611 | 1 | RHOA | 1 | 1 | 1 |
| 0 | 0 | 6 | 0.12 | 2 | 0.14286 | 0.724461627 | 1 | 0.587819901 | RHOH | 1 | 1 | 1 |
| 0 | 0 | 7 | 0.14 | 2 | 0.14286 | 0.652062514 | 1 | 0.346600866 | RHOT1 | 1 | 1 | 0.94492303 |
| 0 | 0 | 1 | 0.02 | 1 | 0.07143 | 0.546464272 | 0.3923611 | 1 | RICTOR | 1 | 1 | 1 |
| 0 | 0 | 4 | 0.08 | 1 | 0.07143 | 1 | 1 | 1 | RIF1 | 1 | 1 | 1 |
| 0 | 0 | 5 | 0.1 | 1 | 0.07143 | 0.825954855 | 1 | 0.588422151 | RINT1 | 1 | 1 | 1 |
| 0 | 0 | 3 | 0.06 | 1 | 0.07143 | 1 | 1 | 1 | RIT1 | 1 | 1 | 1 |
| 0 | 0 | 1 | 0.02 | 0 | 0 | 1 | 1 | 1 | RMRP | 1 | 1 | 1 |
| 0 | 0 | 5 | 0.1 | 1 | 0.07143 | 0.825954855 | 1 | 0.588422151 | RNF43 | 1 | 1 | 1 |
| 0 | 0 | 4 | 0.08 | 2 | 0.14286 | 0.551460797 | 0.6043629 | 0.588422151 | RNF8 | 1 | 1 | 1 |
| 0 | 0 | 8 | 0.16 | 2 | 0.14286 | 0.510166839 | 1 | 0.33939492 | ROS1 | 1 | 1 | 0.94492303 |
| 0 | 0 | 10 | 0.2 | 0 | 0 | 0.073759985 | 0.100854 | 0.33939492 | RPA1 | 0.50697 | 1 | 0.94492303 |
| 0 | 0 | 9 | 0.18 | 2 | 0.14286 | 0.471161887 | 1 | 0.341106332 | RPL26 | 1 | 1 | 0.94492303 |
| 0 | 0 | 3 | 0.06 | 0 | 0 | 1 | 1 | 1 | RPTOR | 1 | 1 | 1 |
| 0 | 0 | 8 | 0.16 | 2 | 0.14286 | 0.510166839 | 1 | 0.33939492 | RSPO2 | 1 | 1 | 0.94492303 |

| | | | | | | | | | | | | |
|---|---|---|---|---|---|---|---|---|---|---|---|---|
| 0 | 0 | 10 | 0.2 | 8 | 0.57143 | 0.003303893 | 0.0151297 | 0.105937061 | RSPO3 | 0.1730076 | 1 | 0.94492303 |
| 2 | 0.2 | 7 | 0.14 | 0 | 0 | 0.252625899 | 0.3315098 | 0.599529563 | RUNX1 | 0.9091937 | 1 | 1 |
| 0 | 0 | 5 | 0.1 | 1 | 0.07143 | 0.825954855 | 1 | 0.588422151 | RUNX1T1 | 1 | 1 | 1 |
| 1 | 0.1 | 14 | 0.28 | 3 | 0.21429 | 0.554981208 | 0.74248 | 0.434053099 | SBDS | 1 | 1 | 1 |
| 0 | 0 | 8 | 0.16 | 6 | 0.42857 | 0.021049833 | 0.0615412 | 0.192132251 | SDHA | 0.3148752 | 1 | 0.94492303 |
| 0 | 0 | 6 | 0.12 | 2 | 0.14286 | 0.724461627 | 1 | 0.587819901 | SDHAF2 | 1 | 1 | 1 |
| 0 | 0 | 8 | 0.16 | 0 | 0 | 0.164079918 | 0.1831159 | 0.587819901 | SDHB | 0.8012382 | 1 | 1 |
| 0 | 0 | 8 | 0.16 | 2 | 0.14286 | 0.510166839 | 1 | 0.33939492 | SDHC | 1 | 1 | 0.94492303 |
| 0 | 0 | 9 | 0.18 | 3 | 0.21429 | 0.356969362 | 0.7153727 | 0.199914832 | SDHD | 0.9763492 | 1 | 0.94492303 |
| 0 | 0 | 8 | 0.16 | 4 | 0.28571 | 0.210057837 | 0.4376874 | 0.199914832 | SERPINA1 | 0.8958294 | 1 | 0.94492303 |
| 2 | 0.2 | 23 | 0.46 | 2 | 0.14286 | 0.050377903 | 0.0356288 | 0.309253717 | SETBP1 | 0.4375048 | 1 | 0.94492303 |
| 0 | 0 | 5 | 0.1 | 1 | 0.07143 | 0.825954855 | 1 | 0.588422151 | SETD2 | 1 | 1 | 1 |
| 0 | 0 | 3 | 0.06 | 0 | 0 | 1 | 1 | 1 | SF1 | 1 | 1 | 1 |
| 1 | 0.1 | 4 | 0.08 | 2 | 0.14286 | 0.711545766 | 0.6043629 | 1 | SF3B1 | 1 | 1 | 1 |
| 1 | 0.1 | 6 | 0.12 | 0 | 0 | 0.487948944 | 0.3248367 | 1 | SH2B3 | 1 | 1 | 1 |
| 0 | 0 | 1 | 0.02 | 0 | 0 | 1 | 1 | 1 | SH2D1A | 1 | 1 | 1 |
| 0 | 0 | 6 | 0.12 | 1 | 0.07143 | 0.835169009 | 1 | 0.583378305 | SLC25A13 | 1 | 1 | 1 |
| 0 | 0 | 4 | 0.08 | 0 | 0 | 0.76152112 | 0.5681297 | 1 | SLC34A2 | 1 | 1 | 1 |
| 0 | 0 | 24 | 0.48 | 10 | 0.71429 | 0.000971787 | 0.1420079 | 0.001362443 | SLITRK6 | 0.1730076 | 1 | 0.248677373 |
| 0 | 0 | 2 | 0.04 | 1 | 0.07143 | 0.697642848 | 0.5295699 | 1 | SLX1A | 1 | 1 | 1 |
| 0 | 0 | 2 | 0.04 | 1 | 0.07143 | 0.697642848 | 0.5295699 | 1 | SLX1B | 1 | 1 | 1 |
| 0 | 0 | 3 | 0.06 | 3 | 0.21429 | 0.133673095 | 0.1128883 | 0.588422151 | SLX4 | 0.6919557 | 1 | 1 |
| 2 | 0.2 | 24 | 0.48 | 2 | 0.14286 | 0.037405813 | 0.0312365 | 0.301440266 | SMAD2 | 0.3968269 | 1 | 0.94492303 |
| 2 | 0.2 | 25 | 0.5 | 3 | 0.21429 | 0.062917015 | 0.0722331 | 0.186529271 | SMAD4 | 0.4797422 | 1 | 0.94492303 |
| 0 | 0 | 4 | 0.08 | 2 | 0.14286 | 0.551460797 | 0.6043629 | 0.588422151 | SMARCA4 | 1 | 1 | 1 |
| 0 | 0 | 11 | 0.22 | 2 | 0.14286 | 0.26168066 | 0.7153727 | 0.192779347 | SMARCB1 | 0.9091937 | 1 | 0.94492303 |
| 0 | 0 | 5 | 0.1 | 0 | 0 | 0.48654037 | 0.577127 | 1 | SMARCE1 | 1 | 1 | 1 |
| 0 | 0 | 11 | 0.22 | 1 | 0.07143 | 0.185567594 | 0.2740515 | 0.199914832 | SMC3 | 0.8624475 | 1 | 0.94492303 |
| 1 | 0.1 | 14 | 0.28 | 4 | 0.28571 | 0.614304199 | 1 | 0.436474262 | SMO | 1 | 1 | 1 |
| 0 | 0 | 5 | 0.1 | 2 | 0.14286 | 0.595089088 | 0.6418802 | 0.583378305 | SOCS1 | 1 | 1 | 1 |
| 0 | 0 | 1 | 0.02 | 0 | 0 | 1 | 1 | 1 | SOS1 | 1 | 1 | 1 |
| 0 | 0 | 3 | 0.06 | 4 | 0.28571 | 0.034904498 | 0.0357776 | 0.583378305 | SOX2 | 0.3785199 | 1 | 1 |
| 1 | 0.1 | 12 | 0.24 | 2 | 0.14286 | 0.625977068 | 0.71588 | 0.675901668 | SOX9 | 1 | 1 | 1 |
| 0 | 0 | 5 | 0.1 | 0 | 0 | 0.48654037 | 0.577127 | 1 | SPOP | 1 | 1 | 1 |
| 0 | 0 | 1 | 0.02 | 0 | 0 | 1 | 1 | 1 | SQSTM1 | 1 | 1 | 1 |
| 0 | 0 | 19 | 0.38 | 5 | 0.35714 | 0.042208669 | 1 | 0.025089129 | SRC | 0.4119566 | 1 | 0.618749838 |
| 0 | 0 | 11 | 0.22 | 2 | 0.14286 | 0.26168066 | 0.7153727 | 0.192779347 | SRSF2 | 0.9091937 | 1 | 0.94492303 |
| 1 | 0.1 | 23 | 0.46 | 2 | 0.14286 | 0.017458964 | 0.0356288 | 0.088306959 | SS18 | 0.3042848 | 1 | 0.94492303 |
| 0 | 0 | 1 | 0.02 | 0 | 0 | 1 | 1 | 1 | STAG1 | 1 | 1 | 1 |
| 0 | 0 | 6 | 0.12 | 0 | 0 | 0.342051159 | 0.3248367 | 0.588422151 | STAG2 | 0.9763492 | 1 | 1 |
| 0 | 0 | 12 | 0.24 | 4 | 0.28571 | 0.181669295 | 0.7355945 | 0.105743209 | STAT3 | 0.8607244 | 1 | 0.94492303 |
| 1 | 0.1 | 8 | 0.16 | 0 | 0 | 0.301201587 | 0.1831159 | 1 | STAT6 | 0.9763492 | 1 | 1 |
| 0 | 0 | 5 | 0.1 | 3 | 0.21429 | 0.26269735 | 0.357359 | 0.587819901 | STK11 | 0.9091937 | 1 | 1 |
| 0 | 0 | 9 | 0.18 | 0 | 0 | 0.122866881 | 0.1870181 | 0.346600866 | SUFU | 0.6919557 | 1 | 0.94492303 |
| 0 | 0 | 11 | 0.22 | 2 | 0.14286 | 0.26168066 | 0.7153727 | 0.192779347 | SUZ12 | 0.9091937 | 1 | 0.94492303 |
| 0 | 0 | 4 | 0.08 | 0 | 0 | 0.76152112 | 0.5681297 | 1 | SYK | 1 | 1 | 1 |
| 0 | 0 | 3 | 0.06 | 3 | 0.21429 | 0.133673095 | 0.1128883 | 0.588422151 | TAL1 | 0.6919557 | 1 | 1 |
| 0 | 0 | 3 | 0.06 | 0 | 0 | 1 | 1 | 1 | TAL2 | 1 | 1 | 1 |
| 0 | 0 | 4 | 0.08 | 1 | 0.07143 | 1 | 1 | 1 | TAZ | 1 | 1 | 1 |
| 0 | 0 | 2 | 0.04 | 2 | 0.14286 | 0.241977845 | 0.2056672 | 1 | TCF3 | 0.9091937 | 1 | 1 |
| 0 | 0 | 12 | 0.24 | 8 | 0.57143 | 0.005755209 | 0.0253683 | 0.05331369 | TCF7L1 | 0.1730076 | 1 | 0.847858997 |
| 0 | 0 | 10 | 0.2 | 0 | 0 | 0.073759985 | 0.100854 | 0.33939492 | TCF7L2 | 0.50697 | 1 | 0.94492303 |
| 0 | 0 | 14 | 0.28 | 8 | 0.57143 | 0.008351085 | 0.0584238 | 0.02789446 | TDG | 0.1730076 | 1 | 0.618749838 |
| 0 | 0 | 5 | 0.1 | 0 | 0 | 0.48654037 | 0.577127 | 1 | TENT5C | 1 | 1 | 1 |
| 0 | 0 | 5 | 0.1 | 5 | 0.35714 | 0.024244074 | 0.0330382 | 0.33939492 | TERC | 0.3148752 | 1 | 0.94492303 |
| 1 | 0.1 | 5 | 0.1 | 1 | 0.07143 | 1 | 1 | 1 | TERT | 1 | 1 | 1 |
| 0 | 0 | 3 | 0.06 | 0 | 0 | 1 | 1 | 1 | TET1 | 1 | 1 | 1 |
| 0 | 0 | 10 | 0.2 | 0 | 0 | 0.073759985 | 0.100854 | 0.33939492 | TET2 | 0.50697 | 1 | 0.94492303 |
| 0 | 0 | 2 | 0.04 | 2 | 0.14286 | 0.241977845 | 0.2056672 | 1 | TFE3 | 0.9091937 | 1 | 1 |
| 0 | 0 | 6 | 0.12 | 4 | 0.28571 | 0.12702587 | 0.2058665 | 0.33939492 | TLR4 | 0.6919557 | 1 | 0.94492303 |
| 0 | 0 | 6 | 0.12 | 1 | 0.07143 | 0.835169009 | 1 | 0.583378305 | TLX3 | 1 | 1 | 1 |
| 0 | 0 | 5 | 0.1 | 3 | 0.21429 | 0.26269735 | 0.357359 | 0.587819901 | TMEM127 | 0.9091937 | 1 | 1 |
| 2 | 0.2 | 8 | 0.16 | 3 | 0.21429 | 0.725508189 | 0.6933892 | 1 | TMPRSS2 | 1 | 1 | 1 |
| 0 | 0 | 6 | 0.12 | 4 | 0.28571 | 0.12702587 | 0.2058665 | 0.33939492 | TNFAIP3 | 0.6919557 | 1 | 0.94492303 |
| 0 | 0 | 2 | 0.04 | 1 | 0.07143 | 0.697642848 | 0.5295699 | 1 | TOPBP1 | 1 | 1 | 1 |
| 0 | 0 | 22 | 0.44 | 4 | 0.28571 | 0.014617963 | 0.3671901 | 0.011773622 | TP53 | 0.2743679 | 1 | 0.383035181 |
| 0 | 0 | 2 | 0.04 | 1 | 0.07143 | 0.697642848 | 0.5295699 | 1 | TP53BP1 | 1 | 1 | 1 |
| 0 | 0 | 10 | 0.2 | 3 | 0.21429 | 0.297408165 | 1 | 0.192779347 | TRAF3 | 0.9763492 | 1 | 0.94492303 |
| 0 | 0 | 1 | 0.02 | 1 | 0.07143 | 0.546464272 | 0.3923611 | 1 | TRAF7 | 1 | 1 | 1 |
| 0 | 0 | 6 | 0.12 | 0 | 0 | 0.342051159 | 0.3248367 | 0.588422151 | TRIM37 | 0.9763492 | 1 | 1 |
| 0 | 0 | 11 | 0.22 | 1 | 0.07143 | 0.185567594 | 0.2740515 | 0.199914832 | TSC1 | 0.8624475 | 1 | 0.94492303 |
| 0 | 0 | 8 | 0.16 | 4 | 0.28571 | 0.210057837 | 0.4376874 | 0.199914832 | TSC2 | 0.8958294 | 1 | 0.94492303 |
| 0 | 0 | 6 | 0.12 | 1 | 0.07143 | 0.835169009 | 1 | 0.583378305 | TSHR | 1 | 1 | 1 |
| 2 | 0.2 | 14 | 0.28 | 5 | 0.35714 | 0.737472001 | 0.7416156 | 0.714632084 | U2AF1 | 1 | 1 | 1 |
| 0 | 0 | 3 | 0.06 | 1 | 0.07143 | 1 | 1 | 1 | UBE2T | 1 | 1 | 1 |
| 0 | 0 | 7 | 0.14 | 5 | 0.35714 | 0.056632381 | 0.1148194 | 0.199914832 | UIMC1 | 0.453059 | 1 | 0.94492303 |
| 0 | 0 | 1 | 0.02 | 0 | 0 | 1 | 1 | 1 | UROD | 1 | 1 | 1 |
| 0 | 0 | 3 | 0.06 | 1 | 0.07143 | 1 | 1 | 1 | USP28 | 1 | 1 | 1 |
| 0 | 0 | 3 | 0.06 | 0 | 0 | 1 | 1 | 1 | USP8 | 1 | 1 | 1 |

| | | | | | | | | | | | | |
|---|---|---|---|---|---|---|---|---|---|---|---|---|
| 0 | 0 | 7 | 0.14 | 4 | 0.28571 | 0.164188161 | 0.2371794 | 0.341106332 | VEGFA | 0.8012382 | 1 | 0.94492303 |
| 0 | 0 | 13 | 0.26 | 6 | 0.42857 | 0.050792685 | 0.3207622 | 0.055597311 | VHL | 0.4375048 | 1 | 0.847858997 |
| 0 | 0 | 11 | 0.22 | 8 | 0.57143 | 0.00490288 | 0.0190206 | 0.055597311 | WAS | 0.1730076 | 1 | 0.847858997 |
| 0 | 0 | 23 | 0.46 | 3 | 0.21429 | 0.006656455 | 0.1293853 | 0.011773622 | WRN | 0.1730076 | 1 | 0.383035181 |
| 0 | 0 | 7 | 0.14 | 1 | 0.07143 | 0.62851523 | 0.6732942 | 0.587819901 | WT1 | 1 | 1 | 1 |
| 0 | 0 | 6 | 0.12 | 0 | 0 | 0.342051159 | 0.3248367 | 0.588422151 | XPA | 0.9763492 | 1 | 1 |
| 0 | 0 | 3 | 0.06 | 1 | 0.07143 | 1 | 1 | 1 | XPC | 1 | 1 | 1 |
| 0 | 0 | 3 | 0.06 | 0 | 0 | 1 | 1 | 1 | XPO1 | 1 | 1 | 1 |
| 0 | 0 | 9 | 0.18 | 6 | 0.42857 | 0.026177212 | 0.0749201 | 0.197470798 | XRCC1 | 0.3148752 | 1 | 0.94492303 |
| 0 | 0 | 8 | 0.16 | 4 | 0.28571 | 0.210057837 | 0.4376874 | 0.199914832 | XRCC2 | 0.8958294 | 1 | 0.94492303 |
| 0 | 0 | 8 | 0.16 | 2 | 0.14286 | 0.510166839 | 1 | 0.33939492 | XRCC3 | 1 | 1 | 0.94492303 |
| 0 | 0 | 1 | 0.02 | 3 | 0.21429 | 0.030267002 | 0.0302199 | 1 | XRCC4 | 0.3434953 | 1 | 1 |
| 0 | 0 | 4 | 0.08 | 1 | 0.07143 | 1 | 1 | 1 | XRCC5 | 1 | 1 | 1 |
| 0 | 0 | 9 | 0.18 | 3 | 0.21429 | 0.356969362 | 0.7153727 | 0.199914832 | XRCC6 | 0.9763492 | 1 | 0.94492303 |
| 0 | 0 | 3 | 0.06 | 3 | 0.21429 | 0.133673095 | 0.1128883 | 0.588422151 | YAP1 | 0.6919557 | 1 | 1 |
| 0 | 0 | 25 | 0.5 | 4 | 0.28571 | 0.005449123 | 0.2262717 | 0.005095848 | ZNF217 | 0.1730076 | 1 | 0.248677373 |
| 0 | 0 | 2 | 0.04 | 0 | 0 | 1 | 1 | 1 | ZNF708 | 1 | 1 | 1 |
| 0 | 0 | 7 | 0.14 | 1 | 0.07143 | 0.62851523 | 0.6732942 | 0.587819901 | ZNRF3 | 1 | 1 | 1 |
| 0 | 0 | 2 | 0.04 | 0 | 0 | 1 | 1 | 1 | ZRSR2 | 1 | 1 | 1 |

**Supplementary Table 4**: Orion multiplexed imaging primary antibody panel

| EX Laser | EM CWL | Exposure (ms) | Fluorophore | Target | Purpose | Clone | Antibody RRID | Concentration |
|---|---|---|---|---|---|---|---|---|
| 405 | 452 | 50 | Hoechst 33342 | DNA | Nuclear localization | N/A | | 10 µg/mL |
| 445 | 485 | 100 | N/A | Autofluoroscence | Tissue architecture | N/A | | |
| 470 | 513 | 100 | ArgoFluor 515 | CD31 | Endothelium | EPR3094 | RareCyte Cat# 52-1005-501, RRID:AB_2922783 | 2 µg/mL |
| | 560 | 100 | ArgoFluor 555L | CD45 | Leukocyte | D9M8I | RareCyte Cat# AFAB-010-B, RRID:AB_2922785 | 1 µg/mL |
| 520 | 540 | 200 | ArgoFluor 535 | CD68 | Macrophage | KP1 + C68/684 | RareCyte Cat# 52-1008-501, RRID:AB_2922780 | 0.5 µg/mL |
| 555 | 572 | 200 | ArgoFluor 572 | CD4 | T Helper | NIUG0 | RareCyte Cat# 52-1002-501, RRID:AB_2922787 | 2 µg/mL |
| | 586 | 200 | ArgoFluor 584 | FOXP3 | T Regulatory | 236A/E7 | RareCyte Cat# AFAB-018-B, RRID:AB_2922788 | 2 µg/mL |
| | 606 | 200 | ArgoFluor 602 | CD8a | T Cytotoxic | AMC908 | RareCyte Cat# 52-1003-601, RRID:AB_2922789 | 0.5 µg/mL |
| | 630 | 200 | ArgoFluor 624 | CD45RO | T Memory | UCHL1 | RareCyte Cat# 52-1007-601, RRID:AB_2922790 | 2 µg/mL |
| | 689 | 200 | ArgoFluor 660L | CD20 | B cell | L26 | RareCyte Cat# 52-1004-601, RRID:AB_2922791 | 1 µg/mL |
| 640 | 662 | 500 | ArgoFluor 662 | PD-L1 | Immune Checkpoint | E1L3N | RareCyte Cat# 52-1018-601, RRID:AB_2922792 | 2 µg/mL |
| | 686 | 500 | ArgoFluor 686 | CD3e | T cell | D7A6E | RareCyte Cat# 52-1001-601, RRID:AB_2922793 | 4 µg/mL |
| | 706 | 500 | ArgoFluor 706 | CD163 | Macrophage | EPR14643 | RareCyte Cat# AFAB-014-B, RRID:AB_2922794 | 1 µg/mL |
| | 732 | 500 | ArgoFluor 730 | E-Cadherin | Epithelium | 4A2 | RareCyte Cat# 52-1010-701, RRID:AB_2922795 | 2 µg/mL |
| 730 | 768 | 500 | ArgoFluor 760 | PD-1 | Immune Checkpoint | EPR4877(2) | RareCyte Cat# 52-1017-701, RRID:AB_2922797 | 1 µg/mL |
| | 798 | 500 | ArgoFluor 795 | Ki67 | Proliferation | D3B5 | RareCyte Cat# AFAB-021-B, RRID:AB_2922798 | 2 µg/mL |
| | 847 | 500 | ArgoFluor 845 | Pan-CK (DIG) | Epithelium | AE1/AE3/C11 | RareCyte Cat# 52-1015-801, RRID:AB_2922799 | 5 µg/mL |
| | 877 | 500 | ArgoFluor 875 | SMA (BIO) | Smooth Muscle | 1A4 | RareCyte Cat# 52-1023-111, RRID:AB_2922801 | 2 µg/mL |

**Supplementary Table 5**: 2D CyCIF antibody panel

| Cycle | Channel | Target | Label | Vendor(s) | Catalog Number(s) | Clone Name | Clonality |
|---|---|---|---|---|---|---|---|
| 1 | 1 | | | | | | |
| 1 | 2 | Donkey anti-C | Alexa Fluor 488 | Invitrogen | A-11055 | | polyclonal |
| 1 | 3 | Rabbit IgG | Alexa Fluor 555 | Thermo Fisher Scientific | A-31572 | | polyclonal |
| 1 | 4 | Mouse IgG | Alexa Fluor 647 | Thermo Fisher Scientific | A-21237 | | polyclonal |
| 2 | 1 | | | | | | |
| 2 | 2 | TROP2 | | R&D Systems | AF650 | | polyclonal |
| 2 | 3 | Donkey anti-C | Alexa Fluor 488 | Invitrogen | A-11055 | | polyclonal |
| 2 | 4 | SLAMF6 | | Cell Signaling Technology | 53988 | E3H1E | monoclonal |
| 2 | 5 | Granzyme B | | Dako | M7235 | GrB-7 | monoclonal |
| 3 | 1 | | | | | | |
| 3 | 2 | Cytokeratin (p | Alexa Fluor 488 | Thermo Fisher Scientific (eBioscience) | 53-9003-82 | AE1/AE3 | monoclonal |
| 3 | 3 | Alpha-actin-2 | eFluor 570 | Thermo Fisher Scientific | 41-9760-82 | 1A4 | monoclonal |
| 3 | 4 | CD45 | Alexa Fluor 647 | BioLegend | 304020, 3040 | HI30 | monoclonal |
| 4 | 1 | | | | | | |
| 4 | 2 | CD45RO | Alexa Fluor 488 | BioLegend | 304212 | UCHL1 | monoclonal |
| 4 | 3 | CD3d | Alexa Fluor 555 | Abcam | ab208514 | EP4426 | monoclonal |
| 4 | 4 | PD-L1 | Alexa Fluor 647 | Cell Signaling Technology | 15005 | E1L3N | monoclonal |
| 5 | 1 | | | | | | |
| 5 | 2 | CD4 | Alexa Fluor 488 | abcam | ab196372 | EPR6855 | monoclonal |
| 5 | 3 | FOXP3 | eFluor 570 | eBioscience | 41-4777-80, 4 | 236A/E7 | monoclonal |
| 5 | 4 | PD-1 | Alexa Fluor 647 | Abcam | ab201825 | EPR4877(2) | monoclonal |
| 6 | 1 | | | | | | |
| 6 | 2 | TCF1/TCF7 | Alexa Fluor 488 | Cell Signaling Technology | 6444 | C63D9 | monoclonal |
| 6 | 3 | LAG3 | Alexa Fluor 555 | Cell Signaling Technology | 56141BC | D2G4O™ | monoclonal |
| 6 | 4 | CTLA4 | Alexa Fluor 647 | Abcam | ab270127 | CAL49 | monoclonal |
| 7 | 1 | | | | | | |
| 7 | 2 | CD20 | Alexa Fluor 488 | eBioscience | 53-0202-80, 5 | L26 | monoclonal |
| 7 | 3 | CD68 | Alexa Fluor 555 | Cell Signaling Technology | 23308S | D4B9C | monoclonal |
| 7 | 4 | CD8a | eFluor 660 | eBioscience | 50-0008-82 | AMC908 | monoclonal |
| 8 | 1 | | | | | | |
| 8 | 2 | TIM3 | Alexa Fluor 488 | Cell Signaling Technology | 54669 | D5D5R | monoclonal |
| 8 | 3 | GITR | Alexa Fluor 555 | Cell Signaling Technology | 98430BC | D9I9D | monoclonal |
| 8 | 4 | CD134 | Alexa Fluor 647 | BioLegend | 350018 | Ber-ACT35 | monoclonal |
| 9 | 1 | | | | | | |
| 9 | 2 | NT5E/CD73 | Alexa Fluor 488 | Cell Signaling Technology | 14627BC | D7F9A | monoclonal |
| 9 | 3 | CD11c | Alexa Fluor 555 | Cell Signaling Technology | 77882BC | D3V1E | monoclonal |
| 9 | 4 | CD103 | Alexa Fluor 647 | Abcam | ab225153 | EPR4166(2) | monoclonal |
| 10 | 1 | | | | | | |
| 10 | 2 | CD31 | Alexa Fluor 488 | Abcam | ab275989 | EPR3094 | monoclonal |
| 10 | 3 | CDX2 | Alexa Fluor 555 | Cell Signaling Technology | 84638 | D11D10 | monoclonal |
| 10 | 4 | CD39 | Alexa Fluor 647 | Abcam | ab312897 | EPR20627 | monoclonal |
| 10 | 5 | Ki-67 | Alexa Fluor 750 | R&D Systems | IC7617S | 1297A | monoclonal |
| 11 | 1 | | | | | | |
| 11 | 2 | HLA-DRB1 | Alexa Fluor 488 | Abcam | ab207376 | EPR6148 | monoclonal |
| 11 | 3 | H3-Trimethyl- | Alexa Fluor 555 | Cell Signaling Technology | 11960 | C42D8 | monoclonal |
| 11 | 4 | HLA-DPB1 | Alexa Fluor 647 | abcam | ab201347 | EPR11226 | monoclonal |
| 12 | 1 | | | | | | |
| 12 | 2 | CD57 | FITC | BD Biosciences | 561906 | NK-1 | monoclonal |
| 12 | 3 | ICOS | Alexa Fluor 555 | Cell Signaling Technology | 79403BC | D1K2T™ | monoclonal |
| 12 | 4 | NKG2A | Alexa Fluor 647 | Abcam | ab300745 | EPR23737-12 | monoclonal |
| 13 | 1 | | | | | | |
| 13 | 2 | CD15 | Alexa Fluor 488 | BioLegend | 301910 | HI98 | monoclonal |
| 13 | 3 | SOX-2 | Alexa Fluor 555 | Cell Signaling Technology | 5179S | D6D9 | monoclonal |
| 13 | 4 | Histone H2A. | Alexa Fluor 647 | BioLegend | 613407, 6134 | 2F3 | monoclonal |
| 14 | 1 | | | | | | |
| 14 | 2 | EZH2 | Alexa Fluor 488 | Abcam | ab225362 | EPR9307(2) | monoclonal |
| 14 | 3 | Histone H3.1 | Alexa Fluor 555 | Cell Signaling Technology | 3475S | D2C8 | monoclonal |
| 14 | 4 | GNLY | Alexa Fluor 647 | Cell Signaling Technology | 64458S | E2T3D | monoclonal |
| 15 | 1 | | | | | | |
| 15 | 2 | Ki-67 | Alexa Fluor 488 | Cell Signaling Technology | 11882S | D3B5 | monoclonal |
| 15 | 3 | Histone H3.1 | phycoerythrin | Cell Signaling Technology | 40724S | C36B11 | monoclonal |
| 15 | 4 | CD206 | Alexa Fluor 647 | Santa Cruz Biotechnology | sc-376108 AF | D-1 | monoclonal |
| 16 | 1 | | | | | | |
| 16 | 2 | EGFR | Alexa Fluor 488 | Cell Signaling Technology | 5616S | D38B1 | monoclonal |
| 16 | 3 | VEGFR-2 | Alexa Fluor 555 | Cell Signaling Technology | 12872S | D5B1 | monoclonal |
| 16 | 4 | Stat1 (pY701) | Alexa Fluor 647 | Cell Signaling Technology | 8009S | 58D6 | monoclonal |

| Antibody Species | Isotype | Species reactivity | Antibody Name | RRID identifier |
|---|---|---|---|---|
| donkey | IgG | goat | Donkey anti-Goat IgG (H+L)-AF488 | AB_2534102 |
| donkey | IgG | rabbit | Donkey anti-Rabbit IgG-AF555 | AB_162543 |
| goat | IgG | mouse | Goat anti-Mouse IgG-AF647 | AB_2535806 |
| goat | IgG | human | TROP2 | AB_2205667 |
| donkey | IgG | goat | Donkey anti-Goat IgG (H+L)-AF488 | AB_2534102 |
| rabbit | IgG | human | SLAMF6 | |
| mouse | IgG2a, kappa | human | Granzyme B | AB_2114697 |
| mouse | IgG1 | human, mous | Cytokeratin (pan)-AF488 | AB_1834350 |
| mouse | IgG2aK | human, mous | Alpha-actin-2-eF570 | AB_2573631 |
| mouse | IgG1, kappa | human, Non-h | CD45-AF647 | AB_493034 |
| mouse | IgG2a, kappa | human, Non-h | CD45RO-AF488 | AB_528823 |
| rabbit | IgG | human | CD3d-AF555 | AB_2728789 |
| rabbit | IgG | human | PD-L1-AF647 | AB_2728832 |
| rabbit | IgG | human | CD4-AF488 | AB_2889191 |
| mouse | IgG1 | human, Non-h | FOXP3-eF570 | AB_2573609 |
| rabbit | IgG | human | PD-1-AF647 | AB_2728811 |
| rabbit | IgG | human, mous | TCF1/TCF7-AF488 | AB_2797627 |
| rabbit | IgG | human | LAG3-AF555 | AB_2798739 |
| rabbit | IgG | human | CTLA4-AF647 | AB_2943179 |
| mouse | IgG2a, kappa | human, Non-h | CD20-AF488 | AB_10734357 |
| rabbit | IgG | human | CD68-AF555 | AB_2922446 |
| mouse | IgG2a, kappa | human | CD8a-eF660 | AB_2574149 |
| rabbit | IgG | human | TIM3-AF488 | AB_2799468 |
| rabbit | IgG | human, Non-h | GITR-AF555 | AB_2799740 |
| mouse | IgG1, k | human | CD134-AF647 | AB_2571938 |
| rabbit | IgG | human, mous | NT5E/CD73-AF488 | AB_2716625 |
| rabbit | IgG | human | CD11c-AF555 | AB_3331656 |
| rabbit | IgG | human | CD103-AF647 | AB_2884945 |
| rabbit | IgG | human | CD31-AF488 | AB_2943189 |
| rabbit | IgG | human | CDX2-AF555 | AB_2943240 |
| rabbit | IgG | human, mous | CD39-AF647 | |
| rabbit | IgG | human | Ki-67-AF750 | AB_2889352 |
| rabbit | IgG | human | HLA-DRB1-AF488 | AB_2889845 |
| rabbit | IgG | human, mous | H3-Trimethyl-K4-AF555 | AB_2797779 |
| rabbit | IgG | human | HLA-BPB1-AD647 | AB_2861375 |
| mouse | IgM, k | human | CD57-FITC | AB_10897013 |
| rabbit | IgG | human | ICOS-AF555 | AB_2800142 |
| rabbit | IgG | human | NKG2A-AF647 | AB_2943211 |
| mouse | IgM | human | CD15-AF488 | AB_493257 |
| rabbit | not specified | human | Sox2-AF555 | AB_10828439 |
| mouse | IgG1, kappa | human, mous | Histone H2A.X-AF647 | AB_2114994 |
| rabbit | IgG | human | EZH2-AF488 | AB_2889274 |
| rabbit | IgG | human, mous | Histone H3.1 (pS10)-AF555 | AB_10694639 |
| rabbit | IgG | human | GNLY-AF647 | |
| rabbit | IgG | human, mous | Ki-67 / clone D3B5 | AB_2687824 |
| rabbit | IgG | human, mous | Histone H3.1 (K27me3)-PE | AB_2799182 |
| mouse | IgG1 kappa | human | CD206-AF647 | AB_2943303 |
| rabbit | IgG | human, mous | EGFR | AB_10691853 |
| rabbit | IgG | human, mous | VEGFR-2 | AB_2728828 |
| rabbit | IgG | human, mous | Stat1 (pY701)-AF647 | AB_10860764 |

**Supplementary Table 6**: 3D CyCIF antibody for LSP32455 tipMMR CRC datasets

| Cycle | Channel | Target | Label | Vendor(s) | Catalog Number(s) | Clone Name | Clonality | Antibody Species | Isotype | Species reactivity | Antibody Name | RRID identifier |
|---|---|---|---|---|---|---|---|---|---|---|---|---|
| 1 | 1 | Hoechst33342 | | | | | | | | | | |
| 1 | 2 | CD23 | Alexa Fluor 4 | Abcam | ab305270 | EPR3617 | monoclonal | rabbit | IgG | human | CD23-AF488 | AB_2943215 |
| 1 | 3 | CD21 | Alexa Fluor 5 | Abcam | ab282007 | EP3093 | monoclonal | rabbit | IgG | human | CD21-AF555 | |
| 1 | 4 | CD20 | eFluor 660 | eBioscience | 50-0202-80, 5 | L26 | monoclonal | mouse | IgG2a | human, Non-I | CD20-eF660 | AB_11151691 |
| 1 | 5 | Ki-67 | Alexa Fluor 7 | R&D Systems | IC7617S | 1297A | monoclonal | rabbit | IgG | human | Ki-67-AF750 | AB_2889352 |
| 2 | 1 | Hoechst33342 | | | | | | | | | | |
| 2 | 2 | Claudin-7 | Alexa Fluor 4 | Abcam | ab309661 | EPR18073 | monoclonal | rabbit | IgG | human | Claudin-7-AF488 | |
| 2 | 3 | E-cadherin | Alexa Fluor 5 | Cell Signaling | 4295 | 24E10 | monoclonal | rabbit | IgG | human | E-cadherin-A | AB_2728822 |
| 2 | 4 | CD31 | Alexa Fluor 6 | Abcam | ab218582 | EPR3094 | monoclonal | rabbit | IgG | human | CD31-AF647 | AB_2857973 |
| 2 | 5 | Cytokeratin (p | Alexa Fluor 7 | Novus Biolog | NBP2-33200/ | AE-1/AE-3 | monoclonal | mouse | IgG1 kappa | human, mous | Cytokeratin (p | AB_2868569 |
| 3 | 1 | Hoechst33342 | | | | | | | | | | |
| 3 | 2 | CD4 | Alexa Fluor 4 | R&D Systems | FAB8165G | | polyclonal | goat | IgG | human | CD4-AF488 | AB_2728839 |
| 3 | 3 | FOXP3 | eFluor 570 | eBioscience | 41-4777-80, 4 | 236A/E7 | monoclonal | mouse | IgG1 | human, Non-I | FOXP3-eF57 | AB_2573609 |
| 3 | 4 | PD-1 | Alexa Fluor 6 | Abcam | ab201825 | EPR4877(2) | monoclonal | rabbit | IgG | human | PD-1-AF647 | AB_2728811 |
| 4 | 1 | Hoechst33342 | | | | | | | | | | |
| 4 | 2 | Connexin43 | Alexa Fluor 4 | Santa Cruz B | sc-271837 AF | F-7 | monoclonal | mouse | IgG2a kappa | human, mous | Connexin43-AF488 | |
| 4 | 3 | HLA-DPB1 | Alexa Fluor 5 | Abcam | ab300098 | EPR11226 | monoclonal | rabbit | IgG | human | HLA-DPB1-A | AB_2943208 |
| 4 | 4 | PD-L1 | Alexa Fluor 6 | Cell Signaling | 15005 | E1L3N | monoclonal | rabbit | IgG | human | PD-L1-AF647 | AB_2728832 |
| 5 | 1 | Hoechst33342 | | | | | | | | | | |
| 5 | 2 | LAG3 | Alexa Fluor 4 | Abcam | ab225277 | EPR4392(2) | monoclonal | rabbit | IgG | human | LAG3-AF488 | |
| 5 | 3 | GZMB | Alexa Fluor 5 | Cell Signaling | 29268 | D6E9W | monoclonal | rabbit | IgG | human, mous | GZMB-AF555 | |
| 5 | 4 | CD8a | eFluor 660 | eBioscience | 50-0008-82 | AMC908 | monoclonal | mouse | IgG2a, kappa | human | CD8a-eF660 | AB_2574149 |
| 6 | 1 | Hoechst33342 | | | | | | | | | | |
| 6 | 2 | pMLC2 | Alexa Fluor 4 | Cell Signaling | 35145BC | E2J8F | monoclonal | rabbit | IgG | human, mous | pMLC2-AF488 | |
| 6 | 3 | ICOS | Alexa Fluor 5 | Cell Signaling | 79403BC | D1K2T™ | monoclonal | rabbit | IgG | human | ICOS-AF555 | AB_2800142 |
| 6 | 4 | CD134 | Alexa Fluor 6 | BioLegend | 350018 | Ber-ACT35 | monoclonal | mouse | IgG1, k | human | CD134-AF64 | AB_2571938 |
| 7 | 1 | Hoechst33342 | | | | | | | | | | |
| 7 | 2 | CD163 | Alexa Fluor 4 | Abcam | ab218293 | EPR14643-3 | monoclonal | rabbit | IgG | human | CD163-AF48 | AB_2889155 |
| 7 | 3 | Tubulin beta ( | Alexa Fluor 5 | Cell Signaling | 2116S | 9F3 | monoclonal | rabbit | IgG | human, mous | Tubulin beta ( | AB_10695881 |
| 7 | 4 | CD103 | Alexa Fluor 6 | Abcam | ab225153 | EPR4166(2) | monoclonal | rabbit | IgG | human | CD103-AF64 | AB_2884945 |
| 8 | 1 | Hoechst33342 | | | | | | | | | | |
| 8 | 2 | AF488 | | | | | | | | | | |
| 8 | 3 | AF555 | | | | | | | | | | |
| 8 | 4 | AF647 | | | | | | | | | | |
| 9 | 1 | Hoechst33342 | | | | | | | | | | |
| 9 | 2 | TCF1/TCF7 | Alexa Fluor 4 | Cell Signaling | 6444 | C63D9 | monoclonal | rabbit | IgG | human, mous | TCF1/TCF7-A | AB_2797627 |
| 9 | 3 | CD11c | Alexa Fluor 5 | Cell Signaling | 77882BC | D3V1E | monoclonal | rabbit | IgG | human | CD11c-AF55 | AB_3331656 |
| 9 | 4 | CD39 | Alexa Fluor 6 | Abcam | ab312897 | EPR20627 | monoclonal | rabbit | IgG | human, mous | CD39-AF647 | |
| 10 | 1 | Hoechst33342 | | | | | | | | | | |
| 10 | 2 | MX1 | Alexa Fluor 4 | Cell Signaling | 7937BC | D3W7I | monoclonal | rabbit | IgG | human | MX1-AF488 | AB_2799122 |
| 10 | 3 | CD3E | Alexa Fluor 5 | Cell Signaling | 57869BC | D7A6E | monoclonal | rabbit | IgG | human | CD3E-AF555 | AB_3668891 |
| 10 | 4 | Transcription | Alexa Fluor 6 | Cell Signaling | 6709S | C63D9 | monoclonal | rabbit | IgG | human, mous | TCF1/TCF7-A | AB_2797631 |
| 10 | 5 | Beta-catenin | Alexa Fluor 7 | Cell Signaling | 37106BC | D10A8 | monoclonal | rabbit | IgG | human, mous | Beta-catenin-AF750 | |
| 11 | 1 | Hoechst33342 | | | | | | | | | | |
| 11 | 2 | TCF1/TCF7 | Alexa Fluor 4 | Cell Signaling | 6444 | C63D9 | monoclonal | rabbit | IgG | human, mous | TCF1/TCF7-A | AB_2797627 |
| 11 | 3 | TIGIT | Alexa Fluor 5 | Cell Signaling | 69779BC | E5Y1W | monoclonal | rabbit | IgG | human | TIGIT-AF555 | |
| 11 | 4 | CTLA4 | Alexa Fluor 6 | Abcam | ab270127 | CAL49 | monoclonal | rabbit | IgG | human | CTLA4-AF64 | AB_2943179 |
| 12 | 1 | Hoechst33342 | | | | | | | | | | |
| 12 | 2 | TIM3 | Alexa Fluor 4 | Cell Signaling | 54669 | D5D5R | monoclonal | rabbit | IgG | human | TIM3-AF488 | AB_2799468 |
| 12 | 3 | Actin, cytopla | Alexa Fluor 5 | Cell Signaling | 8046S | 13E5 | monoclonal | rabbit | IgG | human, mous | Actin, cytopla | AB_11179208 |
| 12 | 4 | VCAM1 | Alexa Fluor 6 | Biotechne | FAB8091R-1 | 1027433 | monoclonal | mouse | IgG | human | VCAM1-AF647 | |
| 13 | 1 | Hoechst33342 | | | | | | | | | | |
| 13 | 2 | TCF1/TCF7 | Alexa Fluor 4 | Cell Signaling | 6444 | C63D9 | monoclonal | rabbit | IgG | human, mous | TCF1/TCF7-A | AB_2797627 |
| 13 | 3 | CD68 | Alexa Fluor 5 | Cell Signaling | 23308S | D4B9C | monoclonal | rabbit | IgG | human | CD68-AF555 | AB_2922446 |
| 13 | 4 | ICAM1 | Alexa Fluor 6 | Abcam | ab214944 | EPR4776 | monoclonal | rabbit | IgG | human | ICAM1-AF647 | |
